# Atypical plume-like events drive glutamate accumulation in metabolic stress conditions

**DOI:** 10.1101/2025.02.05.636645

**Authors:** Tim Ziebarth, Nils Pape, Joel S.E. Nelson, Fleur I.M. van Alphen, Manu Kalia, Hil G.E. Meijer, Christine R. Rose, Andreas Reiner

**Affiliations:** Department of Biology and Biotechnology, Ruhr University Bochum, Universitätsstrasse 150, 44801 Bochum, Germany; Institute of Neurobiology, Heinrich Heine University Düsseldorf, Universitätsstrasse 1, 40225 Düsseldorf, Germany; Department of Applied Mathematics, University of Twente, Drienerlolaan 5, 7522 NB Enschede, Netherlands

**Keywords:** NMDA receptor, excitatory amino acid transporter (EAAT), ATP, epileptiform, excitotoxicity

## Abstract

Neural glutamate homeostasis plays a key role in health and disease. In ischemic conditions, such as stroke, this homeostasis is severely disrupted since energy depletion and ion imbalances lead to more glutamate release and less uptake. We here used the fluorescent glutamate sensor SF-iGluSnFR(A184V) to probe the effects of chemical ischemia on extracellular glutamate dynamics *in situ*, using organotypic slice cultures from mouse cortex. SF-iGluSnFR imaging reported spontaneous glutamate release events, which indicate synchronous network activity, similar to calcium signals detected with GCaMP6f. In addition, glutamate imaging revealed local, asynchronous release events, which were atypically large and long-lasting and showed plume-like characteristics. Under baseline conditions plumes occurred with low frequency, were independent of network activity, and persisted in the presence of TTX. Plume induction was strongly favored by blocking glutamate uptake with TFB-TBOA, whereas blocking ionotropic glutamate receptors (iGluRs) suppressed plumes. Upon inducing chemical ischemia plumes became more pronounced and overly abundant, which resulted in large-scale accumulation of extracellular glutamate. Similar plumes were recently also observed in models of cortical spreading depression and migraine. We therefore propose that plumes represent a more general phenomenon induced by glutamate uptake dysfunction, which may contribute to glutamate-related excitotoxicity in various neurodegenerative and neurological disorders.

## Introduction

Glutamate is key for transmitting signals across excitatory synapses. In the healthy brain, the extracellular glutamate concentrations are strictly regulated to provide the right dosage for activating synaptic glutamate receptors in a spatially and temporally defined manner [Rusakov & Stewart, 2021; Reiner & Levitz, 2018; Rose *et al*., 2018]. However, the underlying processes, namely the vesicular release of glutamate from neurons, the astrocytic uptake of glutamate through excitatory amino acid transporters (EAATs) [Danbolt *et al*., 2016; Rodríguez-Campuzano, 2021], the restoration of ion gradients, and glutamate metabolism [Andersen *et al*., 2021], require substantial amounts of energy [Attwell & Laughlin, 2001].

Many pathological conditions are thought to involve dysregulation or disruption of neurotransmitter homeostasis that results in increased levels of extracellular glutamate and synaptic spill-over [Rusakov & Stewart, 2021]. Increased glutamate concentrations can interfere with normal synaptic transmission and plasticity. In higher doses glutamate may cause overexcitation, disrupt cellular Ca^2+^ homeostasis, and trigger cell death mechanisms, which is commonly referred to as ‘excitotoxicity’ [Choi 2020; Belov Kirdajova *et al*., 2020; Armada-Moreira *et al*., 2020].

An extreme situation is encountered under ischemic conditions, such as stroke. The lack of blood and oxygen supply causes strong energy depletion, yet some aberrant neural activity persists in the penumbra, i.e. the tissue surrounding the infarct core [Dirnagl *et al*., 1999; Dreier & Reiffurth, 2015; Passlick 2021]. The consequences are prolonged and repetitive depolarizations, deterioration of ion gradients, uncontrolled neurotransmitter release, impaired neurotransmitter uptake, acidification, swelling and delayed cell death. These processes are thought to involve glutamate-mediated overactivation of ionotropic glutamate receptors (iGluRs). Indeed, suppression of AMPA and NMDA receptor activity, has been shown to be neuroprotective in experimental stroke models [Choi, 2020] and NMDA receptor activation is deemed particularly important for cell death and survival [Ge *et al*., 2020; Yu *et al*., 2023]. However, the extent to which glutamate contributes to the acute phase of metabolic stress, e.g. by promoting spreading depolarizations (SDs), remains a controversially discussed [Andrew *et al*, 2022].

Until recently, glutamate levels could only be deduced by rather indirect means, i.e. microdialysis or measurement of receptor currents [Wu *et al*., 2022]. The development of genetically encoded fluorescence-based sensors has enabled us to obtain more direct, spatially and temporally defined information on glutamate release and accumulation. Especially iGluSnFR (intensity-based glutamate-sensing fluorescent reporter) [Marvin *et al*., 2013; Marvin *et al*., 2018] variants are now widely used to investigate stimulus-evoked release and spread of glutamate in different physiological paradigms (e.g. [Armbruster *et al*., 2016; Helassa *et al*., 2018; Franke *et al*., 2017; Jensen *et al*., 2019; Barnes *et al*., 2020; Herde *et al*., 2020; Matthews *et al.,* 2022; Aggarwal *et al.,* 2023]), but also in disease models [Brymer *et al*., 2021], including models of cortical SD [Enger *et al*., 2015; Rakers *et al*., 2017; Parker *et al*., 2021], epilepsy [Diaz Verdugo *et al*., 2019; Shimoda *et al*., 2020], depression [McGirr *et al*., 2017], Alzheimer’s [Hefendehl *et al*., 2016; Zott *et al*., 2019; Brymer *et al*., 2023] and Huntington’s disease [Parsons *et al*., 2016; Koch *et al*., 2018; Dvorzhak *et al*., 2019].

Here we set out to study the extracellular glutamate dynamics during metabolic stress conditions using a defined *in situ* model, in which we induced brief chemical ischemia in organotypic slice cultures from mouse cortex. Previous work had shown that this model recapitulates key features of the ischemic penumbra, such as reversible ATP depletion from cells and an increase in intracellular Na^+^ and extracellular K^+^ concentrations [Lerchundi *et al*., 2019a; Eitelmann *et al*., 2022; Meyer *et al*., 2022; Pape & Rose, 2023]. In this study we measured neuronal activity and changes in the extracellular glutamate concentration to focused on how glutamate release and uptake are altered in energy-scarce conditions.

Imaging of SF-iGluSnFR(A184V) expressed on the surface of neurons showed different types of glutamate dynamics, namely synchronous network activity and atypical local glutamate events. The latter are reminiscent of recently described “plume-like” events [Parker *et al*., 2021]. Brief chemical ischemia resulted in a transient loss of synchronous network activity, yet a strong increase in the occurrence of plumes, which were the main driver of the observed extracellular glutamate accumulation. Plumes were favored by blocking glutamate uptake and they were suppressed by inhibiting iGluRs, which points to a feed-forward mechanism, in which elevated levels of glutamate trigger plume-like glutamate release events. The emergence and pathological role of plumes deserves further investigation, as they might also occur under less severe pathological conditions.

## Results

### Chemical ischemia in an organotypic slice model

To investigate the effects of metabolic stress on the extracellular glutamate dynamics, we prepared cortico-hippocampal slice cultures from mice and expressed the genetically encoded fluorescent glutamate sensor SF-iGluSnFR(A184V) [Marvin *et al*., 2018] under control of a synapsin promoter (hSyn1) using recombinant adeno-associated virus particles (rAAVs). After 14-21 days expression *in vitro*, sensor fluorescence was seen throughout all cortical layers (**Fig. 1A**). Healthy slices showed spontaneous, synchronized activity, as seen by spikes in fluorescence intensity (0.19 ± 0.11 Hz, mean ± s.d., n = 11 slices; **Fig. 1B**). This behavior is characteristic for slices of this age [Johnson & Buonomano, 2007; Okamoto *et al*., 2014] and was also observed by neuronal Ca^2+^ imaging using GCaMP6f (**Fig. 1E-H**, 0.16 ± 0.10 Hz, mean ± s.d., n = 8 slices).

**Figure 1.**
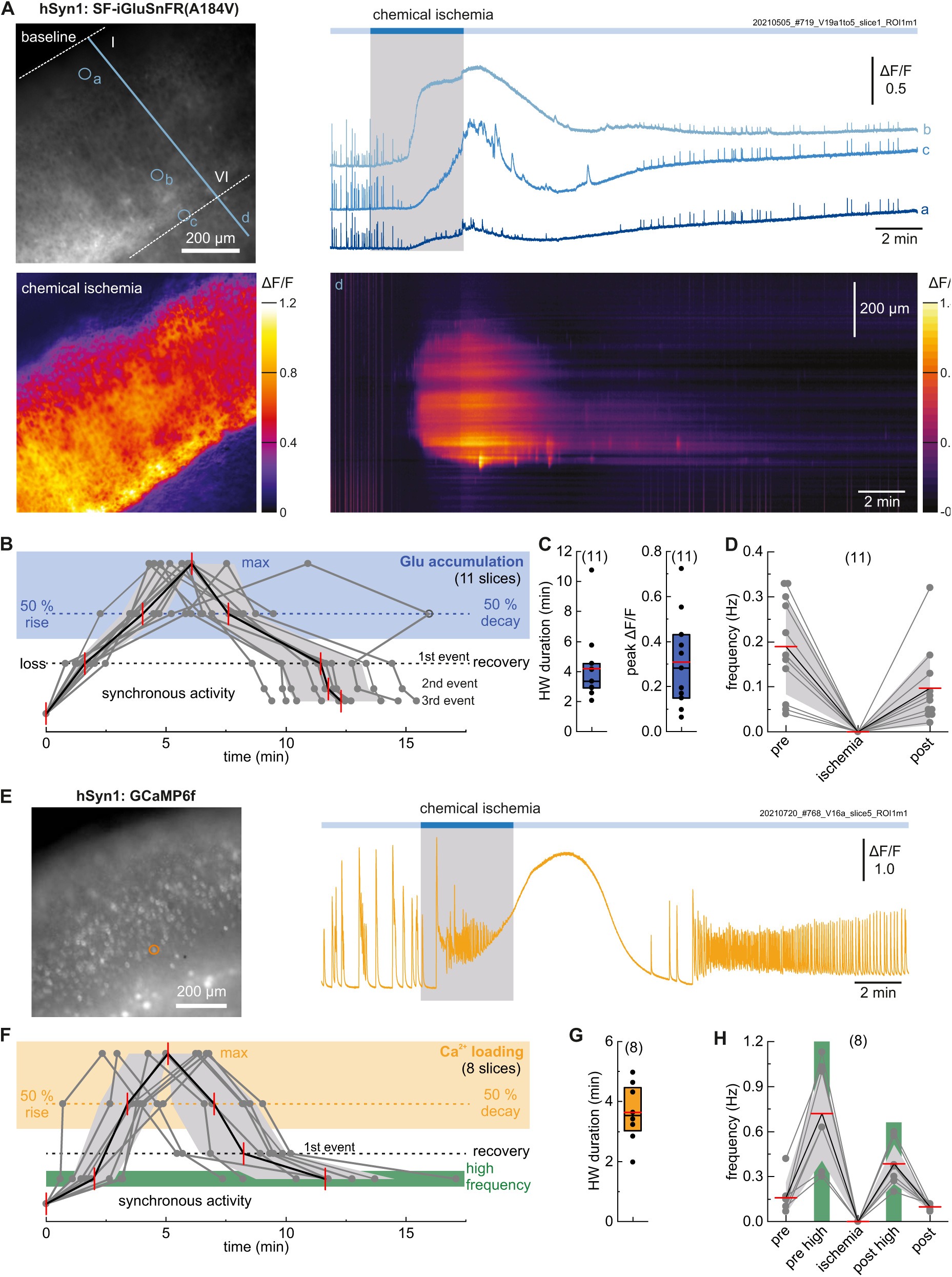
Optical probing of chemical ischemia in organotypic slice cultures from mouse cortex. **(A)** SF-iGluSnFR(A184V) widefield imaging of different cortical layers. Chemical ischemia (no glucose, 2 mM 2-deoxy-D-glucose and 5 mM sodium azide for 2-5 min) results in reversible loss of synchronous activity and large-scale glutamate accumulation. Right: Relative fluorescence changes (ΔF/F) for selected regions (a-c) and across the cortex (line d). **(B)** Timepoints of loss and recovery of synchronous activity (bottom) and glutamate accumulation (50% rise, maximum, and 50% decay times; top) for n = 11 slices. **(C)** Half-width duration of glutamate accumulation and peak ΔF/F of responding regions. **(D)** Frequency of synchronous Glu events before, during, and after ischemia (50% decay time + 10 min). **(E)** GCaMP6f imaging shows increased synchronous activity before and after chemical ischemia, and neuronal Ca^2+^ loading during ischemia. **(F)** Timepoints of Ca^2+^ loading (top), high frequency periods, and recovery of synchronous activity (bottom) for n = 8 slices. **(G)** Half-width duration of Ca^2+^ loading. **(H)** Frequency of synchronous Ca^2+^ activity. Individual data points are shown in grey, mean values in red and s.d. as shaded areas. Boxes show medians and 25-75% percentiles with red bars indicating the mean. For details see Methods.

To mimic energy restriction as it might occur in the ischemic penumbra, we induced transient chemical ischemia by removing glucose and adding 2 mM 2-deoxy-D-glucose and 5 mM sodium azide to block glycolysis and mitochondrial electron transport, respectively [Gerkau *et al*., 2018] (**Fig. 1A**). After 2-5 min, we switched back to normal Ringer’s solution. Inducing chemical ischemia for 2 min caused a transient decrease of the intracellular ATP concentration, as seen by a 18.8 ± 2.6% fluorescence ratio decrease of the FRET-based ATP sensor, ATeam1.03^YEMK^ (“ATeam”) [Imamura *et al*., 2009; Lerchundi *et al*., 2020; Pape & Rose, 2023] compared to baseline levels (mean ± s.d., n = 39 cells in 4 slices; **Fig. S1**). Inducing chemical ischemia for 5 min strengthened the drop in the ATeam fluorescence ratio to 54.1 ± 8.0% (n = 23 cells in 3 slices). Afterwards, the neuronal ATP levels recovered and the ATeam ratio returned to the initial baseline within 10 min after washout (641.6 ± 79.3 s, mean ± s.d.) after inducing ischemia for 2 min, and to 92.4 ± 4.4% of the initial ATP level after inducing ischemia for 5 min.

Energy restriction had pronounced effects on the SF-iGluSnFR signals. Within 2 min after inducing chemical ischemia, the fluorescence signals reporting synchronous activity had fully ceased in all regions. Still, somewhat later, a sustained increase in SF-iGluSnFR fluorescence was seen, which started locally and eventually spread across extended areas of the slice (**Fig. 1A,B** and **Movie S1**). We attribute this signal increase to an overall increase in extracellular glutamate concentration, since SF-iGluSnFR does not respond to any other neurotransmitters [Marvin *et al*., 2013]. In addition, control experiments confirmed that i) SF-iGluSnFR is minimally affected by the applied azide and that ii) acidification of the extracellular space, which often accompanies ischemia [Tóth *et al*., 2020], would decrease rather than increase the baseline fluorescence and signal amplitude (**Fig. S2**).

The half-maximal signal rise in glutamate accumulation (50% rise) was observed 4.03 ± 0.92 min after inducing ischemia (mean ± s.d., n = 11 slices). The fluorescence increase started and was most pronounced in more central regions of the slices. At the peak of the accumulation, the relative fluorescence increase of the responding area was ΔF/F = 0.31 ± 0.19 (mean ± s.d., n = 11 slices; **Fig. 1C**). In some cases, the fluorescence intensity during accumulation exceeded the signal changes associated with synchronous activity. In some areas the intensity was even as high as during a control application of a highly saturating glutamate concentration (10 mM) (**Fig. S3**). However, regions that responded most strongly to bath-applied glutamate, showed no or little glutamate accumulation during ischemia (**Fig. S3B**). This indicates that the observed glutamate accumulations might depend on the degree of bath coupling (see **Fig. S4-6** and **Supplementary Methods** for simulations), i.e. deeper or more dense regions of the slice, which have less bath exchange, may show more glutamate accumulation during ischemia.

Upon reperfusion with normal Ringer’s solution the glutamate accumulation decayed within 4.21 ± 2.67 min (time to half-maximal signal decay; mean ± s.d., n = 11 slices). Synchronous events reappeared 7.28 ± 1.45 min after washout (mean ± s.d., n = 11 slices), but their signal amplitudes were consistently decreased (2.7-fold, **Fig. S7B**). In contrast, the SF-iGluSnFR baseline intensity showed a run-up compared to the beginning of the recording, which was also seen in 40 min control recordings without inducing chemical ischemia (**Fig. S8**).

Besides extracellular glutamate accumulation, chemical ischemia resulted in a strong increase in neuronal GCaMP6f fluorescence, which indicates extensive Ca^2+^ loading (see also **Movie S2**). The onset of neuronal Ca^2+^ loading (3.38 ± 1.17 min to half-maximal signal rise, mean ± s.d., n = 8 slices) and duration (**Fig. 1F,G**) were comparable to extracellular glutamate accumulation (**Fig. 1B,C**). After reperfusion with standard Ringer’s solution spontaneous activity returned (first events 4.94 ± 0.92 min after washout, mean ± s.d., n = 8 slices) albeit with slower decay times (**Fig. S7C**). GCaMP6f expression under control of the interneuron-specific mDlx enhancer [Dimidschstein *et al*., 2016] showed similar behavior for this interneuron sub-population only (**Fig. S9**).

A notable difference between the Ca^2+^ and Glu dynamics was seen in the early phase of ischemia. Here Ca^2+^ imaging showed a high-frequency period, i.e. increased spontaneous activity, 1.96 ± 0.88 min after inducing ischemia (mean ± s.d., n = 8 slices), which coincided with the beginning of intracellular Ca^2+^ loading (**Fig. 1E,F**). In contrast, spontaneous glutamate release events decreased in signal amplitude and became fully undetectable 2.57 ± 1.10 min before large scale glutamate accumulation set in (mean ± s.d., n = 11 slices, **Fig. S7A**). This suggests that in the early ischemic period the amount of glutamate released during synchronous activity decreased and eventually became undetectable, while intracellular Ca^2+^ showed continuing oscillatory activity. During recovery a second high-frequency Ca^2+^ period was seen (8.51 ± 2.26 min after washout, mean ± s.d., n = 8 slices), i.e. around the time when the first synchronous glutamate events reappeared (**Fig. 1B,F**). Autonomous, high-frequency neuronal activity is typically seen at intermediate depolarization levels, as also seen in our simulations (**Supporting Information**, **Fig. S5**, and [Kalia *et al*., 2021]).

Taken together, these results show that the applied chemical ischemia protocol causes transient ATP depletion and Ca^2+^ accumulation in neurons, transient disruption of synchronous network activity, and yet strong glutamate accumulation in the extracellular space.

### SF-iGluSnFR imaging shows local glutamate events under baseline conditions

As described above, SF-iGluSnFR widefield imaging showed synchronous network activity under baseline conditions. Synchronous activity spread over all cortical layers (**Fig. 2A**, average intensity projection) and the events had fast and rather uniform characteristics. Their kinetics were not fully resolved by our 20 fps standard imaging rate but sometimes showed clear bursting behavior (see also **Fig. S11** for 99 fps data). As characterized above, these synchronous glutamate release events showed similar frequencies as Ca^2+^ events (**Fig. 1D,H**), and appear to reflect spontaneous network activity, which is typical for organotypic slices at this age [e.g. Johnson & Buonomano, 2007; Okamoto *et al*., 2014].

**Figure 2.**
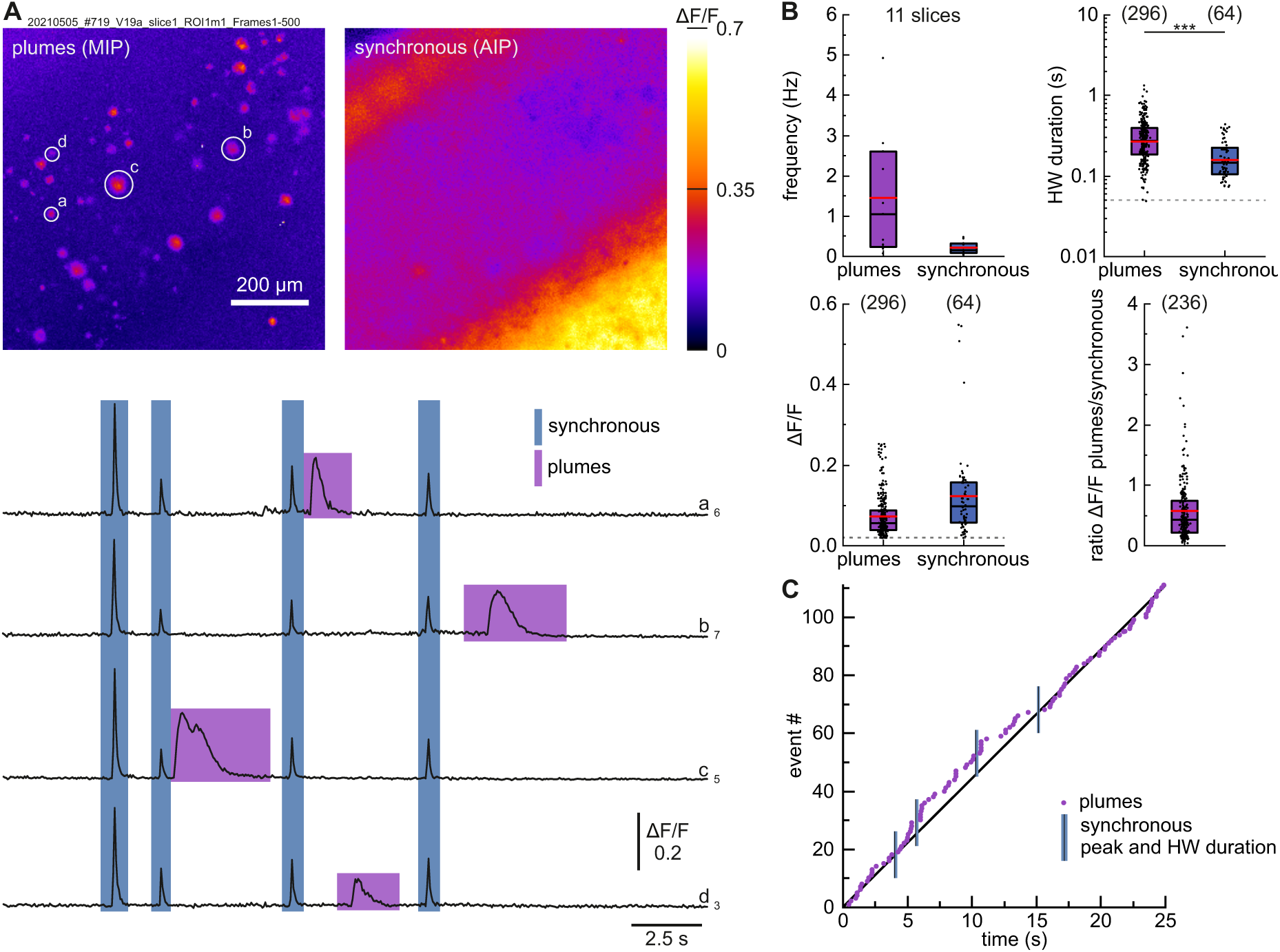
SF-iGluSnFR(A184V) imaging shows local glutamate events (“plumes”) which occur independently of synchronous network activity. **(A)** Left: Glutamate plumes as seen in a maximal intensity projection (MIP; 25 s, excluding frames with synchronous activity). Right: Fluorescence changes during synchronous activity (average intensity projection (AIP) of frames containing the event maxima). Bottom: ΔF/F traces of four selected regions (a-d). Synchronous events and local plumes can be clearly distinguished. Data were taken from the pre-ischemic baseline of the slice shown in Fig. 1A. **(B)** Frequencies of plumes and synchronous activity from n = 11 slices. Half-width duration and maximum ΔF/F values under baseline conditions from n = 10 slices. Most plumes show smaller ΔF/F values than synchronous activity in the same region. Numbers in parentheses give the number of analyzed events, dashed lines show the detection limits. Boxes show medians and 25-75% percentiles, red bars indicate the mean. For details see Methods. **(C)** Timepoints of plume occurrence and periods of synchronous activity during the 25 s imaging example.

Unexpectedly, the imaging data, revealed also another type of activity – local, asynchronous glutamate release events, which were seen in baseline conditions and far more prominently during ischemia and recovery. In the following we characterize these events in more detail.

Under baseline conditions (pre-ischemia), local asynchronous glutamate release events were clearly discernible from global synchronous activity (**Fig. 2**, **Fig. S10**, and **Movie S3**). They were limited to small regions and were quite heterogenous in size and duration (**Fig. 2A**). In order to collect these events, we calculated maximum intensity projections over 25 - 100 s (excluding frames which contained synchronous activity) and analyzed all identifiable events in the imaged region (see Methods for details). In the field-of-view (824 x 824 μm^2^ = 0.679 mm^2^), we captured local events with a median frequency of 1.04 Hz (n = 11 slices; **Fig. 2B** and **Fig. S12**), but the frequency varied strongly between slices (0.06 - 4.92 Hz). Smaller, less intense, or shorter events might have escaped our detection, in particular in slices with low SF-iGluSnFR expression levels (see Methods). This might also explain, why in some other slices under pre-ischemic conditions we did not detect these events at all (see Discussion).

The captured local events had a half-width duration of 267 ± 190 ms (mean ± s.d.; n = 296 events), i.e. most of these events lasted considerably longer than synchronous events (**Fig. 2B**). The average signal amplitude ΔF/F of local events was 0.07 ± 05 (mean ± s.d.; n = 296 events), which is much lower than typical ΔF/F values of synchronous activity in the same region (**Fig. 2B**). Still, for some local events the signal increase exceeded the signal change during synchronous activity. Based on their long-lasting but transient properties we refer to these local events as glutamate “plumes”, a term that was recently introduced by Parker *et al*. [Parker *et al*., 2021].

Analysis of individual movies showed that plumes occurred independently of synchronous activity (**Fig. 2C** and **Fig. S10B**), although the incidence of plumes sometimes appeared higher just before and after global, synchronous events.

### Local glutamate events (“plumes”) have unique properties

We next characterized the spatial and temporal characteristics of individual plumes that we had observed under baseline conditions (pre-ischemia). To determine plume sizes, we used the frame with the maximal plume ΔF/F to measure the full-width at half-maximum (FWHM) in x and y direction, which revealed an aspect ratio of ~1 (1.02 ± 0.20; mean ± s.d., n = 296), i.e. plumes had an overall roundish appearance (**Fig. 3A,B**). Elongated shapes were mostly obtained for small plumes, which might be due to their less precise detection. The mean FWHM size was 17.2 ± 6.4 μm (mean ± s.d., n = 296), with very large plumes having FWHM sizes of 38.6 μm or more. The lowest plume size was 6.44 μm, which reflects our detection limit (see Methods).

**Figure 3.**
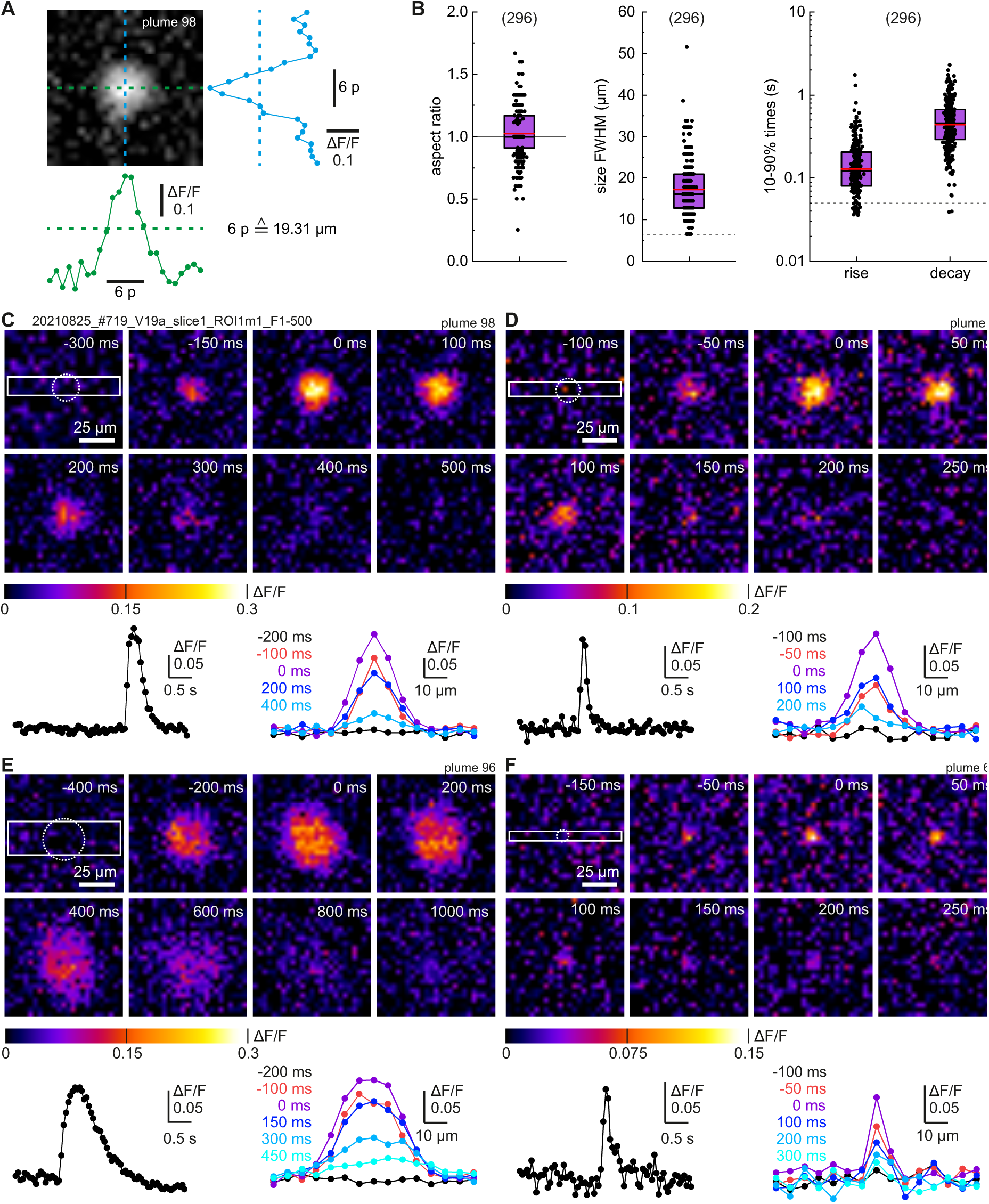
Spatial and temporal characteristics of individual plumes. **(A)** Example plume with ΔF/F cross-sections. **(B)** Aspect ratio and mean full-width at half maximum (FWHM) obtained from ΔF/F images at the peak maximum for n = 296 plumes from 11 slices pre-ischemia (see also **Fig. S12**). (Right) Rise and decay times (10-90%) obtained from ΔF/F traces of individual plumes. **(C-F)** Depiction of four individual plumes with different spatial and temporal characteristics. ΔF/F time traces (left) are based on the circled regions. Spatial profiles at different time points (right) are based on the boxed regions. Time zero reflects the plume maximum. For image processing see **Fig. S13**.

The rise times of individual plumes were rather fast (129 ± 160 ms; 10-90% signal change, mean ± s.d., n = 296; **Fig. 3B**) and close to the temporal detection limit of our recordings. The plume decay times were considerably slower (440 ± 361 ms; mean ± s.d., n = 296). A comparison of different plume parameters reveals a moderate correlation between plume size and duration (Spearman coefficient *r* = 0.56, 296 plumes, **Fig. S13**), but weaker correlations between plume size and plume intensity (*r* = 0.40), and between plume intensity and duration (*r* = 0.36). Despite this weak correlation, even some short plumes showed high signal changes.

Examples plumes with different characteristics are shown in **Fig. 3C-F**. Cross sections show that the highest ΔF/F amplitudes were found in the plume center. However, in large plumes the area with maximal signal changes extended over 10 μm or more (cf. **Fig. 3E**), which could either reflect homogenous glutamate distribution or SF-iGluSnFR saturation. Notably, both, signal increase and decay appeared spatially uniform, i.e. the maximum stayed in the plume center and the plumes did not visibly grow larger or shrink during their formation and disappearance.

The plume properties were rather uniform across different slices (**Fig. S12**) and their key characteristics did not significantly change during 40 min continuous control recordings without inducing ischemia (**Fig. S14**). Only the plume intensity ΔF/F decreased to 75%, which is similar to the signal loss seen for synchronous events (cf. **Fig. S8**), while the plume frequency decreased to 56% (n = 5 slices, **Fig. S14**).

To test whether plumes depend on action potential-driven release we added the sodium channel blocker tetrodotoxin (TTX). Perfusion of 0.2 μM TTX for 4 min resulted in a complete loss of global synchronous activity, but plumes were left largely unaffected (**Fig. 4A** and **Fig. S15**): Plumes were observed throughout the recording and their size, duration, and intensity (ΔF/F) remained unchanged during TTX application (**Fig. S15**). The decrease in plume frequency was comparable to control. After TTX washout, synchronous glutamate release events (and Ca^2+^ events) returned with increased signal amplitudes ΔF/F (**Fig. S15**), most likely due to upscaling [Ibata *et al*., 2008]. Plume size was slightly increased, but the other plume characteristics remained unchanged.

**Figure 4.**
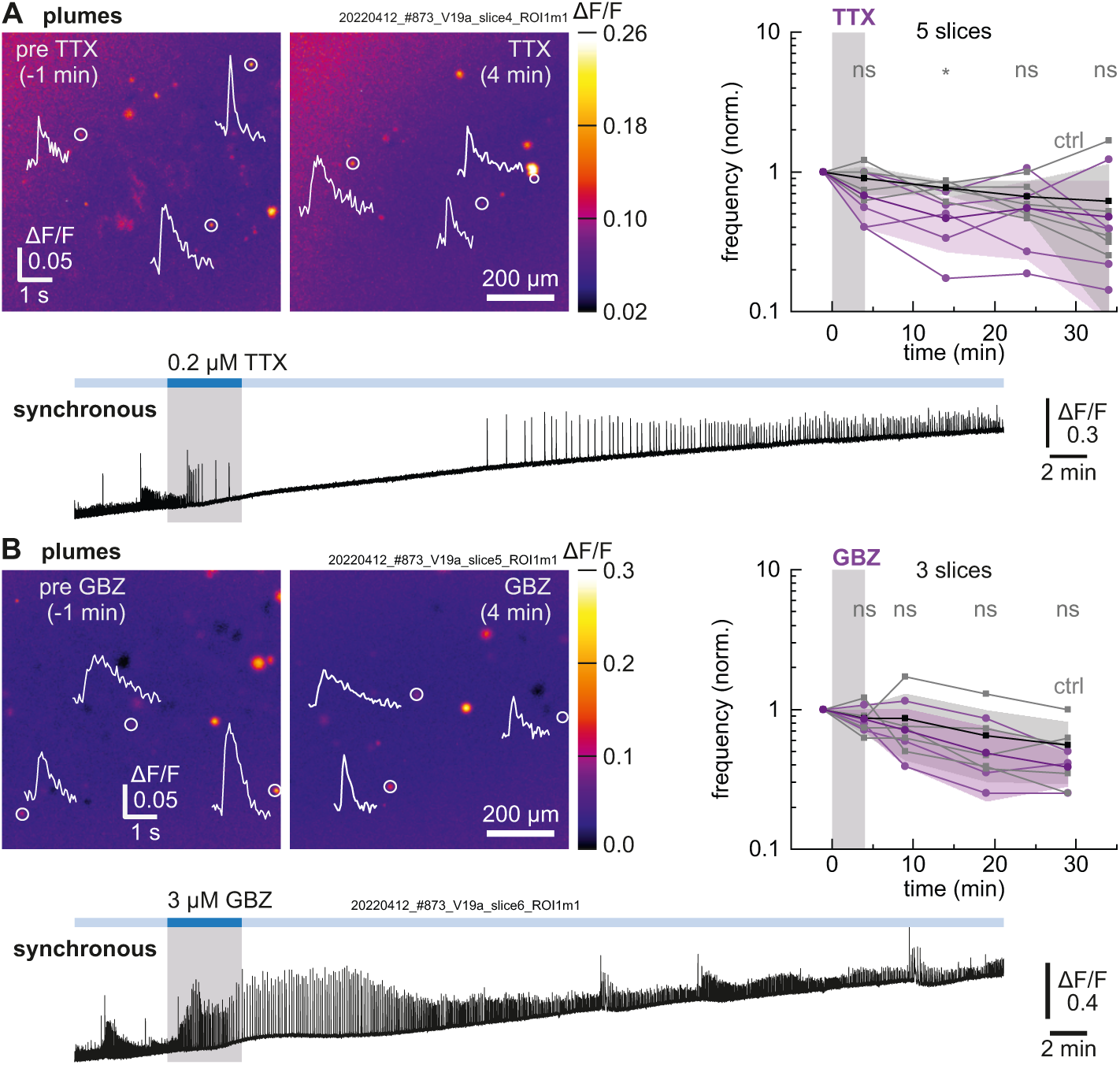
Plumes after blocking network activity with TTX or strengthening network activity with GBZ. **(A)** Addition of 0.2 μM TTX abolished synchronous activity (bottom), but plumes remained detectable (top, maximal intensity projections, 50 s each with traces of selected plumes). Right: Plume frequency at selected timepoints (n = 5 slices, purple) in comparison to control recordings without TTX (n = 5 slices, black). For details on TTX plumes see **Fig. S15**. **(B)** Addition of 3 μM GBZ strengthened synchronous activity (bottom), but the plume frequency (n = 3 slices, purple) did not change compared to control (n = 5 slices, black). For other parameters see **Fig. S16**. Individual data points are shown in light color, means in dark with s.d. as shaded areas. Mann-Whitney *U* tests were used for pair-wise comparison of individual time windows to control (cf. **Fig. S14**), ns not significant, \**p* < 0.05.

We next aimed to strengthen network activity by inhibiting GABA_A_ receptors with 3 μM gabazine (GBZ) [Bright & Smart, 2013]. Indeed, GBZ addition increased the signal amplitudes of both, synchronous glutamate release events and Ca^2+^ signals (**Fig. S16**), probably by reducing inhibitory inputs [Mohajerani & Cherubini, 2005]. However, also this manipulation did not change plume frequency or individual plume parameters compared to baseline (**Fig. 4B** and **Fig. S16**).

Next, we tested the role of glutamate uptake by adding TFB-TBOA, which blocks the glutamate transporters EAAT1, EAAT2 and to some extent also EAAT3 [Shimamoto *et al*., 2004]. Impaired glutamate uptake should prolong the synchronous glutamate transients and might affect network activity [Tsukada *et al*., 2005, Unichenko *et al*., 2015; Marvin *et al.,* 2013; Armbruster *et al.,* 2016; Barnes *et al.,* 2020], but it might also affect plumes directly. As expected, addition of 1 μM TFB-TBOA (**Fig. 5A**) had pronounced effects on synchronous activity resulting in prolonged glutamate events, more bursting behavior, and larger event amplitudes (n = 7 slices). The frequency of synchronous events temporarily decreased in most slices (**Fig. S17A**). The same effects were seen when monitoring Ca^2+^ influx (n = 5 slices; **Fig. S17B**). Notably, TFB-TBOA had a strong effect on the occurrence (induction) of plumes: TFB-TBOA resulted in a 6- to 73-fold increase in plume frequency at times at which synchronous event frequencies were decreased (**Fig. 5A, Fig. S17,** and **Movie S6**). Plume intensity (ΔF/F) or decay times (**Fig. 5A**) were not affected by TFB-TBOA, but a minor increase in rise time and a clear increase in plume size (1.3-fold increase in FWHM; **Fig. S17**) was observed.

**Figure 5.**
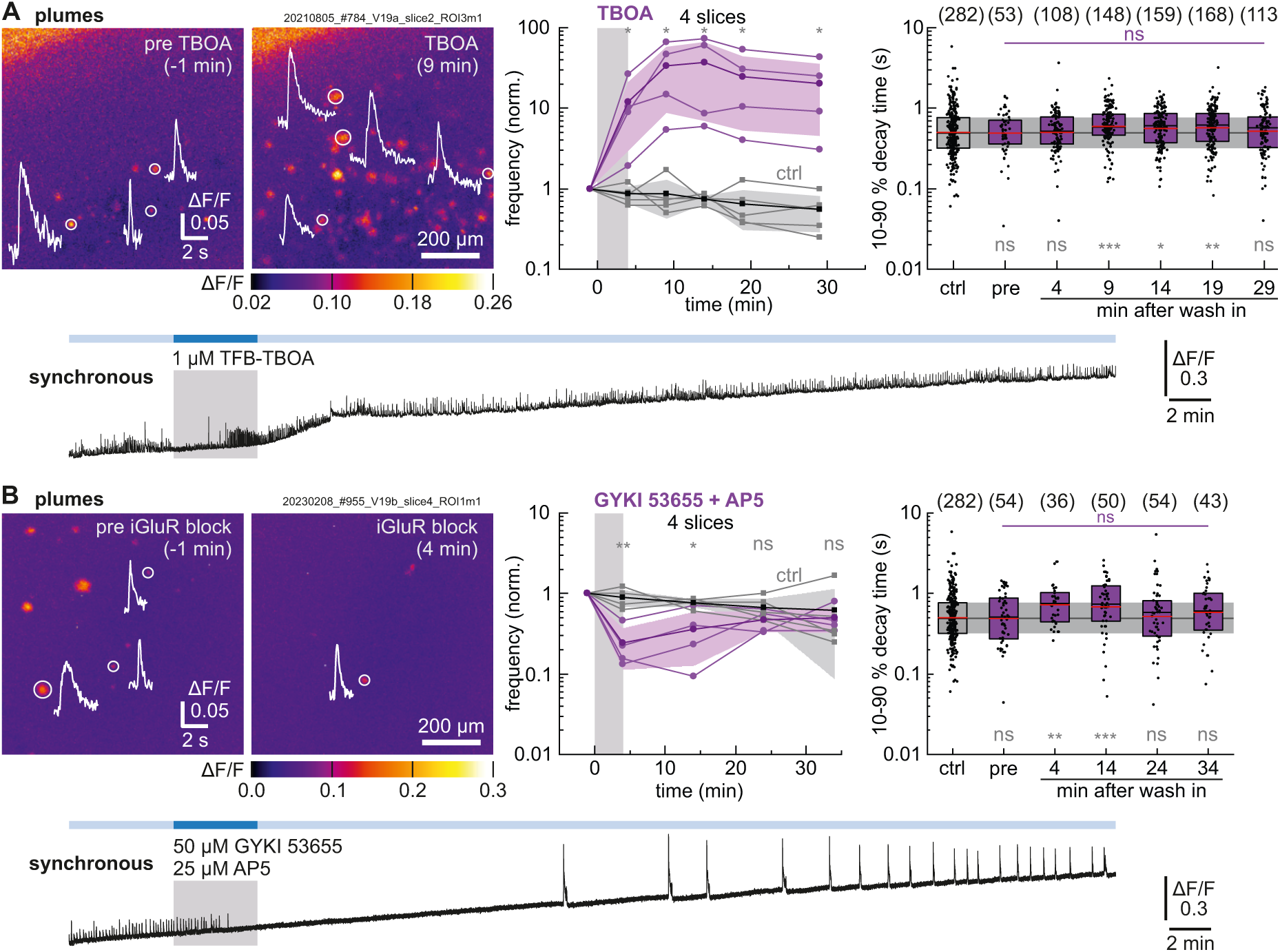
Plume characteristics after blocking glutamate uptake with TBOA or iGluR inhibition. (**A**) Addition of 1 μM TFB-TBOA affected both synchronous activity (bottom) and plumes (top left, maximal intensity projections, 25 s each with traces of selected plumes). Middle: Plume frequency at selected timepoints (n = 4 slices, purple) in comparison to control recordings without TBOA (n = 5 slices, black). Right: Plume 10-90% decay times for the indicated time windows (n = 7 slices). For details see **Fig. S17**. (**B**) Addition of 50 µM GYKI 53655 and 25 µM D-AP5 abolished synchronous activity (bottom) and reduced plumes (top left, maximal intensity projections, 50 s each with traces of selected plumes). Centre: Plume frequency at selected timepoints (n = 4 slices, purple) in comparison to control recordings without blockers (n = 5 slices, black). Right: Plume 10-90% decay times for the indicated time windows. See also **Fig. S18**. Individual data points are shown in light color, means in dark, and s.d. as shaded areas. Boxes show medians and 25-75% percentiles, red bars indicate the mean. Numbers in parentheses give the number of analyzed events. Mann-Whitney *U* tests were used for pair-wise comparison of individual time windows to control (grey; cf. **Fig. S14**), Kruskal-Wallis tests were used to test for differences between the six different time windows, followed by Dunn’s test (purple). ns not significant, \**p* < 0.05, \*\**p* < 0.01, \*\*\**p* < 0.001.

The strong frequency increase in plume occurrence seen with TFB-TBOA suggested that elevated extracellular glutamate concentrations themselves might contribute to the induction of plumes. This prompted us to test whether plume induction was mediated through iGluR activation. Indeed, inhibiting AMPA/non-NMDA receptors and NMDA receptors by coapplication of 50 μM GYKI 53655 and 25 μM D-AP5, respectively [Bleakman *et al*., 1996; Feuerbach *et al*., 2010], caused a strong reduction in plume frequency within 4 min (to 25 ± 13%, mean ± s.d., n = 4 slices), which returned to control levels after washout (**Fig. 5B** and **Fig. S18**). This suggests that iGluR activation does indeed contribute to plume formation.

### ‘Plume-like’ glutamate events are the main driver for glutamate accumulation during chemical ischemia

Having observed and characterized plumes in pre-ischemic baseline conditions, we next asked how these events change upon inducing chemical ischemia and how they contribute to the observed large-scale accumulation of glutamate (cf. **Fig. 1A,B**). For this we analyzed different episodes leading up to glutamate accumulation and during recovery, as shown for a typical example in **Fig. 6A-C** and **Fig. S19-20**.

**Figure 6.**
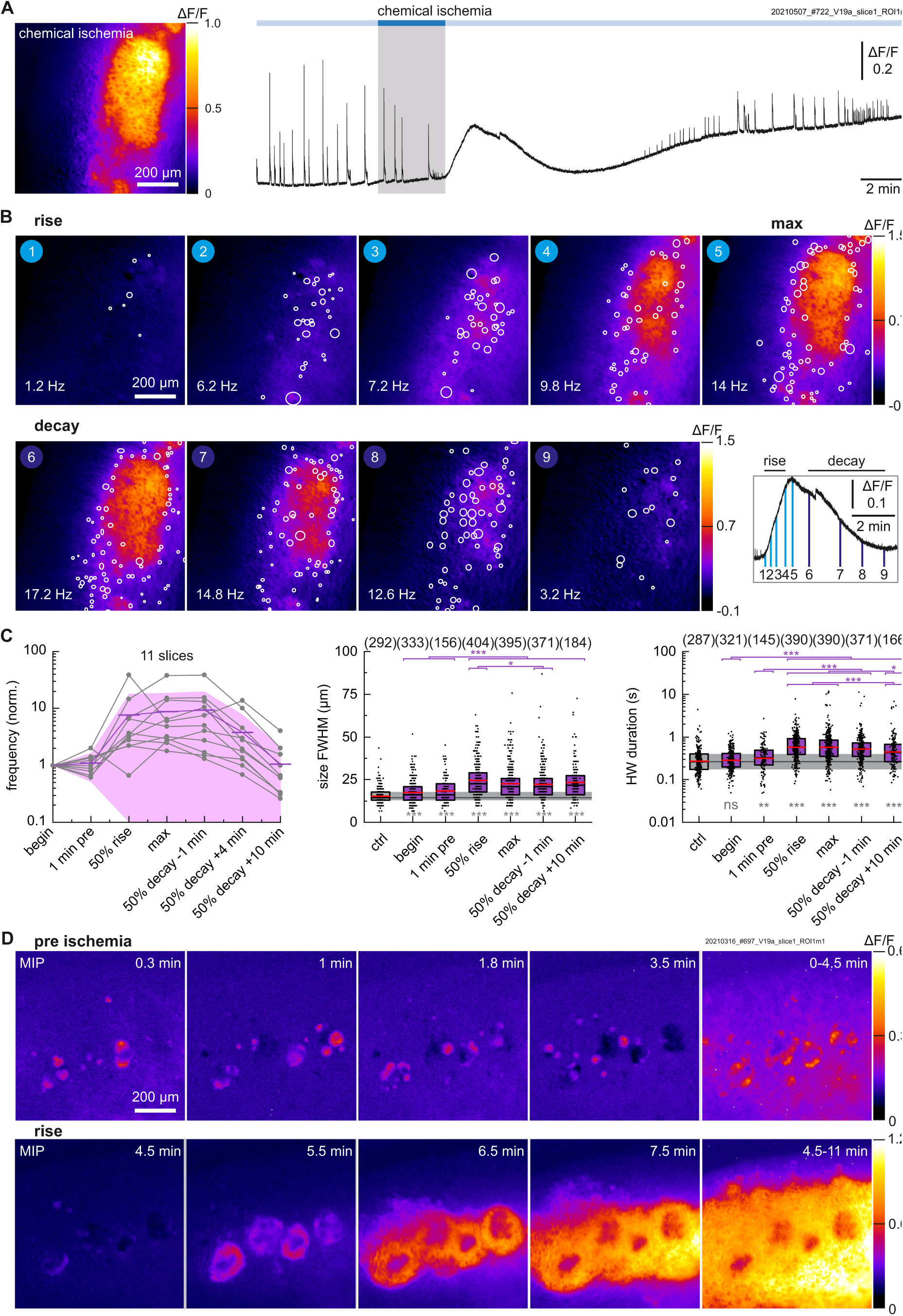
Plumes become more frequent and pronounced during chemical ischemia and are associated with defect regions. **(A)** Glutamate accumulation during chemical ischemia and ΔF/F trace of the responding region. **(B)** Localization and size of plumes (white circles, full-width) in different time windows during the ‘rise’ to maximal glutamate accumulation and the ‘decay’ period. The glutamate accumulation is shown in the background (minimum intensity projections, 5 s each). In addition, the respective plume frequency is given. The analyzed time windows are indicated on the right. See also **Fig. S19**. **(C)** Plume frequency and selected properties (see also **Fig. S20**). Individual data points are shown in light colors, means in dark, and s.d. as shaded areas. Numbers in parentheses give the number of analyzed plumes. Boxes show medians and 25-75% percentiles with red bars indicating the mean. Mann-Whitney *U* tests were used for pair-wise comparison of individual time windows to control (ctrl; grey cf. **Fig. S14**), Kruskal-Wallis tests were used to test for differences between the six different time windows, followed by Dunn’s test (purple). ns not significant, \**p* < 0.05, \*\**p* < 0.01, \*\*\**p* < 0.001. **(D)** Regions with high plume-incidence act as nuclei for ischemic glutamate accumulation. (Top) Pre-ischemia baseline imaging in a slice that had developed defects during culturing. Unusually large and frequent plumes were observed in the vicinity of defects (dark regions). (Bottom) Rise phase: Chemical ischemia caused large-scale glutamate accumulation, starting from the regions of high plume incidence. Shown are ΔF/F maximal intensity projections (MIP) of different time windows (10 s or longer time windows as indicated). See also **Fig. S21**.

As described above, the onset of the ischemic period was characterized by a loss of synchronous activity some time before extracellular glutamate accumulation occurred (**Fig. 6A**). After synchronous activity had ceased, more and more plumes were detected (**Fig. 6B**). Over time, regions with a high incidence of plumes showed a gradual fluorescence increase. When the global glutamate accumulation had reached 50% of the maximal accumulation (ΔF/F), the plume frequency was 7.0-fold higher compared to baseline (5 s or 25 s time windows, n = 11 slices, **Fig. 6C**). Eventually, the central regions reached a maximum fluorescence signal also meaning that individual events became indiscernible, possibly due to sensor saturation in these regions (**Fig. 6B**). However, plume-like dynamics were still seen at the edges of the central regions and plumes were abundantly detected in less saturated regions (see also **Movies S1, S7**). High plume activity was also detected when the fluorescence intensity gradually decreased during recovery (decay). The measured plume frequency only decreased later, i.e. at about 4 min after the accumulation had decayed to the 50% mark. Around that time also synchronous release events reappeared (**Fig. 1B**). Eventually, the plume frequency returned to baseline levels (**Fig. 6C**). Control measurements without inducing ischemia showed a steady decrease in plume frequency over 40 min (**Fig. S14**).

Chemical ischemia also had pronounced effects on plume size and duration. The plume size increased 1.4-fold, from 18 ± 7 μm before ischemia to 25 ± 10 μm during ischemia (at 50% rise, mean ± s.d.) (**Fig. 6C**). Plume HWD increased from 317 ± 331 ms before ischemia to 576 ± 592 ms (mean ± s.d.). This lengthening of the HWD was attributable to both, longer rise and longer plume decay times (**Fig. S20**). Despite the post-ischemic normalization of plume frequency, the plume size and also plume duration remained elevated at later time points, suggesting that persistent changes in glutamate homeostasis might have occurred.

This data shows that glutamate accumulation during chemical ischemia was mostly driven by local, plume-like events rather than by high-frequency, tetanic network activity. Ischemic conditions appear to favor the occurrence of plumes, as a high number of plumes became observable along with significant changes in their properties. Notably, plumes persisted during periods of maximal energy depletion.

Although causality is difficult to prove, it appeared that the plumes drive glutamate accumulation in organotypic slices. A notably example is also provided by a slice, which showed defect regions after cultivation (**Fig. 6D** and **Fig. S21**). In the vicinity of these defects, atypically large plumes were already observed at a high rate before inducing chemical ischemia (**Fig. 6D top**). Later, during chemical ischemia, these defect regions acted as nuclei for sustained glutamate accumulation (**Fig. 6D bottom**).

Since we had found that plumes can be suppressed by inhibiting iGluRs (**Fig. 5B**), we used this to test their contribution to extracellular glutamate accumulation during chemical ischemia. Indeed, pre-application of 50 μM GYKI 53655 and 25 μM AP5 for 4 min did fully suppress the plume frequency increase and glutamate accumulation during 5 min chemical ischemia (**Fig. 7A,B**). Both, plumes and glutamate accumulation, were seen when we induced a second control chemical ischemia after removing the iGluR inhibitors. This suggested that iGluRs play a key role in driving plume formation and that plumes eventually cause the accumulation of glutamate in metabolic stress conditions.

**Figure 7.**
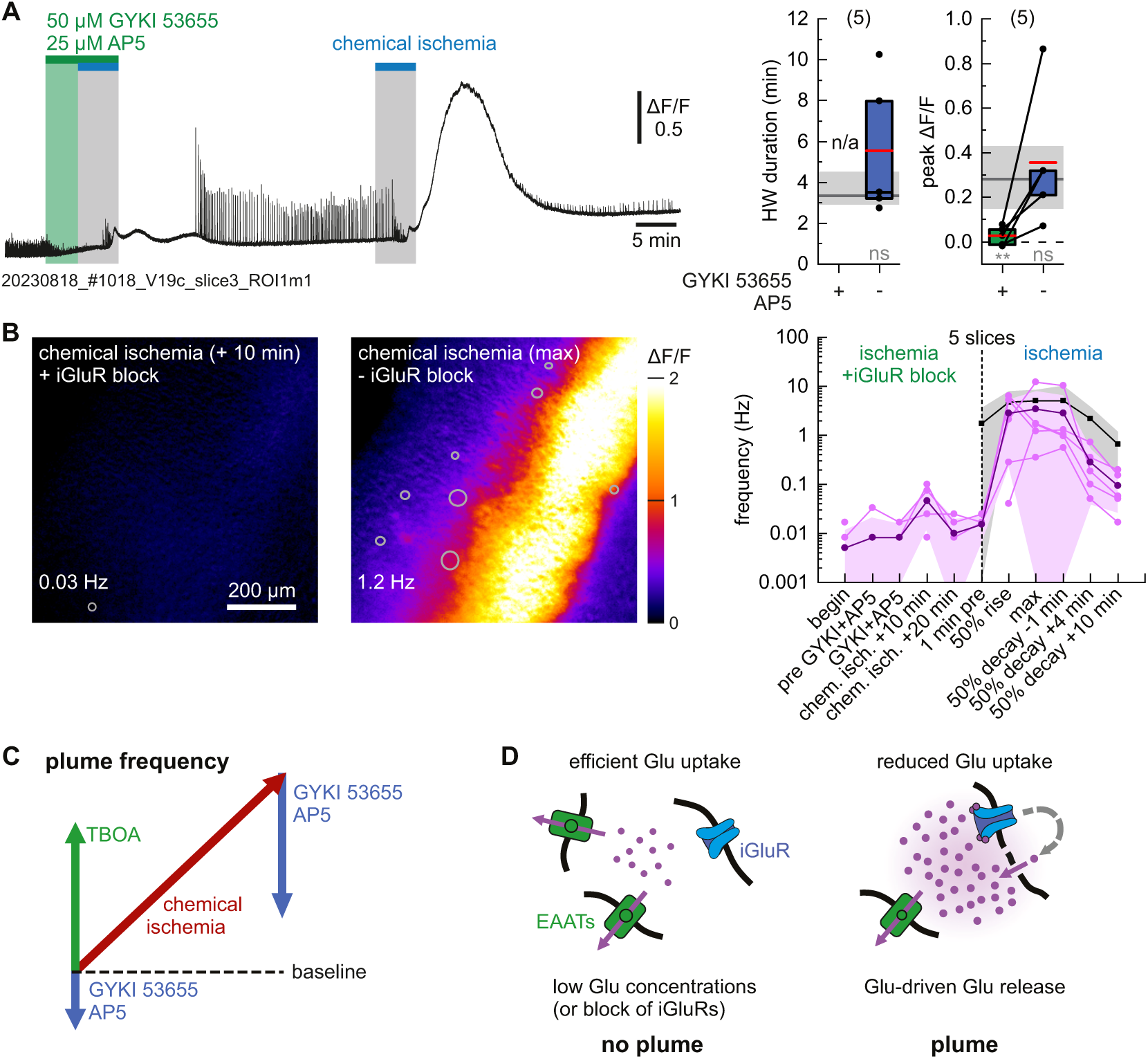
Chemical ischemia in the presence of iGluR inhibitors and graphical summary. **(A)** (Left) Glutamate dynamics upon inducing chemical ischemia in the presence of iGluR inhibitors (50 μM GYKI 53655 and 25 μM D-AP5) followed by a 2^nd^ chemical ischemia without inhibitors. Shown is ΔF/F of the responding region. (Right) Half-width duration of glutamate accumulation and peak ΔF/F in responding regions of n = 5 slices. Boxes show medians and 25-75% percentiles with red bars indicating the mean. Mann-Whitney *U* tests were used for pair-wise comparison to control (grey, cf. Fig. 1C), ns not significant, \**p* < 0.05, \*\**p* < 0.01. **(B)** (Left) Minimum intensity projections (25 s) of glutamate accumulation with plume localization (circles; full-width) and plume frequencies. (Right) Plume frequency at the indicated time points (purple). Individual data points are shown in light colors, means in dark, and s.d. as shaded areas. The black/grey curves show the data of previous chemical ischemia experiments (cf. Fig. 6C). **(C)** Observed changes in plume frequency: Under baseline conditions the number of observable plumes was low, but it increased upon blocking glutamate uptake with TBOA, and it was reduced upon inhibiting iGluRs with GYKI 53655 and AP5. Chemical ischemia caused a strong increase in plume frequency, which was strongly reduced by adding GYKI 53655 and AP5. **(D)** Model: Plume formation seems to be at least partially driven by elevated extracellular glutamate concentrations and iGluRs, i.e. by an ‘auto-catalytic’ recruitment of additional glutamate sources.

## Discussion

In this study we used chemical ischemia in organotypic slices to investigate how metabolic stress affects network activity and glutamate homeostasis. Previous work in this paradigm had shown that brief energy depletion causes strong, yet reversible effects, namely astrocyte depolarization, increases of intracellular Na^+^ and extracellular K^+^ concentrations [Lerchundi *et al*., 2019a; Eitelmann *et al*., 2022; Meyer *et al*., 2022], and intracellular ATP depletion [Pape & Rose, 2023]. Additional measurements with the fluorescent ATP sensor ATeam1.03^YEMK^ performed here (**Fig. S1**), in combination with earlier *in situ* calibrations [Lerchundi *et al*., 2020] and assuming a resting ATP concentration of 2.8 mM [Pape & Rose, 2023], indicated that inducing chemical ischemia for 2 min resulted in a transient decrease in ATP concentration by ~1.2 mM. Inducing chemical ischemia for 5 min caused a maximal drop in ATP concentration by ~2.6 mM, i.e. almost complete ATP depletion. Still, neurons almost fully regained their pre-ischemic ATP levels. Similarly, all other effects investigated here, i.e. the effect of chemical ischemia on neuronal network activity, neuronal Ca^2+^ loading, and extracellular glutamate dynamics, were mostly reversible (**Fig. 1**). This suggests that our protocol mimics mild to moderate transient metabolic stress as it is encountered in the penumbra of the infarct core experiencing SDs [Gerkau *et al.,* 2018; Dreier & Reiffurth, 2015; Pietrobon & Moskowitz 2014].

### Monitoring synchronous network activity with SF-iGluSnFR before and during chemical ischemia

Widefield SF-iGluSnFR(A184V) imaging reported on synchronous network activity similar to GCaMP6f (**Fig. 1** and **Fig. S7/8**). As expected, the measured glutamate transients decayed faster than the intracellular GCaMP6f signals. In some cases, bursting behavior was observed, but these events were not fully resolved at our imaging rates of 20 fps and 99 fps (**Fig. S11**). Also, the measured ΔF/F peak intensities may thus remain underestimated. The fast kinetics are in line with the high temporal resolution of SF-iGluSnFR(A184V) and are also seen in measurements of evoked release events [Marvin *et al*., 2018; Helassa *et al*., 2018; Armbruster *et al*., 2016; Barnes *et al*., 2020, Herde *et al*., 2020].

Our SF-iGluSnFR(A184V) measurements of synchronous activity were sensitive enough to detect an increased glutamate release upon recovery from TTX applications (**Fig. 4A** and **Fig. S15A)**. This increase could either be explained by rapid upscaling of synaptic glutamate release or network effects, and also neuronal Ca^2+^ influx was strongly increased after TTX treatment (**Fig. S15B)**. Similarly, blocking GABA_A_ receptors with gabazine increased the amount of released glutamate (**Fig. 4B** and **Fig. S16)**. This can be explained by either higher burst frequencies, prolonged bursts and/or inclusion of more neurons into the network due to disinhibition [Dawitz *et al*., 2020]. In the early phase of gabazine addition, the latter two explanations appear to be more likely, since the highest signal amplitudes were observed while the event frequency was still low.

As expected, blocking glutamate uptake with TFB-TBOA had pronounced effects on synchronous events (**Fig. 5A** and **Fig. S17**). Half-width duration, decay times and event amplitudes were strongly increased, all of which can be explained by reduced EAAT-mediated glutamate uptake [Marvin *et al*. 2013; Armbruster *et al*. 2016; Barnes *et al*. 2020]. In some slices, TFB-TBOA eventually reduced the frequency of synchronous events (**Fig. S17**), which indicates that synaptic transmission became impaired. Finally, blocking iGluRs with GYKI 53655 and D-AP5 resulted in a complete loss of network activity, which, after washout, resumed with enhanced glutamate transients at low frequency (**Fig. 5B** and **Fig. S18**). In summary, we find that SF-iGluSnFR(A184V) faithfully reports on synchronous network activity in slice cultures. This could be further utilized to study network maturation, synaptic homeostasis, synchronicity, or epileptiform activity.

We next focused on how synchronous activity is affected by chemical ischemia. Upon transient energy depletion, synchronous SF-iGluSnFR signals were lost (**Fig. 1**), which was mostly due to a loss of signal amplitude: The signal amplitudes gradually decreased, while in some slices the event frequency increased in this period (**Fig. S7**). This suggests that energy restriction has an immediate effect on the amount of released glutamate. This is not surprising given that vesicle filling, docking and replenishment are energy-costly processes [Attwell & Laughlin, 2001]. In contrast, the synchronous activity reported by GCaMP6f did not cease at this point, but turned into high-frequency Ca^2+^ influx events that resulted in intracellular Ca^2+^ loading. The fact that Ca^2+^ activity continued despite reduced glutamate release suggests that the network remained functional for some more time. Oscillatory behavior has also been described in the context of anoxic depolarizations (e.g. [Dzhala *et al.,* 2001]. At later stages, network desynchronization may occur and cell autonomous high-frequency firing may originate from permanent, above-threshold depolarization (e.g. [Kalia *et al.,* 2021] and **Fig. S5**). Future dual-color imaging experiments with fast, red-fluorescent Ca^2+^ sensor variants might allow to better dissect the timing of these processes.

Our 2-5 min chemical ischemia protocol was short enough to allow for recovery of synchronous activity in all slices (**Fig. 1** and **Fig. S7**). Post-ischemic glutamate transients showed increased decay times, yet a bias due to lower signal intensities cannot be excluded. However, also post-ischemic Ca^2+^ events, despite having similar signal intensity, showed increased decay times, which indicates that the chemical ischemia protocol might have caused some persistent effects, e.g. extracellular space changes due to cell swelling [Juzekaeva *et al*., 2020; Meyer *et al.,* 2022] and/or impairment of astrocyte function [Risher *et al*., 2012; van Putten *et al*., 2021; Eitelmann *et al.,* 2023]. Interneurons, which are thought to be particularly sensitive to energy deprivation and excitotoxicity [Meade *et al*., 2000; Epsztein *et al*., 2006], survived the protocol at least partially, as confirmed by our GCaMP6f-imaging experiments with the interneuron-specific mDlx enhancer (**Fig. S9**). During recovery, synchronous Ca^2+^ events showed another high-frequency period, around the time when the first glutamate events began to reappear. Apparently, network activity resumed, although the amount of released glutamate had not reached normal levels, yet. SF-iGluSnFR variants with higher affinity or improved synaptic localization [Aggarwal *et al*., 2023; Hao *et al*., 2023] might provide higher sensitivity to capture these processes in more detail.

### Plumes are atypical glutamate release events which are rarely seen under normal conditions

Besides synchronous activity, SF-iGluSnFR(A184V) widefield imaging revealed heterogenous and locally confined glutamate release events. We refer to these events as ‘plumes’, which reflects their spontaneous occurrence, slow kinetics and relatively large size (**Fig. 2** and **Fig. 3**). This term was recently introduced by Parker *et al*. [Parker *et al*., 2021; Eikermann-Haerter, 2021], who observed a similar phenomenon in 2-photon *in vivo* imaging experiments in a familial hemiplegic migraine type 2 (FHM2) mouse model with reduced astrocytic Na^+^/K^+^-ATPase function.

Glutamate plumes showed remarkable features: Plumes were not correlated with network activity but occurred randomly, which suggests that they are triggered locally. In line with this, the plume frequency neither decreased, when we suppressed network activity with TTX (**Fig. 4** and **Fig S15**), nor did it increase, when we strengthened network activity by adding gabazine (**Fig. S16**). Plumes occurred in the presence of TTX, i.e. they do not require Na_v_ channel activation, which distinguishes them from action potential-driven synaptic glutamate release. Also, the heterogeneity of plumes, their relatively large size (FWHM 17.2 ± 6.4 μm) and long duration (several hundred milliseconds) sets them apart from synaptic release events. The slow kinetics of plumes are particularly striking. Plumes decayed far more slowly than the transients associated with synchronous network activity, although the latter had higher overall signal amplitudes and report on synchronous and bursting release from thousands of synapses across the whole slice. Also published measurements of evoked release with various iGluSnFR variants show much faster decay kinetics than plumes [Marvin *et al*., 2018; Helassa *et al*., 2018; Armbruster *et al*., 2016; Jensen *et al*., 2019; Matthews *et al.,* 2022]. Spontaneous release events [Hao *et al*., 2023] or miniature events, which are also TTX-insensitive, give rise to even faster iGluSnFR signals [Aggarwal *et al*., 2023; Farsi *et al*., 2021].

The amount of glutamate that is released in individual plumes is hard to estimate, since SF-iGluSnFR(A184V) is a non-ratiometric sensor, the resolution in *z* dimension was limited, and glutamate calibrations are difficult to perform *in situ* (**Fig. S3**). However, based on the affinity of SF-iGluSnFR(A184V) (reported EC_50_= 2.1 μM) [Marvin *et al*., 2018], it seems likely that during plumes glutamate reached hundred nanomolar to low micromolar concentrations. Given the large size and slow decay of plumes, it seems unlikely that this amount of glutamate originates from single sources, such as synaptic boutons of individual neurons. More quantitative insight might be provided by measurements with additional low-affinity SF-iGluSnFR variants, imaging with higher spatial resolution, or modeling, also taking glutamate buffering by SF-iGluSnFR into account [Armbruster *et al*., 2020]. In any case, we presume that plumes either originate from very large individual glutamate sources like astrocytes and/or multiple cells within a region, possibly linked by a feed-forward mechanism.

Support for a feed-forward mechanism comes from our experiments, in which we artificially increased the glutamate levels by blocking glutamate uptake. When we used TFB-TBOA to block EAAT1-3, the plume frequency increased 6- up to 73-fold (**Fig. 5A** and **Fig. S17**). Besides this, TFB-TBOA caused a sustained 1.4-fold increase in the mean plume size, whereas plume half-width duration, decay time and intensity (ΔF/F) remained unaffected. The fact that TFB-TBOA had a direct effect on plume frequency but less on plume characteristics indicates that increased glutamate concentrations are important for plume induction/formation (**Fig. 7C,D**), whereas EAAT function appears to be less relevant for plume termination and dissipation/removal of the released glutamate.

Last, the number of plumes was rapidly reduced upon inhibiting iGluRs, namely AMPA and NMDA receptors, by adding GYKI 53655 and D-AP5, respectively (**Fig. 5B**). This supports the findings obtained with TFB-TBOA and suggests that glutamate itself is a key driver for plume formation by activating iGluRs (**Fig. 7C,D**). Once more, this distinguishes plumes from miniature release events, which should be insensitive to iGluR block. Importantly, pharmacological inhibition of iGluRs provides an experimental way to suppress plumes. NMDA receptors, which are activated by low micromolar glutamate concentrations [Reiner & Levitz, 2018], might be key contributing factor, but also AMPA receptors (and possibly also mGluRs activation), should be considered further.

It should be noted that plumes were observed under baseline conditions, albeit rarely. The observed frequencies were low and varied between slices (**Fig. S12**) and a substantial number of slices showed no plumes. Overall, we detected plumes in approx. 30% of the investigated slices under baseline conditions. Higher SF-iGluSnFR expression levels might have increased the detection probability for plumes.

Parker *et al*., who performed 2-photon *in vivo* imaging experiments reported that plumes were mostly absent in wildtype animals but detected plumes in FHM2 mice, mostly in superficial cortical layer 1, but also layer 2/3 [Parker *et al*., 2021]. They further showed that the occurrence of plumes was related to a reduced number of astrocytes and EAAT2 (GLT-1) expression in these regions. Similarly, our organotypic slice cultures may not fully recapitulate the situation encountered in healthy brain tissue *in vivo*. For instance, reactive astrocytes, which may have reduced glutamate uptake capacity, can be overly present in slice cultures, mainly on the surface [Benediktsson *et al*., 2005; Schreiner *et al*., 2023; Lerchundi *et al*., 2019b].

Further pharmacological experiments may help to differentiate whether plumes originate from vesicular or non-vesicular release. Besides activity-dependent vesicular release from the presynapse, other glutamate release mechanisms have been described, such as vesicular glutamate release from astrocytes/glia [Malarkey & Parpura, 2008; Hamilton & Attwell, 2010; de Ceglia *et al*., 2023], or release through other channels such as P2X receptors, bestrophin-1 (Best1) or Swell1 [Yang *et al*., 2019; Owji *et al*., 2022]. Notably, even when EAATs are blocked with TBOA, glutamate maintains to have a strongly depolarizing effect on astrocytes through NMDA receptor activation and elevated extracellular K^+^ concentrations [Sriavastava *et al*., 2020].

### Plumes drive glutamate accumulation during chemical ischemia

Inducing metabolic stress caused a strong increase in plume frequency (**Fig. 6B**). Eventually, these local and asynchronous events led to extensive glutamate accumulation covering large areas of the slices. Chemical ischemia did not only increase the plume frequency, but also plume size, signal intensity, duration and decay times were significantly increased (**Fig. 6C** and **Fig. S19,20**). As such, energy depletion elicited far stronger effects than just blocking glutamate uptake with TFB-TBOA.

It appears that plumes were the main source of extracellular glutamate accumulation during chemical ischemia: Synchronous activity had already ceased at this time (**Fig. 1**) and regions with high plume activity were the first to reach plateauing signals, which most likely reflect sensor saturation (**Fig. 6**). Plumes reappeared in these regions, as soon as the extracellular glutamate concentration began to drop in the recovery phase. At the edges of these saturated regions, plumes were visible throughout the ischemic period. Apparently, plumes persisted throughout energy depletion, which indicates that the underlying release mechanisms are not strongly reliant on continuous energy supply. At later stages the plume frequency normalized, but plume sizes and durations remained increased compared to pre-ischemic conditions, which again points to some persistent changes, as earlier described for synchronous activity.

Another link between plumes and glutamate accumulation was found in a slice with defect regions. There the defect regions clearly served as nuclei for further glutamate accumulation (**Fig. 6D**). Quite strikingly, inducing chemical ischemia in the presence of GYKI 53655 and D-AP5, did not only suppress the occurrence of plumes but also suppressed ischemic glutamate accumulation (**Fig. 7**). Importantly, this experiment demonstrates that extracellular glutamate accumulation is not merely caused by ion disbalances but shows that glutamate itself plays a driving role for glutamate accumulation in the acute ischemic phase.

### Glutamate plumes may have broader pathological significance

Glutamate accumulation has long been considered to be a key factor in ischemia and cortical SDs [Andrew *et al*., 2022; Dreier & Reiffurth, 2015; Pietrobon & Moskowitz, 2014], the latter also occurring in the context of brain injuries, migraine and epilepsy. Indeed, previous *in vivo* SF-iGluSnFR measurements demonstrated wave-like glutamate accumulations after SD induction through focal KCl application [Enger *et al*., 2015; Rakers *et al*., 2017]. In addition, Parker *et al*. observed glutamate plumes which preceded the SD onset at the initiation site and observed plumes again in the decaying glutamate wavefront [Parker *et al*., 2021]. Still, it remains debated how much elevated extracellular glutamate levels contribute to SD initiation and spreading, also in comparison to extracellular K^+^ waves, cellular depolarization, and Ca^2+^ elevations [Andrew *et al*., 2022; Pietrobon & Moskowitz 2014].

We here found evidence that increased extracellular glutamate concentrations can trigger more glutamate release, specifically, in form of plume-like events. We further found that this mechanism operates under energy-scarce conditions and in the absence of network function. It is thus able to mediate large-scale glutamate accumulation in the acute phase of metabolic stress. We further showed that iGluR activation and reduced glutamate uptake play important roles in triggering plumes, while the cellular sources and release mechanisms that cause plumes remain to be identified.

Given our observations, it seems likely that plumes do not only occur upon severe disruptions of glutamate homeostasis, but also in the context of more subtle disbalances. Plumes were even observed under baseline conditions in our slice cultures but also in FHM2 mice [Parker *et al*., 2021]. Plumes may thus become significant as soon as glutamate uptake becomes suboptimal, e.g., due to transporter malfunction, increased metabolic demands, reduced astrocyte function, or reduced neurovascular coupling [Tarantini *et al*., 2017]. Reduced glutamate uptake, for instance, has been observed in the context Alzheimer’s disease, where, in a mouse model, A*β* plaques caused a local reduction of EAATs that slowed glutamate uptake and enhanced transients after evoked glutamate release [Hefendehl *et al*., 2016] (see also [Zott *et al*., 2019; Brymer 2023]). Similar effects have been reported for a model of Huntington’s disease [Dvorzhak 2019] and might cooccur with age, as brain integrity becomes compromised. Other diseases are thought to involve increased glutamate release (glutamatergic hyperactivity), such as Parkinson’s disease or inflammations [Campanelli *et al*., 2022]. As we show here, increased glutamate concentrations are sufficient to provide enough feed-forward drive for triggering plumes. Individual plumes could interfere with synaptic function or may not cause any physiological effects at all. However, plumes can easily result in the accumulation of larger amounts of extracellular glutamate, which will inevitably affect network function or might even contribute to excitotoxicity. Thus, more research seems warranted, both, on the origin and actions of plume-like glutamate release events, and, the role of glutamate uptake deficiencies as a contributing factor in neurological diseases.

## Material and Methods

### Preparation of organotypic slice cultures and transduction with rAAVs

Animals for post-mortem tissue removal were obtained from the Animal Facility of the Faculty of Biology and Biotechnology, Ruhr University Bochum, approved by the Landesamt für Natur, Umwelt und Verbraucherschutz (LANUV) according to §11 TierSchG. There, animals were bred and housed under standard conditions with a 12 h light/dark cycle and food and water provided *ad libitum*. All efforts were taken to minimize animal suffering.

Organotypic slice cultures were prepared from male and female CB57BL/6 mice on postnatal day 7-9 following a previously published protocol [Lerchundi *et al*., 2019b]. The animals were sacrificed by decapitation and the brain was quickly transferred to ice-cold Ringer’s solution (125 mM NaCl, 2.5 mM KCl, 1.25 mM NaH_2_PO_4_, 26 mM NaHCO_3_, 2 mM CaCl_2_, 1 mM MgCl_2_ and 20 mM D-glucose; 95% O_2_ and 5% CO_2_). The hemispheres were separated and a 45° parasagittal cut (see **Fig. S22A**) was performed before cutting 250 µm slices using a vibratome (Leica VT1200; 1 mm amplitude, 0.9-1.0 mm/s, 15° angle) in ice-cold Ringer’s solution. Then, the hippocampus with the adjacent cortex was excised and the slices were stored in 34°C Ringer’s solution until the second hemisphere was sliced. Under sterile conditions, slices were washed five times in HBSS (Sigma, H9394) with minimal solution transfer. Finally, the slices were placed on membrane inserts (Millicell PICM0RG50, hydrophilized PTFE, pore size 0.4 µm) for cultivation according to Stoppini [Stoppini *et al*., 1991; Gee *et al*., 2017]. The slices were supplied with organotypic slice culture medium (20% heat-inactivated horse serum, GIBCO/Life Technologies; 1 mM L-glutamine, 0.001 mg/ml insulin, 14.5 mM NaCl, 2 mM MgSO_4_, 1.44 mM CaCl_2_, 0.00012 % ascorbic acid and 13 mM D-glucose in MEM (Sigma, M7278) and incubated at 37°C and 5% CO_2_. A full medium exchange was performed three to four times a week. In the first 24 h after slicing, the slices, which encompassed portions of cortex and hippocampus, showed swelling. Until DIV (day *in vitro*) 7 slices deswelled, became translucent and started to flatten, with some cells migrating outwards.

On DIV 1, the slices were transduced with rAAVs of serotype 8 produced in house. pAAV.hSynapsin.SF-iGluSnFR(A184V) was a gift from Loren Looger (Addgene plasmid #106175; [Marvin *et al*., 2018]), pAAV.Syn.GCaMP6f.WPRE.SV40 was a gift from Douglas Kim & GENIE Project (Addgene plasmid #100837; [Chen *et al*., 2013]) and pAAV.mDlx.GCaMP6f.WPRE.SV40 was a gift from was a gift from Gordon Fishell (Addgene plasmid #83899; [Dimidschstein *et al*., 2016]).

rAAVs were produced in HEK293Tsa cells (for details see **Supplementary Methods** and **Fig. S22B**). AAV particles were diluted 1:3, 1:5 or 1:7 and for slice culture transduction, 1 µl was added at the center of the cortical region.

### Imaging, chemical ischemia and pharmacological manipulations

Imaging experiments were performed at DIV 14-21. Slices were placed in a custom-made chamber and superfused with Ringer’s solution (see Preparation section) at 1-3 ml/min and 24 °C using a peristaltic pump (Gilson Minipuls 3) and in-line heater, respectively. The chamber was placed under an upright microscope (Zeiss Axioscope) fitted with a 10x/0.3 water immersion objective (Zeiss N-Achroplan M27). Epifluorescence excitation was provided by a xenon light source (Sutter Instruments, Lambda DG-4) with a 470/40 nm excitation filter and a light-guide (Thorlabs, LLG3-6H), a 495 nm dichroic mirror (AHF, T495LPXR) and a 525/39 nm emission filter (Semrock, 525/39 BrightLine HC). For high-speed imaging the DG4 and light guide were replaced by a collimator-coupled LED (Thorlabs, M470L4) using the same filter set. The light intensity in the sample plane was adjusted to ~0.4 mW/mm^2^. Images were acquired at 20 fps or 99 fps for glutamate imaging and at 10 fps for calcium imaging with an EMCCD camera (Photometrics, Evolve 512delta) at 16 bit in pre-sequence clearing mode with 512 x 512 or 256 x 256 pixels using MicroManager2.0 [Edelstein *et al*., 2014]. At 256 pixels one pixel corresponded to 3.22 µm.

The slices were allowed to equilibrate in the experimental chamber for 15-20 min prior to imaging. Typically, 5 min baseline imaging was performed before blockers were added or chemical ischemia was evoked by applying an ischemic Ringer’s solution (with 0 mM D-glucose, 2 mM 2-deoxyglucose and 5 mM NaN_3_) for 2-5 min. For pharmacological characterization 1 µM TFB-TBOA (Tocris, 2532), 0.2 µM TTX (Tocris, 1078), 3 µM GBZ (SR 95531; Tocris, 1262), 50 µM GYKI 53655 (HelloBio, HB0312) or 25 µM D-AP5 (HelloBio, HB0225) in Ringer’s solution were applied for 4 min. The shown application bars are corrected for the experimentally determined 1 min dead-time. Typical ischemia or pharmacological recordings lasted up to 50 min, experiments with a second ischemia (**Fig. 7**) up to 1.5 h. Only healthy slices (with exception of the slice shown in **Fig. 6D**) with high SF-iGluSnFR(A184V) expression levels across all cortical layers and synchronous activity before and after drug application were considered for analysis. Since we were interested in analyzing plumes, we selected slices with baseline plume frequencies ≥0.12 Hz for control, TTX, GBZ and GYKI 53655/D-AP5 recordings. For ischemia experiments all slices had baseline plume frequencies ≥0.06 Hz.

### Data analysis, presentation, and statistics

The obtained images were processed using FIJI 1.53t [Schindelin *et al*., 2012]. Movies acquired with 512 x 512 pixels were binned to 256 x 256 pixels by averaging. ΔF/F denotes relative fluorescence changes, which were calculated with F_0_ as an average of 10-200 frames with no synchronous activity. Typical time windows for analysis were 500 frames with a local F_0_; in cases of low frequencies, stack sizes were increased up to 6000 frames. Subsequently, stacks were processed with the ImageJ built-in image calculator to obtain ΔF/F stacks.

For measuring glutamate and Ca^2+^ accumulation as well as synchronous activity the responding area was manually selected as region of interest (polygon). For plume analysis, maximum intensity projections (MIPs) were calculated after excluding all frames with synchronous activity by setting them to zero intensity. The obtained MIPs served to identify possible plumes, which were manually confirmed in the corresponding image stack. For further analysis of each plume, a circular region of interest was manually defined at the peak frame (maximal plume intensity).

The mean ΔF/F of all regions of interest (synchronous activity or plumes) was exported as time series to ClampFit 11.1 (Molecular Devices) to analyze the traces for peak frames, max ΔF/F, HW duration and 10-90 % rise and decay times. Excel (Microsoft) was used to calculate logarithmic values for durations and rise and decay times. At 20 fps bursting synchronous activity remained unresolved (see also **Fig. S11**) and contributed to the corresponding decay times. Plume sizes were determined in the corresponding maximum intensity frame (full-width at half-maximum; FWHM) at the center of the plume in x- and y-direction (**Fig. 3A**). Plumes were mostly round (**Fig. 3B**) but the size (FWHM) was calculated as an average of the x- and y-size to account for asymmetries. Plumes were only included into analysis, if they possessed a ΔF/F signal change of at least 2%, lasted >50 ms (HW duration) and were at least 2×2 pixels in size. Small or brief events might thus have escaped our detection. Some long-lasting plumes might have been missed when they overlapped with synchronous activity or occurred during overall increasing glutamate concentrations. We also noted that the ability to detect plumes depended on the expression density of SF-iGluSnFR, i.e. plumes were hardly detected in poorly transduced slices.

In all measurements, both in slice cultures and HEK cells, we noted a run-up of the SF-iGluSnFR baseline intensity of ∽30% over 40 min continuous imaging, which was apparently accompanied by a decrease of the signal change ΔF/F. The origin of this effect is presently unclear, but it complicated our efforts to calibrate the sensor by applying defined glutamate concentrations.

All data were statistically tested for normality (including frequencies) or log normality (for durations, rise and decay times) with the Kolmogorov-Smirnov test (*p* <0.05) using Origin Pro 2021 (OriginLabs). As several datasets were not normally distributed, all experimental groups were compared with the Kruskal-Wallis test followed by posthoc Dunn’s test (not significant ns, \**p* <0.05; \*\**p* <0.01; \*\*\**p* <0.001). Furthermore, duration, rise and decay times and plume sizes were tested individually against a control group with the Mann-Whitney *U* test (not significant ns, \**p* <0.05; \*\**p* <0.01; \*\*\**p* <0.001). Representative ΔF/F images are displayed with the ImageJ look-up table ‘fire’ and the indicated scaling. Figures were assembled in Corel Draw 2018 (Corel). For detailed descriptions of analysis and data presentations in each figure, see **Supplementary Methods**.

## Acknowledgements

We thank the staff of the animal facility and Silvia Schweer for technical assistance at Ruhr University Bochum. Tatjana Surdin, Stefan Herlitze and Melanie Mark (Ruhr University Bochum) provided advice on rAAV production. Nina Dietzel contributed to SF-iGluSnFR pH control measurements. We also thank Claudia Roderigo and Simone Durry for technical assistance at Heinrich Heine University. Last we thank all members of the DFG Research Unit 2795 “*Synapses under stress*“, which provided the framework for this project.

## Availability of data and code

Data obtained in this study can be made available upon reasonable request. Simulation code for the Supplementary Material is available at https://gitlab.utwente.nl/m7686441/focalglutamate_modelsimulations

## Author contributions

TZ and AR designed the overall project. TZ and JSEN established organotypic slice cultures at RUB. TZ conducted all glutamate and Ca^2+^ imaging experiments and analyzed them with AR. NP performed ATP imaging experiments at HHU that were designed and supervised by CRR. FIMA, MK and HGEM performed simulations. AR and TZ wrote the manuscript with input from all authors.

## References

Aggarwal A, Liu R, Chen Y, Ralowicz AJ, Bergerson SJ, Tomaska F, Mohar B, Hanson TL, Hasseman JP, Reep D, Tsegaye G, Yao P, Ji X, Kloos M, Walpita D, Patel R, Mohr MA, Tillberg PW, Looger LL, … Podgorski K. (2023). Glutamate indicators with improved activation kinetics and localization for imaging synaptic transmission. Nature Methods 20: 925–934.

Andersen JV, Markussen KH, Jakobsen E, Schousboe A, Waagepetersen HS, Rosenberg PA, & Aldana BI. (2021). Glutamate metabolism and recycling at the excitatory synapse in health and neurodegeneration. Neuropharmacology 196: 108719.

Andrew RD, Farkas E, Hartings JA, Brennan KC, Herreras O, Müller M, Kirov SA, Ayata C, Ollen-Bittle N, Reiffurth C, Revah O, Robertson RM, Dawson-Scully KD, Ullah G, & Dreier JP. (2022). Questioning glutamate excitotoxicity in acute brain damage: The importance of spreading depolarization. Neurocritical Care 37: 11–30.

Armada-Moreira A, Gomes JI, Pina CC, Savchak OK, Gonçalves-Ribeiro J, Rei N, Pinto S, Morais TP, Martins RS, Ribeiro FF, Sebastião AM, Crunelli V, & Vaz SH. (2020). Going the extra (synaptic) mile: Excitotoxicity as the road toward neurodegenerative diseases. Frontiers in Cellular Neuroscience 14: 90.

Armbruster M, Dulla CG, & Diamond JS. (2020). Effects of fluorescent glutamate indicators on neurotransmitter diffusion and uptake. ELife 9: e54441.

Armbruster M, Hanson E, & Dulla CG. (2016). Glutamate clearance is locally modulated by presynaptic neuronal activity in the cerebral cortex. Journal of Neuroscience 36: 10404–10415.

Attwell D, & Laughlin SB. (2001). An energy budget for signaling in the grey matter of the brain. Journal of Cerebral Blood Flow and Metabolism 21: 1133–1145.

Barnes JR, Mukherjee B, Rogers BC, Nafar F, Gosse M, & Parsons MP. (2020). The relationship between glutamate dynamics and activity-dependent synaptic plasticity. Journal of Neuroscience 40: 2793–2807.

Belov Kirdajova D, Kriska J, Tureckova J, & Anderova M. (2020). Ischemia-triggered glutamate excitotoxicity from the perspective of glial cells. Frontiers in Cellular Neuroscience 14: 51.

Benediktsson AM, Schachtele SJ, Green SH, & Dailey ME. (2005). Ballistic labeling and dynamic imaging of astrocytes in organotypic hippocampal slice cultures. Journal of Neuroscience Methods 141: 41–53.

Bleakman D, Ballyk BA, Schoepp DD, Palmer AJ, Bath CP, Sharpe EF, Wooley ML, Bufton HR, Kamboj RK, Tarnawa I, & Lodge D. (1996). Activity of 2,3-benzodiazepines at native rat and recombinant human glutamate receptors in vitro: Stereospecificity and selectivity profiles. Neuropharmacology 35: 1689–1702.

Bright DP, & Smart TG. (2013). Methods for recording and measuring tonic GABA_A_ receptor-mediated inhibition. Frontiers in Neural Circuits 7: 193.

Brymer KJ, Barnes JR, & Parsons MP. (2021). Entering a new era of quantifying glutamate clearance in health and disease. Journal of Neuroscience Research 99: 1598–1617.

Brymer KJ, Hurley EP, Barron JC, Mukherjee B, Barnes JR, Nafar F, & Parsons MP. (2023). Asymmetric dysregulation of glutamate dynamics across the synaptic cleft in a mouse model of Alzheimer’s disease. Acta Neuropathologica Communications 11: 27.

Campanelli F, Natale G, Marino G, Ghiglieri V, & Calabresi P. (2022). Striatal glutamatergic hyperactivity in Parkinson’s disease. Neurobiology of Disease 168: 105697.

Chen TW, Wardill TJ, Sun Y, Pulver SR, Renninger SL, Baohan A, Schreiter ER, Kerr RA, Orger MB, Jayaraman V, Looger LL, Svoboda K, & Kim DS. (2013). Ultrasensitive fluorescent proteins for imaging neuronal activity. Nature 499: 295–300.

Choi DW. (2020). Excitotoxicity: Still hammering the ischemic brain in 2020. Frontiers in Neuroscience 14: 579953.

Danbolt NC, Furness DN, & Zhou Y. (2016). Neuronal vs glial glutamate uptake: Resolving the conundrum. Neurochemistry International 98: 29–45.

Dawitz J, Kroon T, Hjorth JJJ, Mansvelder HD, & Meredith RM. (2020). Distinct synchronous network activity during the second postnatal week of medial entorhinal cortex development. Frontiers in Cellular Neuroscience 14: 91.

de Ceglia R, Ledonne A, Litvin DG, Lind BL, Carriero G, Latagliata EC, Bindocci E, Di Castro MA, Savtchouk I, Vitali I, Ranjak A, Congiu M, Canonica T, Wisden W, Harris K, Mameli M, Mercuri N, Telley L, & Volterra A. (2023). Specialized astrocytes mediate glutamatergic gliotransmission in the CNS. Nature 622: 120–129.

Diaz Verdugo C, Myren-Svelstad S, Aydin E, Van Hoeymissen E, Deneubourg C, Vanderhaeghe S, Vancraeynest J, Pelgrims R, Cosacak MI, Muto A, Kizil C, Kawakami K, Jurisch-Yaksi N, & Yaksi E. (2019). Glia-neuron interactions underlie state transitions to generalized seizures. Nature Communications 10: 3830.

Dimidschstein J, Chen Q, Tremblay R, Rogers SL, Saldi GA, Guo L, Xu Q, Liu R, Lu C, Chu J, Grimley JS, Krostag AR, Kaykas A, Avery MC, Rashid MS, Baek M, Jacob AL, Smith GB, Wilson DE, … Fishell G. (2016). A viral strategy for targeting and manipulating interneurons across vertebrate species. Nature Neuroscience 19: 1743–1749.

Dirnagl U, Iadecola C, & Moskowitz MA. (1999). Pathobiology of ischaemic stroke: An integrated view. Trends in Neuroscience 22: 391–397.

Dreier JP, & Reiffurth C. (2015). The stroke-migraine depolarization continuum. Neuron 86: 902– 922.

Dvorzhak A, Helassa N, Török K, Schmitz D, & Grantyn R. (2019). Single synapse indicators of impaired glutamate clearance derived from fast iGluu imaging of cortical afferents in the striatum of normal and huntington (Q175) mice. Journal of Neuroscience 39: 3970–3982.

Dzhala V, Khalilov I, Ben-Ari Y, & Khazipov R. (2001). Neuronal mechanisms of the anoxia-induced network oscillations in the rat hippocampus in vitro. Journal of Physiology 536: 521–531.

Edelstein AD, Tsuchida MA, Amodaj N, Pinkard H, Vale RD, & Stuurman N. (2014). Advanced methods of microscope control using μManager software. Journal of Biological Methods 1: e10.

Eikermann-Haerter K. (2021). Neuronal plumes initiate spreading depolarization, the electrophysiologic event driving migraine and stroke. Neuron 109: 563–565.

Eitelmann S, Everaerts K, Petersilie L, Rose CR, & Stephan J. (2023). Ca^2+^-dependent rapid uncoupling of astrocytes upon brief metabolic stress. Frontiers in Cellular Neuroscience 17: 1151608.

Eitelmann S, Stephan J, Everaerts K, Durry S, Pape N, Gerkau NJ, & Rose CR. (2022). Changes in astroglial K+ upon brief periods of energy deprivation in the mouse neocortex. International Journal of Molecular Sciences 23: 4836.

Enger R, Tang W, Vindedal GF, Jensen V, Helm PJ, Sprengel R, Looger LL, & Nagelhus EA. (2015). Dynamics of ionic shifts in cortical spreading depression. Cerebral Cortex 25: 4469–4476.

Epsztein J, Milh M, Id Bihi R, Jorquera I, Ben-Ari Y, Represa A, & Crepel V. (2006). Ongoing epileptiform activity in the post-ischemic hippocampus is associated with a permanent shift of the excitatory-inhibitory synaptic balance in CA3 pyramidal neurons. Journal of Neuroscience 26: 7082–7092.

Farsi Z, Walde M, Klementowicz AE, Paraskevopoulou F, & Woehler A. (2021). Single synapse glutamate imaging reveals multiple levels of release mode regulation in mammalian synapses. IScience 24: 101909.

Feuerbach D, Loetscher E, Neurdin S, & Koller M. (2010). Comparative pharmacology of the human NMDA-receptor subtypes R1-2A, R1-2B, R1-2C and R1-2D using an inducible expression system. European Journal of Pharmacology 637: 46–54.

Franke K, Berens P, Schubert T, Bethge M, Euler T, & Baden T. (2017). Inhibition decorrelates visual feature representations in the inner retina. Nature 542: 439–444.

Ge Y, Chen W, Axerio-Cilies P, & Wang YT. (2020). NMDARs in cell survival and death: Implications in stroke pathogenesis and treatment. Trends in Molecular Medicine 26: 533–551.

Gee CE, Ohmert I, Wiegert JS, & Oertner TG. (2017). Preparation of slice cultures from rodent hippocampus. Cold Spring Harbor Protocols 2017: 126–130.

Gerkau NJ, Rakers C, Durry S, Petzold GC, & Rose CR. (2018). Reverse NCX attenuates cellular sodium loading in metabolically compromised cortex. Cerebral Cortex 28: 4264–4280.

Hamilton NB, & Attwell D. (2010). Do astrocytes really exocytose neurotransmitters? Nature Reviews Neuroscience 11: 227–238.

Hao Y, Toulmé E, König B, Rosenmund C, & Plested AJ. (2023). Targeted sensors for glutamatergic neurotransmission. ELife 12: e84029.

Hefendehl JK, LeDue J, Ko RWY, Mahler J, Murphy TH, & MacVicar BA. (2016). Mapping synaptic glutamate transporter dysfunction in vivo to regions surrounding Aβ plaques by iGluSnFR two-photon imaging. Nature Communications 7: 13441.

Helassa N, Dürst CD, Coates C, Kerruth S, Arif U, Schulze C, Wiegert JS, Geeves M, Oertner TG, & Török K. (2018). Ultrafast glutamate sensors resolve high-frequency release at Schaffer collateral synapses. Proceedings of the National Academy of Sciences USA 115: 5594–5599.

Herde MK, Bohmbach K, Domingos C, Vana N, Komorowska-Müller JA, Passlick S, Schwarz I, Jackson CJ, Dietrich D, Schwarz MK, & Henneberger C. (2020). Local efficacy of glutamate uptake decreases with synapse size. Cell Reports 32: 108182.

Ibata K, Sun Q, & Turrigiano GG. (2008). Rapid synaptic scaling induced by changes in postsynaptic firing. Neuron 57: 819–826.

Imamura H, Huynh Nhat KP, Togawa H, Saito K, Iino R, Kato-Yamada Y, Nagai T, & Noji H. (2009). Visualization of ATP levels inside single living cells with fluorescence resonance energy transfer-based genetically encoded indicators. Proceedings of the National Academy of Sciences USA 106: 15651–15656.

Jensen TP, Zheng K, Cole N, Marvin JS, Looger LL, & Rusakov DA. (2019). Multiplex imaging relates quantal glutamate release to presynaptic Ca^2+^ homeostasis at multiple synapses in situ. Nature Communications 10: 1414.

Johnson HA, & Buonomano DV. (2007). Development and plasticity of spontaneous activity and up states in cortical organotypic slices. Journal of Neuroscience 27: 5915–5925.

Juzekaeva E, Gainutdinov A, Mukhtarov M, & Khazipov R. (2020). Reappraisal of anoxic spreading depolarization as a terminal event during oxygen–glucose deprivation in brain slices in vitro. Scientific Reports 10: 18970.

Kalia M, Meijer HGE, van Gils SA, van Putten MJAM, & Rose CR. (2021). Ion dynamics at the energy-deprived tripartite synapse. PLOS Computational Biology 17: e1009019.

Koch ET, Woodard CL, & Raymond LA. (2018). Direct assessment of presynaptic modulation of cortico-striatal glutamate release in a Huntington’s disease mouse model. Journal of Neurophysiology 120: 3077–3084.

Lerchundi R, Huang N, & Rose CR. (2020). Quantitative imaging of changes in astrocytic and neuronal adenosine triphosphate using two different variants of ATeam. Frontiers in Cellular Neuroscience 14: 80.

Lerchundi R, Kafitz KW, Winkler U, Färfers M, Hirrlinger J, & Rose CR. (2019a). FRET-based imaging of intracellular ATP in organotypic brain slices. Journal of Neuroscience Research 97: 933– 945.

Lerchundi R, Kafitz KW, Färfers M, Beyer F, Huang N, & Rose CR. (2019b). Imaging of intracellular ATP in organotypic tissue slices of the mouse brain using the FRET-based sensor ATeam1.03^YEMK^. Journal of Visualized Experiments 154: e60294.

Malarkey EB, & Parpura V. (2008). Mechanisms of glutamate release from astrocytes. Neurochemistry International 52: 142–154.

Marvin JS, Borghuis BG, Tian L, Cichon J, Harnett MT, Akerboom J, Gordus A, Renninger SL, Chen TW, Bargmann CI, Orger MB, Schreiter ER, Demb JB, Gan WB, Hires SA, & Looger LL. (2013). An optimized fluorescent probe for visualizing glutamate neurotransmission. Nature Methods 10: 162–170.

Marvin JS, Scholl B, Wilson DE, Podgorski K, Kazemipour A, Müller JA, Schoch S, Quiroz FJU, Rebola N, Bao H, Little JP, Tkachuk AN, Cai E, Hantman AW, Wang SS-H, DePiero VJ, Borghuis BG, Chapman ER, Dietrich D, … Looger LL. (2018). Stability, affinity, and chromatic variants of the glutamate sensor iGluSnFR. Nature Methods 15: 936–939.

Matthews EA, Sun W, McMahon SM, Doengi M, Halka L, Anders S, Müller JA, Steinlein P, Vana NS, van Dyk G, Pitsch J, Becker AJ, Pfeifer A, Kavalali ET, Lamprecht A, Henneberger C, Stein V, Schoch S, & Dietrich D. (2022). Optical analysis of glutamate spread in the neuropil. Cerebral Cortex 32: 3669–3689.

McGirr A, LeDue J, Chan AW, Xie Y, & Murphy TH. (2017). Cortical functional hyperconnectivity in a mouse model of depression and selective network effects of ketamine. Brain 140: 2210–2225.

Meade CA, Figueredo-Cardenas G, Fusco F, Nowak TS, Pulsinelli WA, & Reiner A. (2000). Transient global ischemia in rats yields striatal projection neuron and interneuron loss resembling that in Huntington’s disease. Experimental Neurology 166: 307–323.

Meyer J, Gerkau NJ, Kafitz KW, Patting M, Jolmes F, Henneberger C, & Rose CR. (2022). Rapid fluorescence lifetime imaging reveals that TRPV4 channels promote dysregulation of neuronal Na+ in ischemia. Journal of Neuroscience 42: 552–566.

Mohajerani MH, & Cherubini E. (2006). Role of giant depolarizing potentials in shaping synaptic currents in the developing hippocampus. Critical Reviews in Neurobiology 18: 13–23.

Okamoto K, Ishikawa T, Abe R, Ishikawa D, Kobayashi C, Mizunuma M, Norimoto H, Matsuki N, & Ikegaya Y. (2014). Ex vivo cultured neuronal networks emit in vivo-like spontaneous activity. The Journal of Physiological Sciences 64: 421–431.

Owji AP, Wang J, Kittredge A, Clark Z, Zhang Y, Hendrickson WA, & Yang T. (2022). Structures and gating mechanisms of human bestrophin anion channels. Nature Communications 13: 3836.

Pape N, & Rose CR. (2023). Activation of TRPV4 channels promotes the loss of cellular ATP in organotypic slices of the mouse neocortex exposed to chemical ischemia. The Journal of Physiology 601: 2975–2990.

Parker PD, Suryavanshi P, Melone M, Sawant-Pokam PA, Reinhart KM, Kaufmann D, Theriot JJ, Pugliese A, Conti F, Shuttleworth CW, Pietrobon D, & Brennan KC. (2021). Non-canonical glutamate signaling in a genetic model of migraine with aura. Neuron 109: 611–628.

Parsons MP, Vanni MP, Woodard CL, Kang R, Murphy TH, & Raymond LA. (2016). Real-time imaging of glutamate clearance reveals normal striatal uptake in Huntington disease mouse models. Nature Communications 7: 11251.

Passlick S, Rose CR, Petzold GC, & Henneberger C. (2021). Disruption of glutamate transport and homeostasis by acute metabolic stress. Frontiers in Cellular Neuroscience 15: 637784.

Pietrobon D, & Moskowitz MA. (2014). Chaos and commotion in the wake of cortical spreading depression and spreading depolarizations. Nature Reviews Neuroscience 15: 379–393.

Rakers C, Schmid M, & Petzold GC. (2017). TRPV4 channels contribute to calcium transients in astrocytes and neurons during peri-infarct depolarizations in a stroke model. Glia 65: 1550–1561.

Reiner A, & Levitz J. (2018). Glutamatergic signaling in the central nervous system: Ionotropic and metabotropic receptors in concert. Neuron 98: 1080–1098.

Risher WC, Croom D, & Kirov SA. (2012). Persistent astroglial swelling accompanies rapid reversible dendritic injury during stroke-induced spreading depolarizations. Glia 60: 1709–1720.

Rodríguez-Campuzano AG, & Ortega A. (2021). Glutamate transporters: Critical components of glutamatergic transmission. Neuropharmacology 192: 108602.

Rose CR, Felix L, Zeug A, Dietrich D, Reiner A, & Henneberger C. (2018). Astroglial glutamate signaling and uptake in the hippocampus. Frontiers in Molecular Neuroscience 10: 451.

Rusakov DA, & Stewart MG. (2021). Synaptic environment and extrasynaptic glutamate signals: The quest continues. Neuropharmacology 195: 108688.

Schindelin J, Arganda-Carreras I, Frise E, Kaynig V, Longair M, Pietzsch T, Preibisch S, Rueden C, Saalfeld S, Schmid B, Tinevez J-Y, White DJ, Hartenstein V, Eliceiri K, Tomancak P, & Cardona A. (2012). Fiji: An open-source platform for biological-image analysis. Nature Methods 9: 676–682.

Schreiner AE, Berlinger E, Langer J, Kafitz KW, & Rose CR. (2013). Lesion-induced alterations in astrocyte glutamate transporter expression and function in the hippocampus. ISRN Neurology 2013: 893605.

Shimamoto K, Sakai R, Takaoka K, Yumoto N, Nakajima T, Amara SG, & Shigeri Y. (2004). Characterization of novel L-threo-β-benzyloxyaspartate derivatives, potent blockers of the glutamate transporters. Molecular Pharmacology 65: 1008–1015.

Shimoda Y, Leite M, Graham RT, Marvin JS, Hasseman J, Kolb I, Looger LL, Magloire V, & Kullmann DM. (2024). Extracellular glutamate and GABA transients at the transition from interictal spiking to seizures. Brain 1011–1024.

Srivastava I, Vazquez-Juarez E, & Lindskog M. (2020). Reducing glutamate uptake in rat hippocampal slices enhances astrocytic membrane depolarization while down-regulating CA3– CA1 synaptic response. Frontiers in Synaptic Neuroscience 12: 37.

Stoppini L, Buchs P-A, & Muller D. (1991). A simple method for organotypic cultures of nervous tissue. Journal of Neuroscience Methods 37: 173–182.

Tarantini S, Tran CHT, Gordon GR, Ungvari Z, & Csiszar A. (2017). Impaired neurovascular coupling in aging and Alzheimer’s disease: Contribution of astrocyte dysfunction and endothelial impairment to cognitive decline. Experimental Gerontology 94: 52–58.

Tóth OM, Menyhárt Á, Frank R, Hantosi D, Farkas E, & Bari F. (2020). Tissue acidosis associated with ischemic stroke to guide neuroprotective drug delivery. Biology 9: 460.

Tsukada S, Iino M, Takayasu Y, Shimamoto K, & Ozawa S. (2005). Effects of a novel glutamate transporter blocker, (2S, 3S)-3-{3-[4-(trifluoromethyl)benzoylamino]benzyloxy}aspartate (TFB-TBOA), on activities of hippocampal neurons. Neuropharmacology 48: 479–491.

Unichenko P, Yang J-W, Luhmann HJ, & Kirischuk S. (2015). Glutamatergic system controls synchronization of spontaneous neuronal activity in the murine neonatal entorhinal cortex. Pflügers Archiv - European Journal of Physiology 467: 1565–1575.

van Putten MJAM, Fahlke C, Kafitz KW, Hofmeijer J, & Rose CR. (2021). Dysregulation of astrocyte ion homeostasis and its relevance for stroke-induced brain damage. International Journal of Molecular Sciences 22: 5679.

Wu Z, Lin D, & Li Y. (2022). Pushing the frontiers: Tools for monitoring neurotransmitters and neuromodulators. Nature Reviews Neuroscience 23: 257–274.

Yang J, Vitery M del C, Chen J, Osei-Owusu J, Chu J, & Qiu Z. (2019). Glutamate-releasing SWELL1 channel in astrocytes modulates synaptic transmission and promotes brain damage in stroke. Neuron 102: 813–827.

Yu SP, Jiang MQ, Shim SS, Pourkhodadad S, & Wei L. (2023). Extrasynaptic NMDA receptors in acute and chronic excitotoxicity: Implications for preventive treatments of ischemic stroke and late-onset Alzheimer’s disease. Molecular Neurodegeneration 18: 43.

Zott B, Simon MM, Hong W, Unger F, Chen-Engerer H-J, Frosch MP, Sakmann B, Walsh DM, & Konnerth A. (2019). A vicious cycle of β amyloid–dependent neuronal hyperactivation. Science 365: 559–565.

