## Supplementary Information for "Atypical plume-like events drive glutamate accumulation in metabolic stress conditions"

This document includes:

**Supplementary Figures S1-22** p. 2
**Supplementary Movie Information** p. 31
**Supplementary Methods** p. 32
**Supplementary References** p. 41

### Supplementary Figures S1-S22

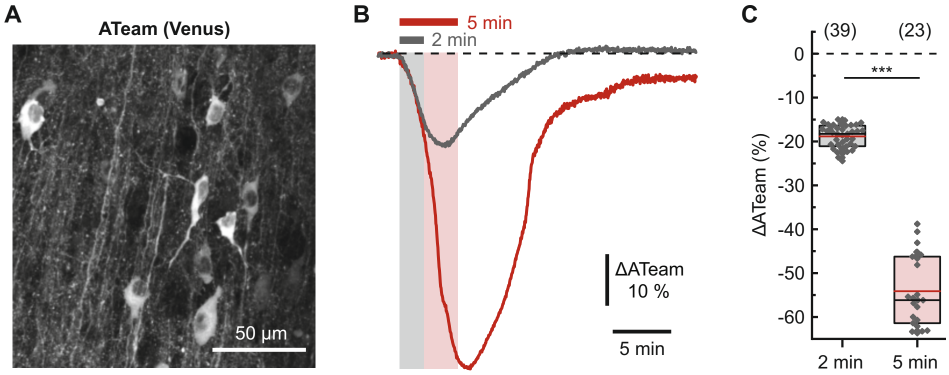


**Figure S1. Changes in neuronal ATP levels during chemical ischemia. (A)** Confocal image of the Venus fluorescence of the FRET-based ATP sensor ATeam1.03$^{YEMK}$ driven by a hSyn1 promotor in an organotypic cultured slice of the mouse somatosensory cortex. **(B)** Changes in ATeam fluorescence ratio in two individual neurons upon applying chemical ischemia for 2 min (grey trace) or 5 min (red trace) as indicated. **(C)** Maximum decrease in ATeam fluorescence ratio induced by chemical ischemia. Numbers in parentheses represent the number of analyzed neurons in a total of 4 slices (2 min) and 3 slices (5 min). Boxes show medians and 25-75% percentiles, red bars indicate the mean. Data were statistically analyzed by one-way ANOVA and a *post hoc* Bonferroni test, ***$p$ <0.001. For details see **Supplementary Methods**.

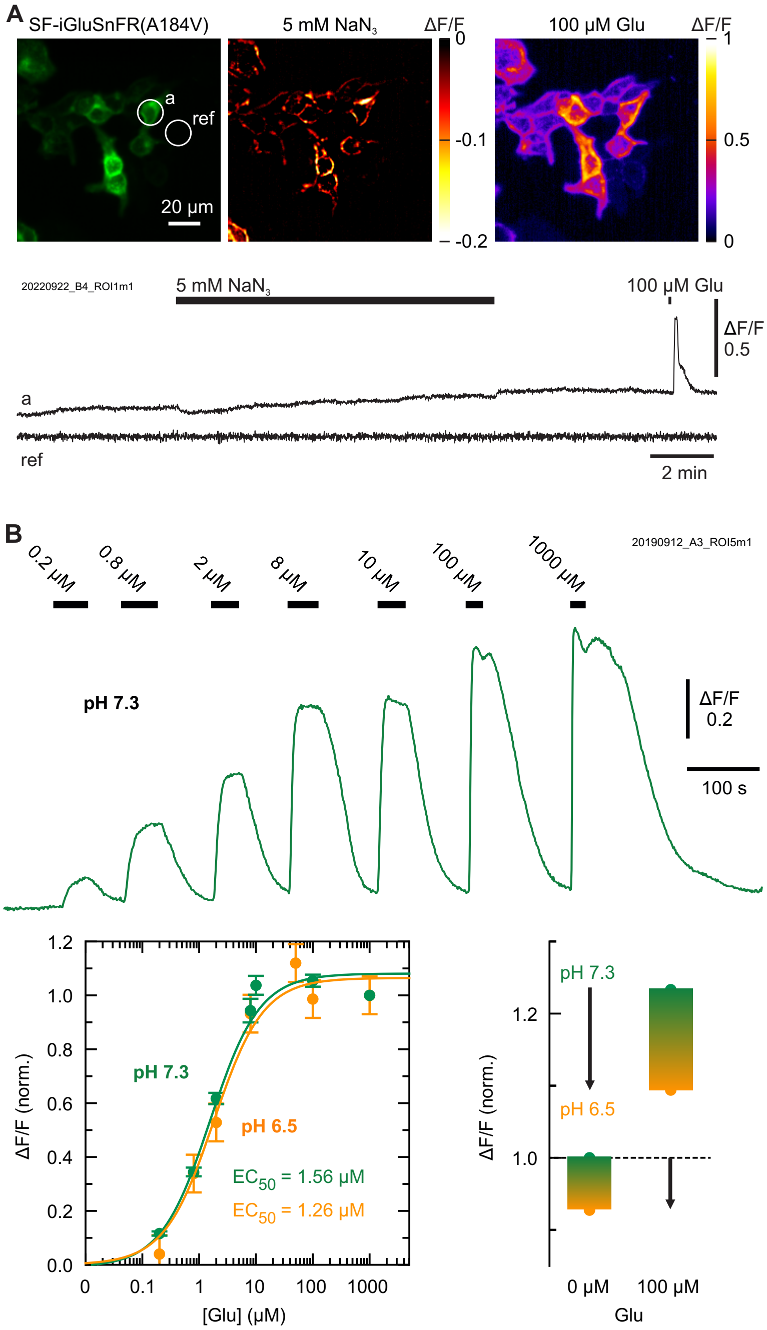


**Figure S2. Effect of azide and pH on SF-iGluSnFR(A184V) fluorescence in HEK cells. (A)** Fluorescence of HEK cells expressing SF-iGluSnFR(A184V) (left), rel. fluorescence decrease ΔF/F upon addition of 5 mM sodium azide (center) and ΔF/F increase upon addition of 100 µM glutamate (right). Bottom: Addition of azide causes a small, stable and reversible decrease of ΔF/F in region a (cell) but not in a control region (background, ref). **(B)** Measurement of a glutamate dose-response curve at pH 7.3 in HEK cells (top). $EC_{50}$ values were determined by fitting the Hill equation (solid lines, $h$ = 1) to normalized ΔF/F values (mean ± s.d.) from 4 experiments at pH 7.3 and pH 6.5, respectively (bottom). The affinity remains unchanged. Right: Acidification from pH 7.3 to pH 6.5 reduced the signal ΔF/F both in the ligand-free state (-7%) and glutamate-bound state (-14%). For details see **Supplementary Methods**.

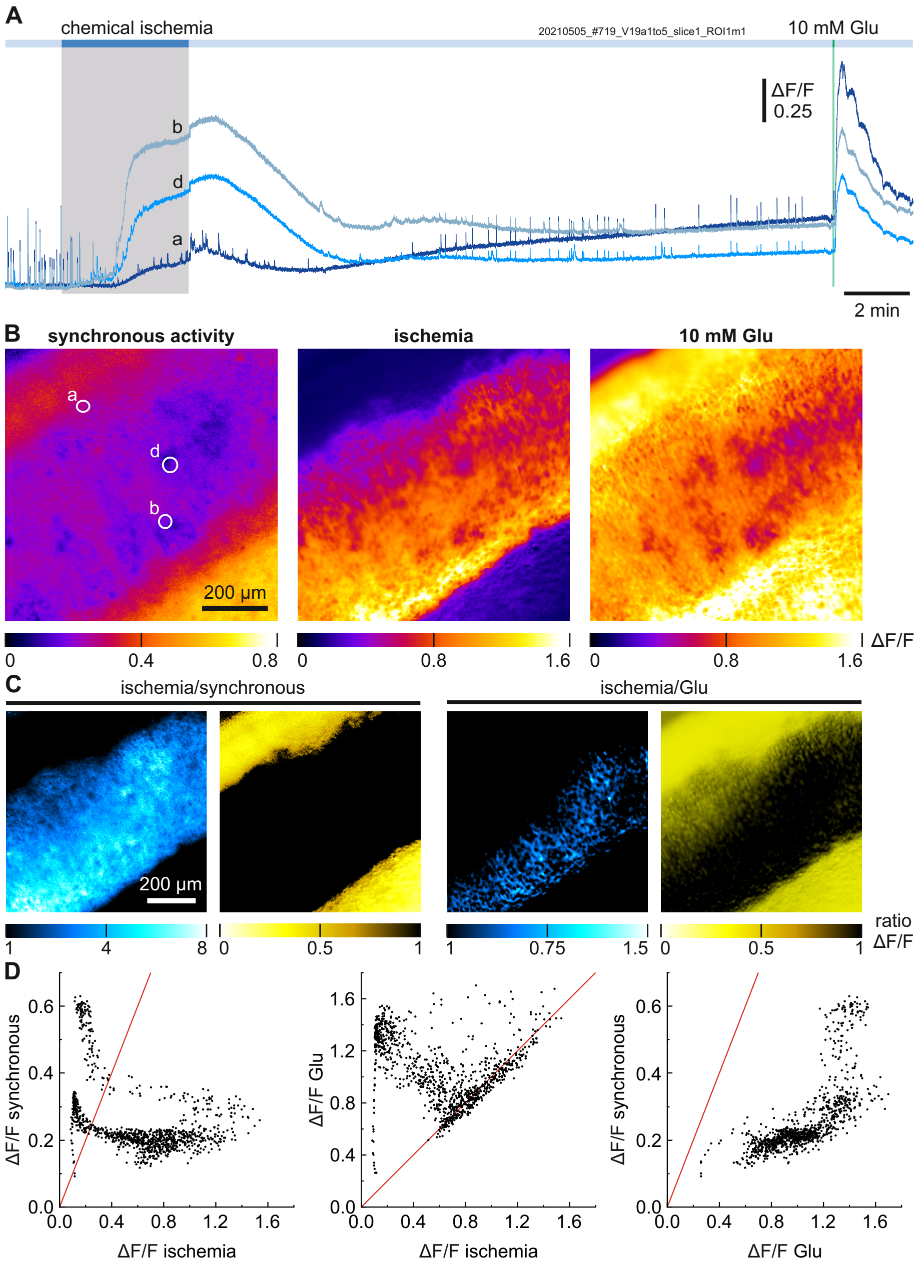


**Figure S3. SF-iGluSnFR(A184V) fluorescence increase upon inducing chemical ischemia and bath application of 10 mM glutamate. (A)** ΔF/F trace of the slice regions indicated in (B) from the same example as shown in **Fig. 1A**. At the end of the recording 10 mM glutamate were added to the bath, as indicated. **(B)** ΔF/F images of synchronous activity, ischemia, and 10 mM glutamate show regional differences in signal intensity. In all three cases, F from the beginning of the recording ($F_0$, 200 frames) was used for normalization. For details see **Supplementary Methods**. **(C)** ΔF/F ratio images (based on (B)) show the regions in which the signals were higher during the ischemic glutamate accumulation (blue) or during synchronous activity or 10 mM exogenous glutamate application (yellow). **(D)** Pixelwise comparison between the fluorescence intensity of the different signals. Left: Glutamate signals during synchronous activity and ischemia are not correlated. Centre: Some regions reach saturation during the ischemic period (same signal change as with 10 mM glutamate). Right: Signals from synchronous activity stay below the signals from 10 mM glutamate. Further quantitative interpretation is complicated by the SF-iGluSnFR(A184V) baseline run-up (see **Fig. S8**) and the reduction of fluorescence changes (ΔF/F values).

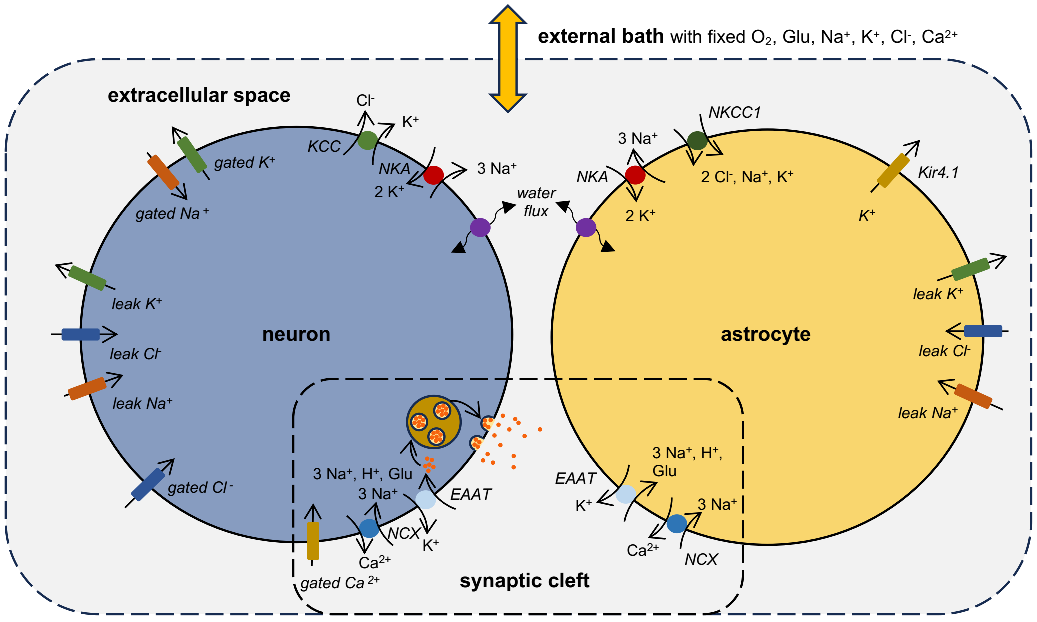


**Figure S4. Model used for computational assessment of key physiological parameters and bath coupling.** The model, which is an extension of Kalia *et al.* [Kalia *et al.*, 2021], comprises different compartments, including a neuron and an astrocyte with a somatic and synaptic compartment each, a synaptic cleft, extracellular space and an external bath. Calcium and glutamate are restricted to the synaptic compartments and synaptic cleft. All active and passive transport processes incorporated into the model are indicated by arrows. The external bath has constant concentrations of all species and equilibrates the $Na^+$, $K^+$ and $Cl^-$ concentrations in the extracellular space, and $Ca^{2+}$ and glutamate in the synaptic cleft. Only the bath oxygen level is reduced to simulate transient chemical ischemia. For details and parameterization see **Supplementary Methods**.

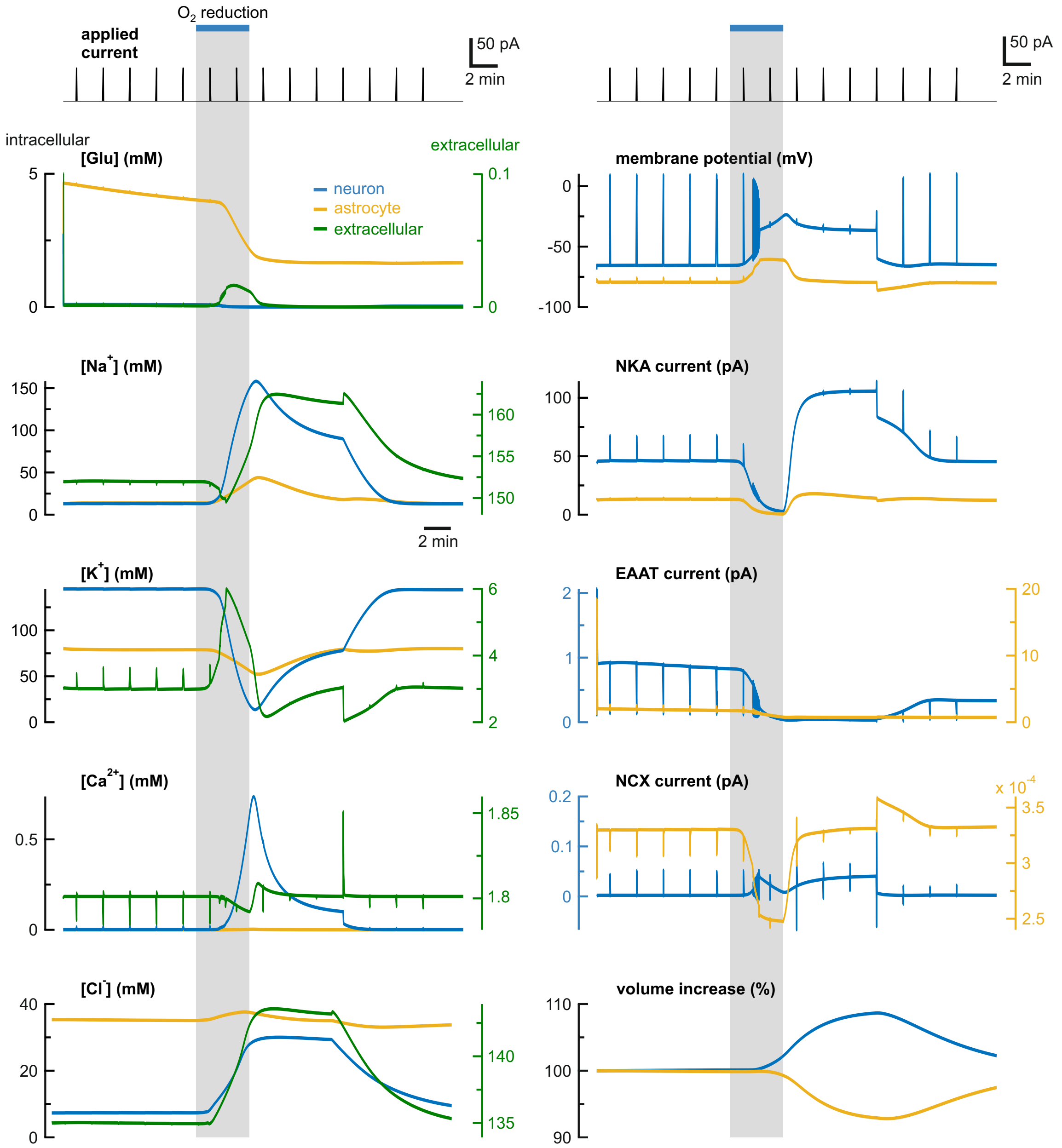


**Figure S5. Computational assessment of key physiological parameters during network activity and chemical ischemia. (Top panels)** Current injections (every 2 min) and oxygen reduction to 10% (for 4 min) used to simulate network activity and ischemia, respectively. **(Lower panels)** Resulting changes in ion concentrations, membrane potential, cellular volume and transporter currents. Effects on neurons are shown in blue, on extracellular space in green, and on astrocytes in yellow. Data are shown for medium bath coupling ($B = 1*10^{-3}$ $ms^{-1}$). For details see **Supplementary Methods**.

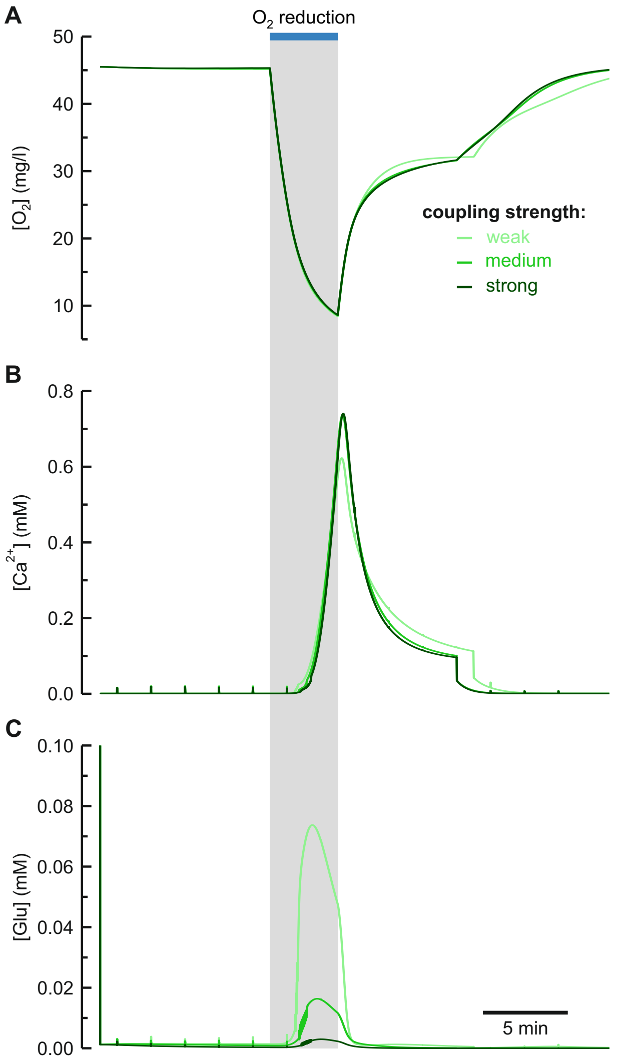


**Figure S6. Computational assessment of bath coupling.** The influence of different degrees of bath coupling (weak, medium, strong indicated by color) is shown for selected components (see also **Fig. S5**). **(A)** The net oxygen reduction (ischemia) in the extracellular space is similar in all cases, but recovery is different. **(B)** Strong neuronal $Ca^{2+}$ accumulation is seen in all conditions. **(C)** Extracellular glutamate accumulation is more pronounced for weaker bath coupling. For details see **Supplementary Methods**.

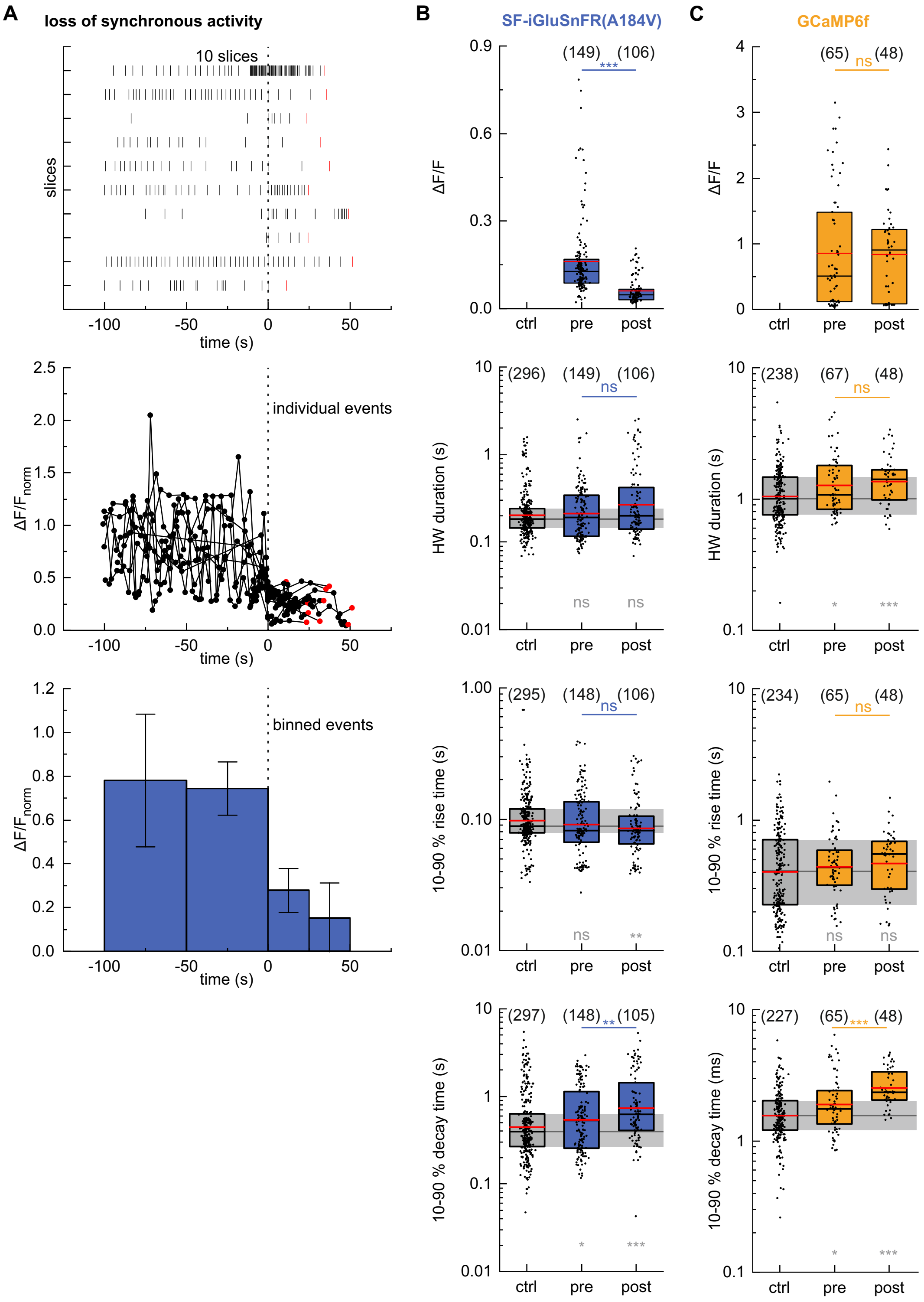


**Figure S7. Effects of chemical ischemia on synchronous activity detected by SF-iGluSnFR(A184V) and GCamP6f. (A)** SF-iGluSnFR(A184V) reports on synchronous activity, which ceased in all regions upon inducing chemical ischemia (cf. **Fig. 1A-D**). More detailed analysis showed a major decrease in ΔF/F in the 50 s period preceding the last detectable event (red). Some slices (3 of 10) showed a frequency increase in this period (top, raster plot). The event sequences of the 10 slices were aligned to the timepoint after which ΔF/F remained <50% of the average ΔF/F (center, time 0 s) and were then binned accordingly (bottom). One other slice was excluded from this analysis due to weak signal of synchronous activity. Bars with error bars show means ± s.d.. For details see **Supplementary Methods**. **(B,C)** Analysis of synchronous events before (pre) and after the ischemic period, as detected by hSyn1 SF-iGluSnFR(A184V) imaging (B) and hSyn1 GCaMP6f imaging (C). The imaging of synchronous events with SF-iGluSnFR was partly limited by the imaging frame rate (20 fps; for faster imaging see **Fig. S11**). The half-width duration (HWD) and 10-90% rise and decay times are compared to the respective 40 min control recordings (ctrl column and grey bar; cf. **Fig. S8**). Kruskal-Wallis tests were used to test for differences between the different time windows, followed by Dunn's test (blue/orange symbols). Additionally, Mann-Whitney *U* tests were used for pair-wise comparison of individual time windows to control (grey symbols). Numbers in parentheses give the number of analyzed events from 10 slices (SF-iGluSnFR) and 8 slices (GCamP6f). Boxes show medians and 25-75% percentiles, red bars indicate the mean. ns not significant, *$p$ <0.05, **$p$ <0.01, ***$p$ <0.001.

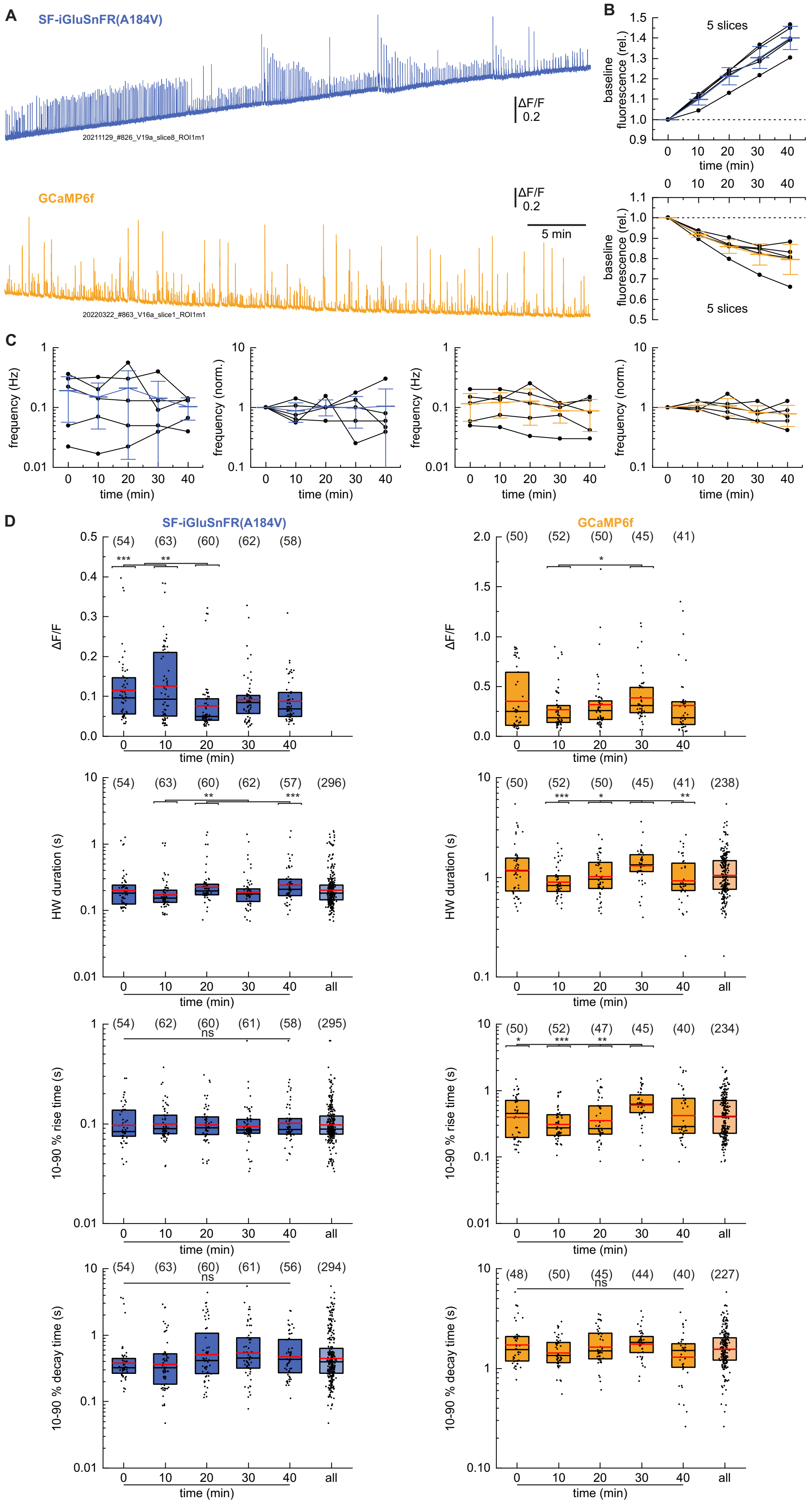
A
SF-iGluSnFR(A184V)
ΔF/F 0.2
GCaMP6f
5 min
B
5 slices
baseline fluorescence (rel.)
time (min)
C
frequency (Hz)
frequency (norm.)
time (min)
D
ΔF/F
HW duration (s)
10-90 % rise time (s)
10-90 % decay time (s)
time (min)
ns

**Figure S8. Baseline changes and synchronous activity without inducing chemical ischemia.** In these control recordings no chemical ischemia was induced, otherwise the conditions were identical. For plume analysis of these recordings see **Fig. S14**. **(A)** ∆F/F traces of continuous 50 min hSyn1 SF-iGluSnFR(A184V) (top) and hSyn1 GCaMP6f imaging (after 15-20 min preequilibration). **(B)** Relative changes in baseline fluorescence intensity. The fluorescence of SF-iGluSnFR(A184V) showed a continuous run-up, which was also seen in all other experimental conditions. Bars with error bars show means ± s.d.. **(C)** Absolute and normalized frequencies of synchronous events. Bars with error bars show means ± s.d.. **(D)** Analysis of ∆F/F, half-width duration (HWD), and 10-90% rise and decay times for individual events over a 40 min timecourse (25-300 s analysis windows). Kruskal-Wallis tests were used to test for differences between the different time windows, followed by Dunn's test. Despite some statistically significant differences, the datasets (with exception of ∆F/F) showed no systematic biases and were merged into 'all' groups to serve for control comparisons. Numbers in parentheses give the number of analyzed events from 5 slices for each, SF-iGluSnFR and GCaMP6f. Boxes show medians and 25-75% percentiles, red bars indicate the mean. ns not significant, $^{*}p < 0.05$, $^{**}p < 0.01$, $^{***}p < 0.001$.

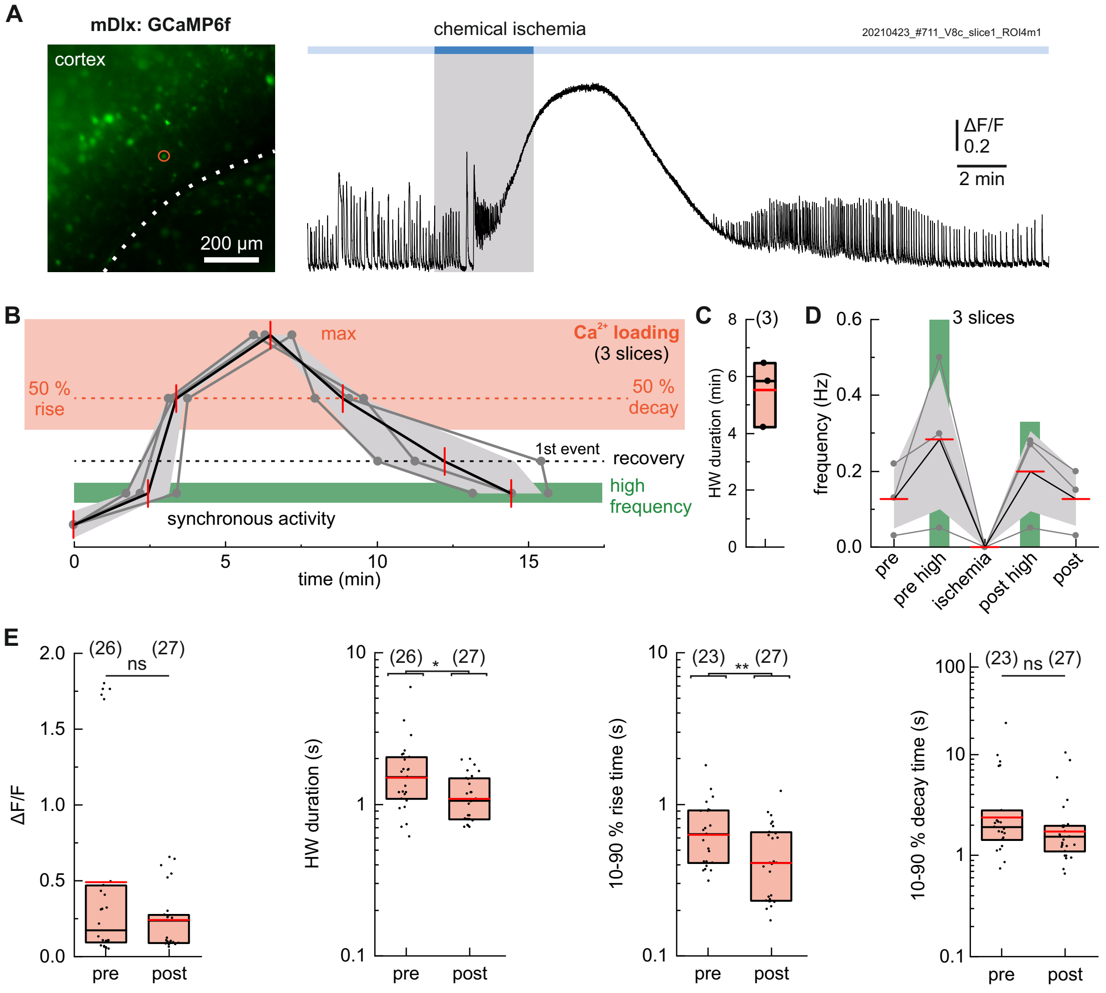


**Figure S9. Ca$^{2+}$ responses of the mDlx interneuron subpopulation only. (A)** Expression of GCaMP6f under control of the interneuron-specific mDlx enhancer and Ca$^{2+}$ imaging during chemical ischemia. Similar to hSyn1-driven GCaMP6f expression (**Fig. 1E-H**), increased synchronous activity was observed before and after chemical ischemia, as well as Ca$^{2+}$ loading during ischemia. **(B)** Timepoints of Ca$^{2+}$ loading (50% rise, maximum and 50% decay times; top), high frequency periods and recovery of synchronous activity (bottom) for 3 slices. **(C)** Half-width duration of interneuron Ca$^{2+}$ loading. **(D)** Frequency of synchronous Ca$^{2+}$ activity (60-300 s analysis windows). **(E)** Analysis of ΔF/F, half-width duration (HWD), and 10-90% rise and decay times of synchronous Ca$^{2+}$ events before and after chemical ischemia (60 s or 180 s analysis windows). Mann-Whitney *U* tests were used for pair-wise comparison. Numbers in parentheses give the number of analyzed events. Individual data points are shown in grey, mean values in red and s.d. as shaded areas. Boxes show medians and 25-75% percentiles, red bars indicate the mean. ns not significant, *$p$ <0.05, **$p$ <0.01, ***$p$ <0.001.

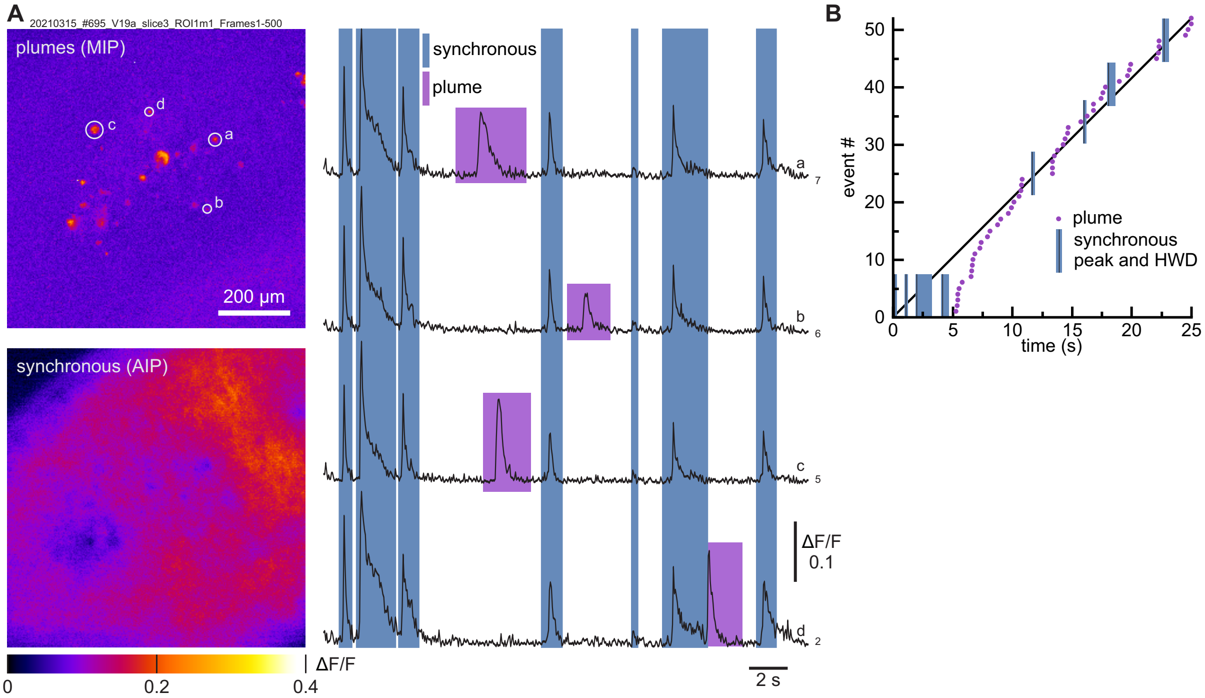


**Figure S10. SF-iGluSnFR(A184V) imaging shows synchronous activity and local glutamate plumes.** Another example slice showing plumes and synchronous activity in the pre-ischemic period. In contrast to the example in **Fig. 2A**, bursting behavior is seen for synchronous activity. **(A)** Top: Glutamate plumes in a maximal intensity projection (MIP; 25 s, frames with synchronous activity were excluded). Bottom: Fluorescence changes during synchronous activity (average intensity projection (AIP) of frames at the event maxima). Right: ΔF/F traces of four selected regions (circled). Synchronous events (blue boxes) and local plumes (purple boxes) can be clearly distinguished. **(B)** Timepoints of plume occurrence (purple) and phases of synchronous activity (blue) during the 25 s imaging period.

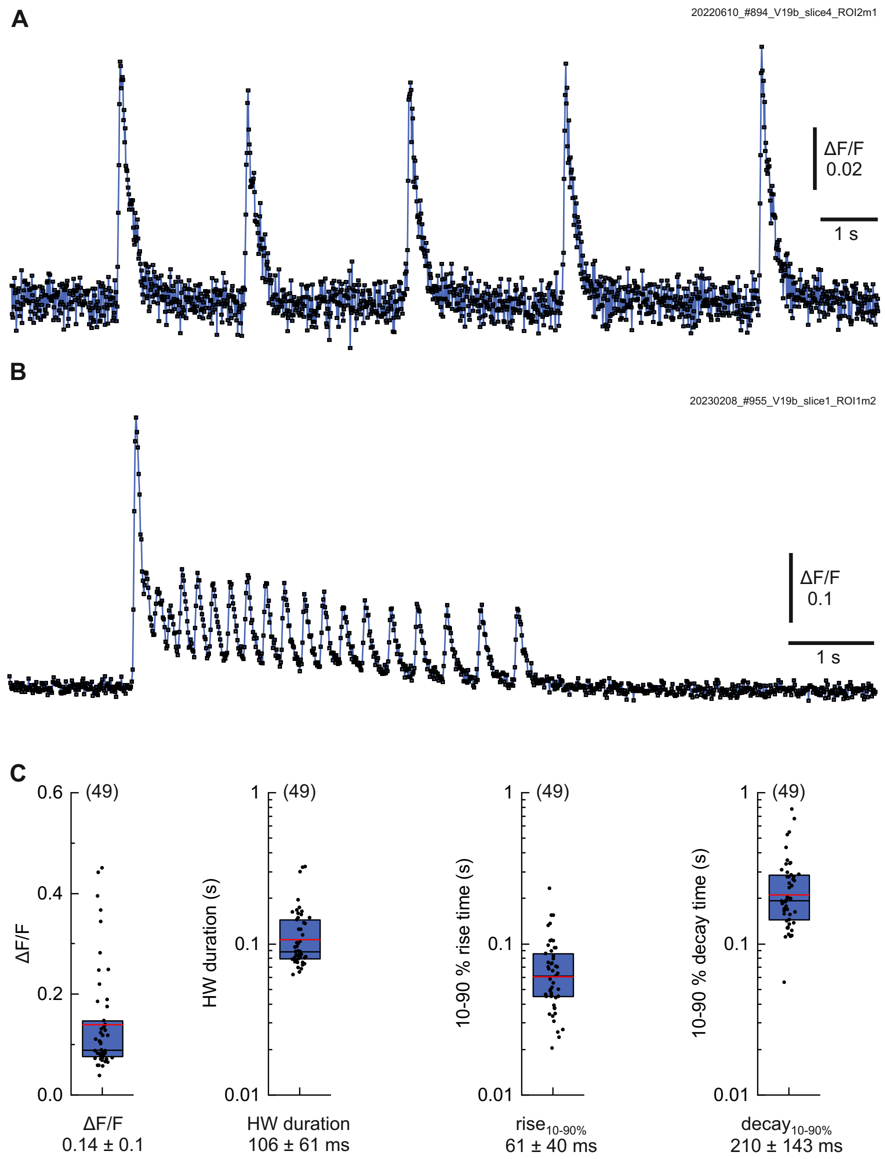


**Figure S11. Synchronous activity observed with fast SF-iGluSnFR(A184V) imaging.** Synchronous events from a pre-ischemic period recorded with 99 fps frame rate (vs 20 fps for standard recordings). **(A)** ΔF/F trace with points showing the values from individual frames. **(B)** ΔF/F trace from an example slice that showed bursting behavior. **(C)** Analysis of ΔF/F, half-width duration (HWD), and 10-90% rise and decay times for 49 individual events from 5 slices. Boxes show medians and 25-75% percentiles with red bars indicating the mean.

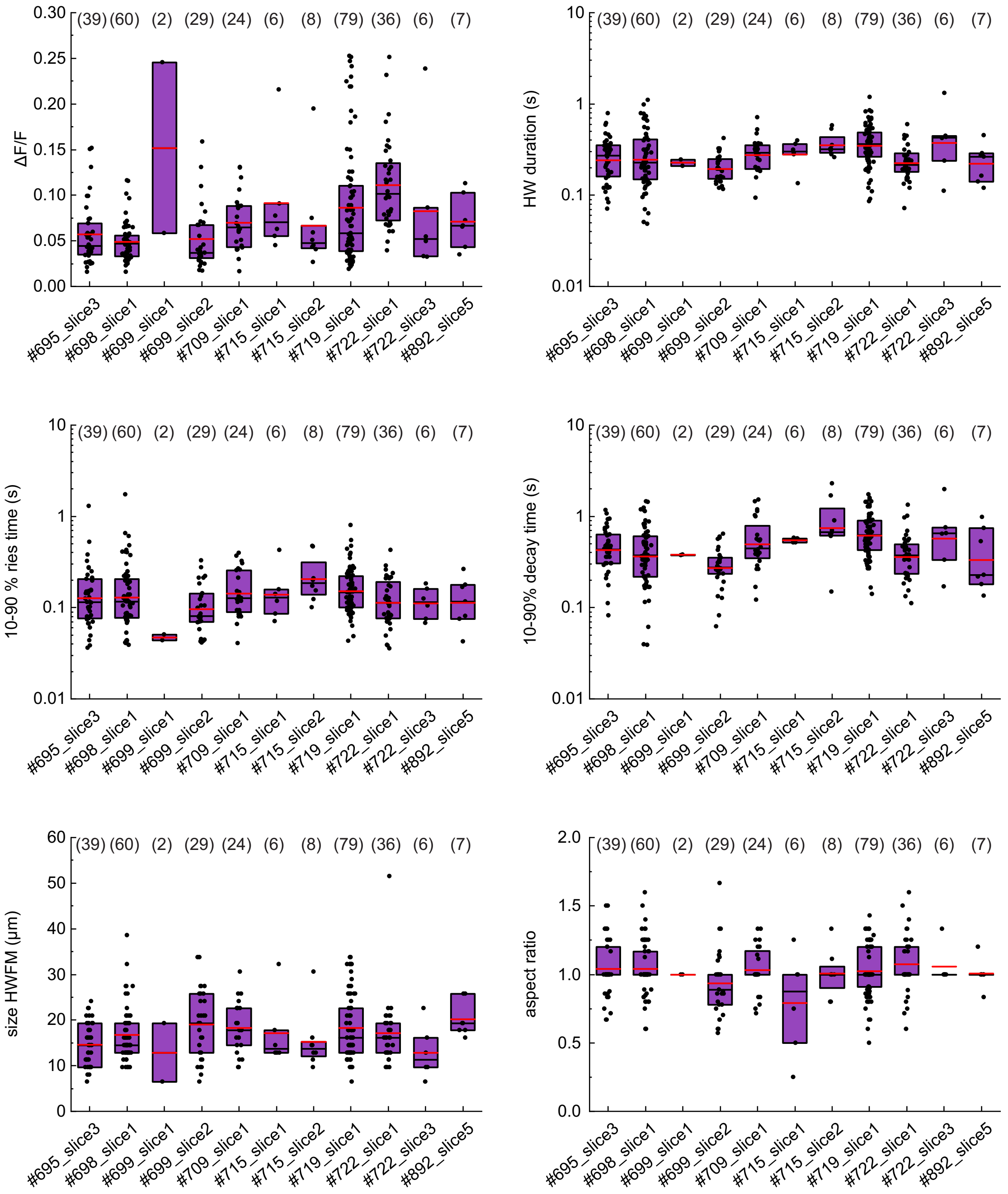


**Figure S12. Comparison of plume parameters obtained from different slice cultures.** Plume parameters obtained from SF-iGluSnFR(A184V) imaging of 25 s pre-ischemic time windows separated by slice (11 slices which showed plumes pre-ischemia; for combined data see **Fig. 2B** and **Fig. 3B**). Numbers in parentheses give the numbers of analyzed plumes. All plumes for which all parameters could be obtained were included. Boxes show medians and 25-75% percentiles, red bars indicate the mean.

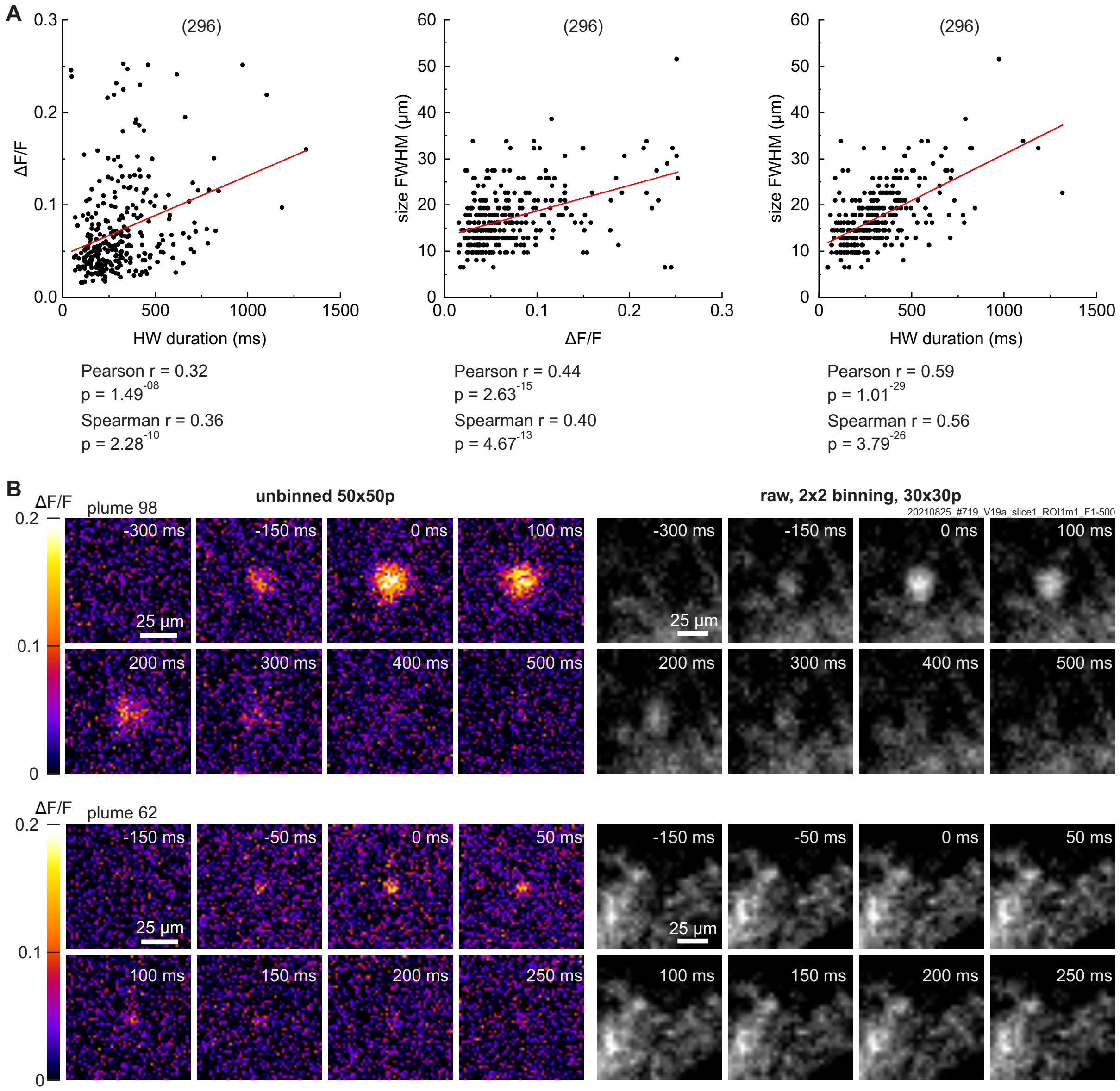


**Figure S13. Correlations of plume characteristics and data processing. (A)** Correlation analysis of parameters from 296 pre-ischemic plumes: Plume fluorescence intensity ΔF/F, half-width duration (HWD) and full-width at half maximum (FWHM) (data from 11 slices, see **Fig. 2** and **Fig. 3**). Pearson and Spearman tests were performed. The red lines show the linear correlation obtained by the Pearson test. **(B)** Effects of data processing illustrated for two example plumes (see **Fig. 3C,F**). Left: ΔF/F maps of plumes from unbinned movies (512 x 512 pixels). The plume regions were previously identified in 2 x 2 binned movies. Right: Unprocessed, raw intensity images (F) of plumes from binned movies (256 x 256 pixels).

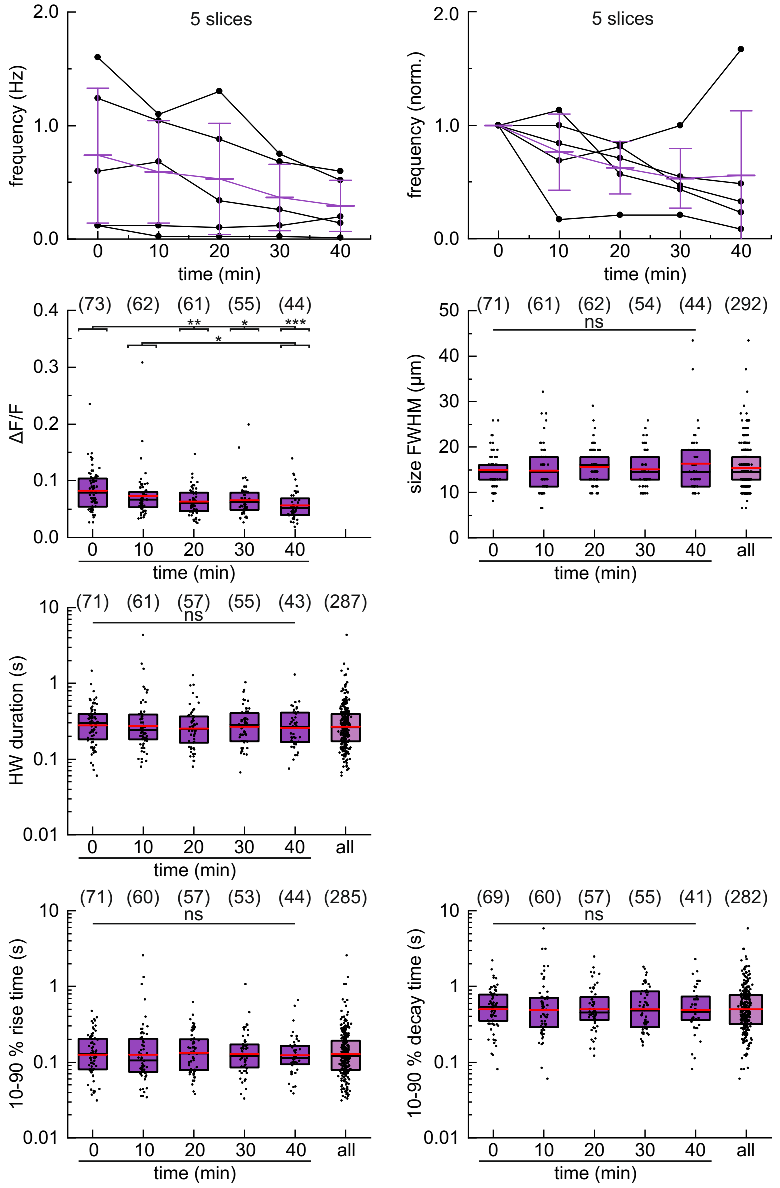


**Figure S14. Plume parameters over 50 min imaging without inducing chemical ischemia.** Data from hSyn1 SF-iGluSnFR(A184V) control recordings (5 slices; see also **Fig. S8**) at the indicated timepoints. Top: Absolute and normalized frequencies of detected plumes (10-300 s analysis windows). Bars with error bars show means ± s.d.. Below: Analysis of ΔF/F, full-width at half maximum (FWHM) size, half-width (HW) duration, and 10-90% rise and decay times of individual plumes (10-200 s analysis windows). Kruskal-Wallis tests were used to test for differences between the different time windows, followed by Dunn's test. All parameters, with exception of ΔF/F, showed no statistically significant differences between the different timepoints and were merged into 'all' groups for control comparisons. Numbers in parentheses give the number of analyzed events from 5 slices. Boxes show medians and 25-75% percentiles with red bars indicating the mean. ns not significant, **p* <0.05, ***p* <0.01, ****p* <0.001.

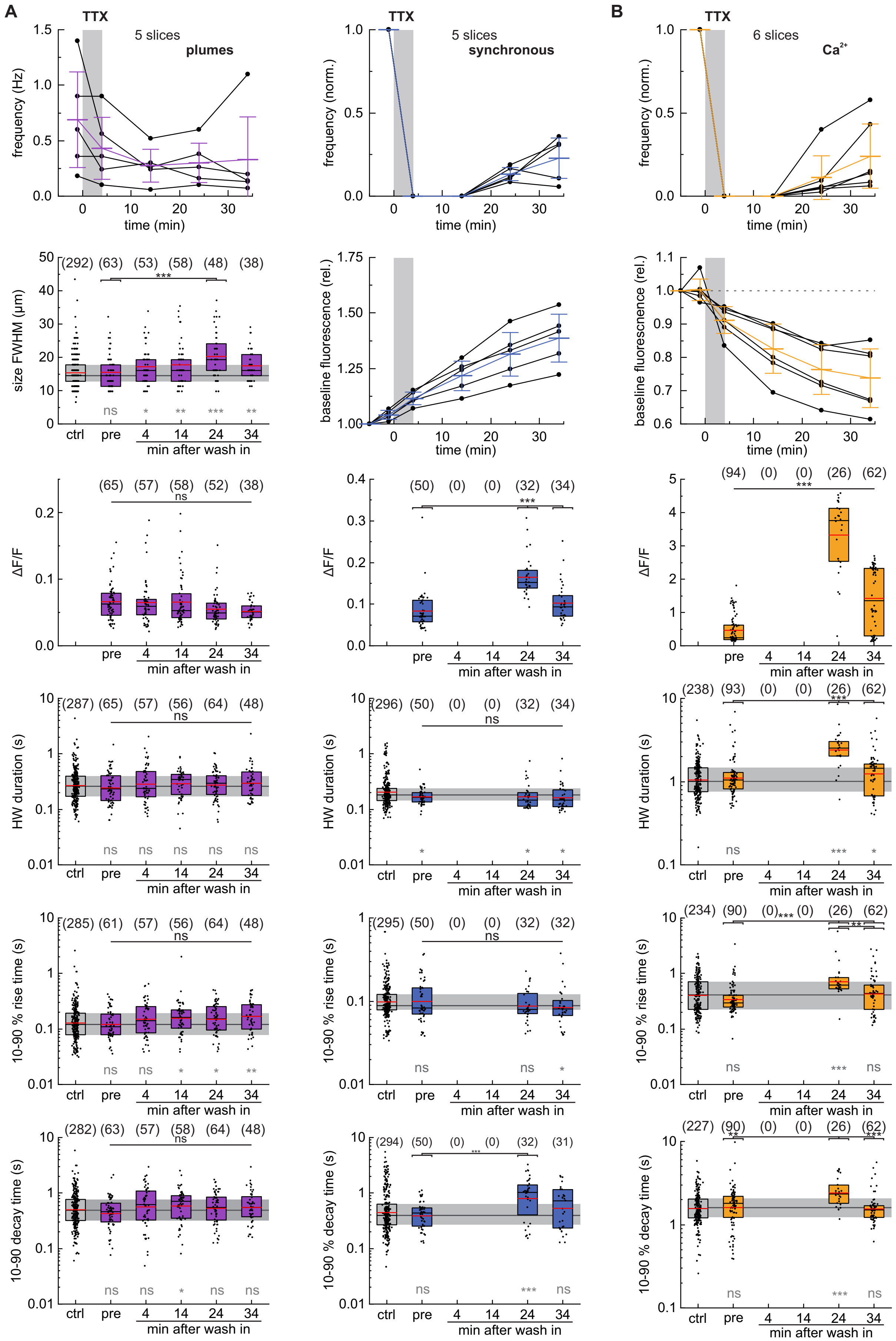
A
B
TTX
5 slices
plumes
synchronous
6 slices
$Ca^{2+}$
frequency (Hz)
frequency (norm.)
time (min)
size FWHM (µm)
baseline fluorescence (rel.)
ΔF/F
HW duration (s)
10-90 % rise time (s)
10-90 % decay time (s)
ctrl
pre
min after wash in

**Figure S15. TTX effects on plumes and synchronous activity.** TTX (0.2 µM) did not affect plumes but fully suppressed synchronous activity. After wash-out the signal amplitudes $\Delta F/F$ of synchronous activity were strongly increased. **(A)** Parameter summary of TTX effects on glutamate plumes and synchronous events detected with SF-iGluSnFR(A184V) (5 slices). The investigation of synchronous glutamate events was in part limited by the imaging frame rate (20 fps). An example experiment and normalized plume frequencies are shown in **Fig. 4A**. **(B)** Parameter summary of TTX effects on synchronous activity detected with GCaMP6f (6 slices). Where possible, data were compared to the respective 40 min control recordings (ctrl column and grey bar; cf. **Fig. S8** and **Fig. S14**). Kruskal-Wallis tests were used to test for differences between the different time windows, followed by Dunn's test (black symbols). Additionally, Mann-Whitney *U* tests were used for pair-wise comparison of individual time windows to control (grey symbols). Numbers in parentheses give the number of analyzed events. Bars with error bars show means ± s.d.. Boxes show medians and 25-75% percentiles, red bars indicate the mean. ns not significant, *$p$ <0.05, **$p$ <0.01, ***$p$ <0.001.

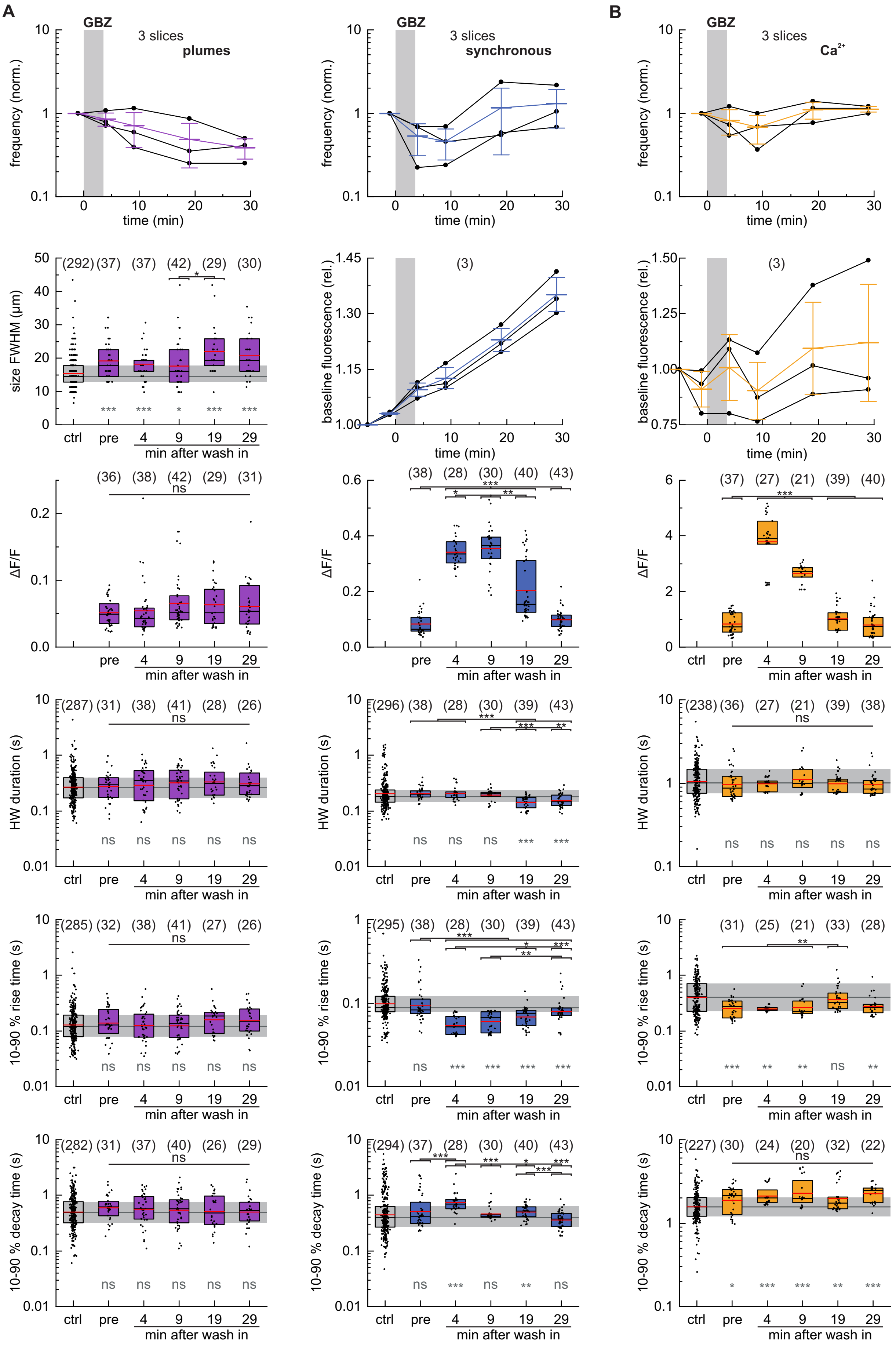
A
B
GBZ
3 slices
plumes
synchronous
Ca2+
frequency (norm.)
time (min)
size FWHM (µm)
baseline fluorescence (rel.)
ΔF/F
HW duration (s)
10-90 % rise time (s)
10-90 % decay time (s)
ctrl
pre
4
9
19
29
min after wash in
ns
(292) (37) (37) (42) (29) (30)
(36) (38) (42) (29) (31)
(287) (31) (38) (41) (28) (26)
(285) (32) (38) (41) (27) (26)
(282) (31) (37) (40) (26) (29)
(3)
(38) (28) (30) (40) (43)
(296) (38) (28) (30) (39) (43)
(295) (38) (28) (30) (39) (43)
(294) (37) (28) (30) (40) (43)
(37) (27) (21) (39) (40)
(238) (36) (27) (21) (39) (38)
(31) (25) (21) (33) (28)
(227) (30) (24) (20) (32) (22)

**Figure S16. Gabazine effects on plumes and synchronous activity.** GBZ (3 µM) affected synchronous event amplitudes but did not strongly affect plumes. **(A)** Parameter summary of glutamate plumes and synchronous activity detected with SF-iGluSnFR (3 slices). The investigation of synchronous glutamate events was partly limited by the imaging frame rate (20 fps). For an example experiment and normalized plume frequencies see **Fig. 4B**. **(B)** Parameter summary of synchronous activity detected with GCaMP6f (3 slices). Where possible, data were compared to the respective 40 min control recordings (ctrl column and grey bar; cf. **Fig. S8** and **Fig. S14**). Kruskal-Wallis tests were used to test for differences between the different time windows, followed by Dunn's test (black symbols). Additionally, Mann-Whitney *U* tests were used for pair-wise comparisons to control (grey symbols). Numbers in parentheses give the number of analyzed events. Bars with error bars show means ± s.d.. Boxes show medians and 25-75% percentiles, bars indicate the mean. ns not significant, *$p$ <0.05, **$p$ <0.01, ***$p$ <0.001.

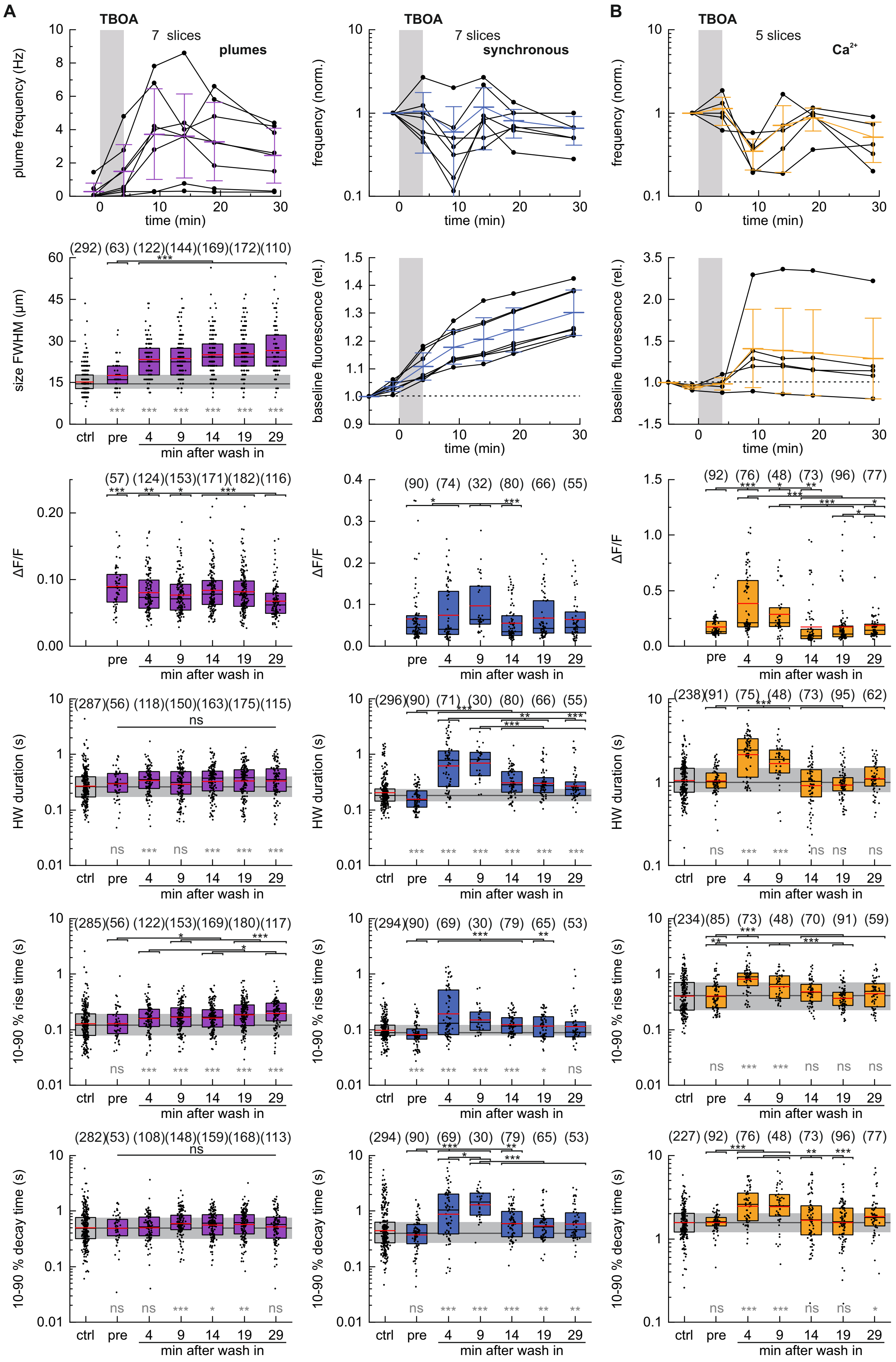
A
B
TBOA
7 slices
plumes
synchronous
5 slices
Ca2+
plume frequency (Hz)
frequency (norm.)
time (min)
size FWHM (µm)
baseline fluorescence (rel.)
ctrl
pre
min after wash in
ΔF/F
HW duration (s)
10-90 % rise time (s)
10-90 % decay time (s)
ns

**Figure S17. TFB-TBOA effects on plumes and synchronous activity.** TFB-TBOA (1 µM) increased plume frequency and had sustained effects on plume size and rise times. It also affected synchronous events. **(A)** Parameter summary of TFB-TBOA effects on glutamate plumes and synchronous events detected with SF-iGluSnFR(A184V) (7 slices). The investigation of synchronous glutamate events is in part limited by the imaging frame rate (20 fps). An example experiment and normalized plume frequencies are shown in **Fig. 5A**. **(B)** Parameter summary of TFB-TBOA effects on synchronous activity detected with GCaMP6f (5 slices). Where possible, data were compared to the respective 40 min control recordings (ctrl column and grey bar; cf. **Fig. S8** and **Fig. S14**). In most slices TBOA also caused an increase in the GCaMP6f baseline signal. Kruskal-Wallis tests were used to test for differences between the different time windows, followed by Dunn's test (black symbols). Additionally, Mann-Whitney *U* tests were used for pair-wise comparisons to control (grey symbols). Numbers in parentheses give the number of analyzed events. Bars with error bars show means ± s.d.. Boxes show medians and 25-75% percentiles, red bars indicate the mean. ns not significant, *$p < 0.05$, **$p < 0.01$, ***$p < 0.001$.

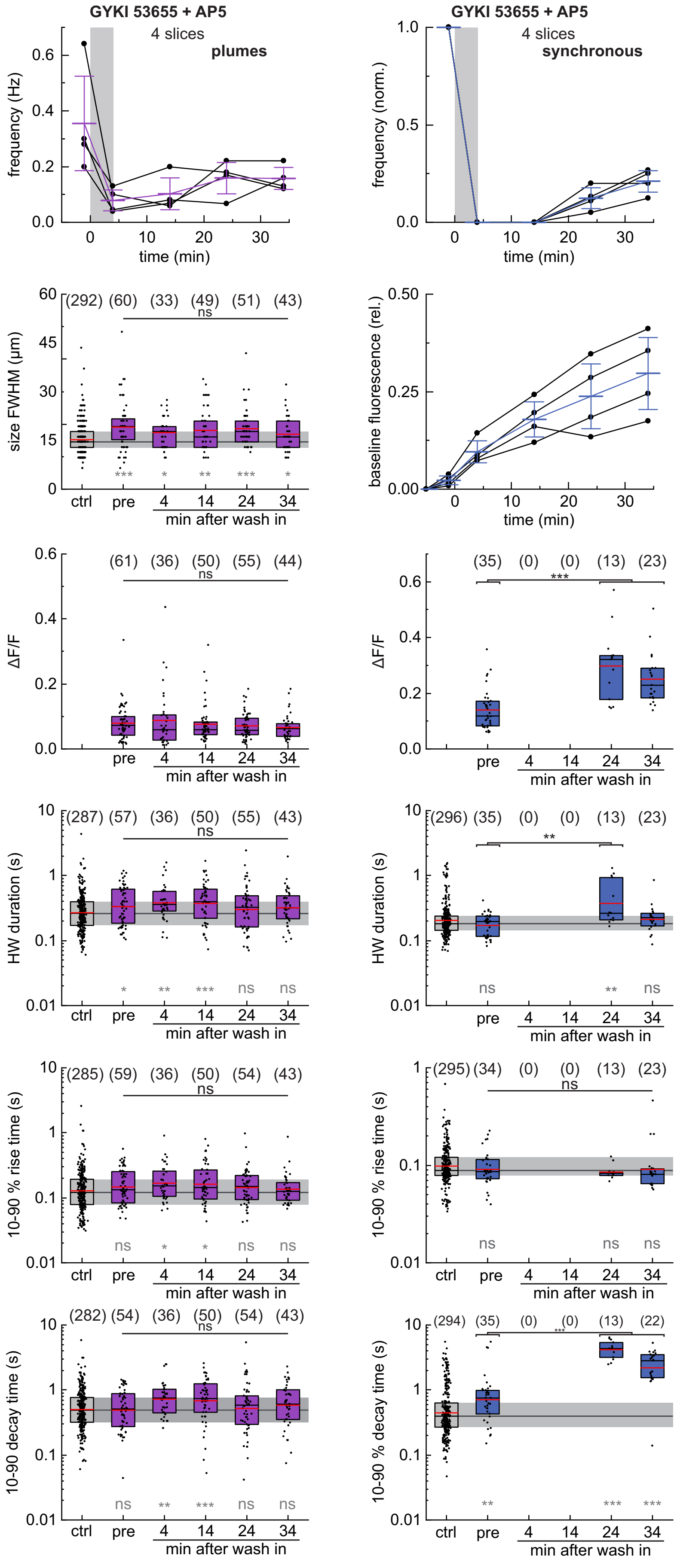
GYKI 53655 + AP5
4 slices
plumes
frequency (Hz)
time (min)
GYKI 53655 + AP5
4 slices
synchronous
frequency (norm.)
time (min)
(292) (60) (33) (49) (51) (43)
ns
size FWHM (µm)
*** * ** *** *
ctrl pre 4 14 24 34
min after wash in
baseline fluorescence (rel.)
time (min)
(61) (36) (50) (55) (44)
ns
ΔF/F
pre 4 14 24 34
min after wash in
(35) (0) (0) (13) (23)
***
ΔF/F
pre 4 14 24 34
min after wash in
(287) (57) (36) (50) (55) (43)
ns
HW duration (s)
* ** *** ns ns
ctrl pre 4 14 24 34
min after wash in
(296) (35) (0) (0) (13) (23)
**
HW duration (s)
ns ** ns
pre 4 14 24 34
min after wash in
(285) (59) (36) (50) (54) (43)
ns
10-90 % rise time (s)
ns * * ns ns
ctrl pre 4 14 24 34
min after wash in
(295) (34) (0) (0) (13) (23)
ns
10-90 % rise time (s)
ns ns ns
ctrl pre 4 14 24 34
min after wash in
(282) (54) (36) (50) (54) (43)
ns
10-90 decay time (s)
ns ** *** ns ns
ctrl pre 4 14 24 34
min after wash in
(294) (35) (0) (0) (13) (22)
***
10-90 % decay time (s)
** *** ***
ctrl pre 4 14 24 34
min after wash in

**Figure S18. GYKI 53655/AP5 effects on plumes and synchronous activity.** Coapplication of GYKI 53655 (50 µM) and D-AP5 (25 µM) caused a transient suppression of plumes and synchronous activity. Parameter summary of GYKI 53655/AP5 effects on glutamate plumes (left) and synchronous events (right) detected with SF-iGluSnFR(A184V) (4 slices). The investigation of synchronous glutamate events was partly limited by the imaging frame rate (20 fps). An example experiment and normalized plume frequencies are shown in **Fig. 5B**. Where possible, data were compared to the respective 40 min control recordings (ctrl column and grey bar; cf. **Fig. S8** and **Fig. S14**). Kruskal-Wallis tests were used to test for differences between the different time windows, followed by Dunn's test (black symbols). Additionally, Mann-Whitney *U* tests were used for pair-wise comparisons to control (grey symbols). Numbers in parentheses give the number of analyzed events. Bars with error bars show means ± s.d.. Boxes show medians and 25-75% percentiles, red bars indicate the mean. ns not significant, *$p$ <0.05, **$p$ <0.01, ***$p$ <0.001.

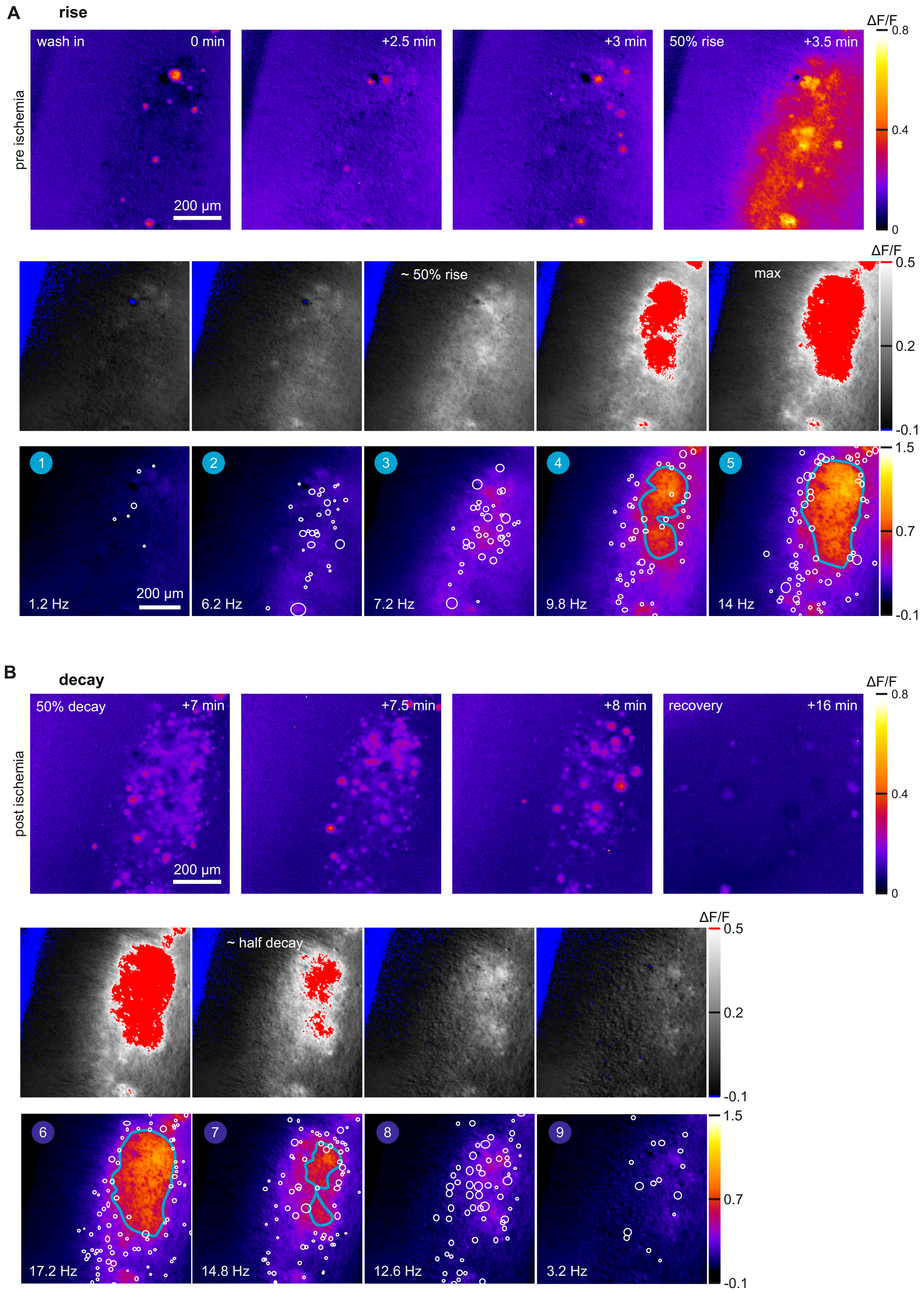


**Figure S19. Plumes and glutamate accumulation during chemical ischemia.** The figure shows the data underlying the depiction in **Fig. 6A-C**. **(A)** Analysis of ΔF/F signals at different timepoints during the rise in glutamate. Top: Maximum intensity projections over 500 frames to display plumes and Glu accumulation. Center: Minimum intensity projections of 100 frames to show glutamate accumulation without plumes. The look-up-table ‘Hi-Lo’ with thresholds at <0.1 (blue) and >0.5 (red) was used to separate non-responding regions and regions with high/maximal Glu accumulation. Bottom: Same minimal intensity projections displayed with the ‘fire’ look-up-table to indicate Glu accumulation. The blue lines depict regions, which have already reached their maximal signal change, white circles show plumes (full-width) occurring in the same time 100 frame window. **(B)** Analysis of different timepoints during the decay phase. Panels as shown in (A).

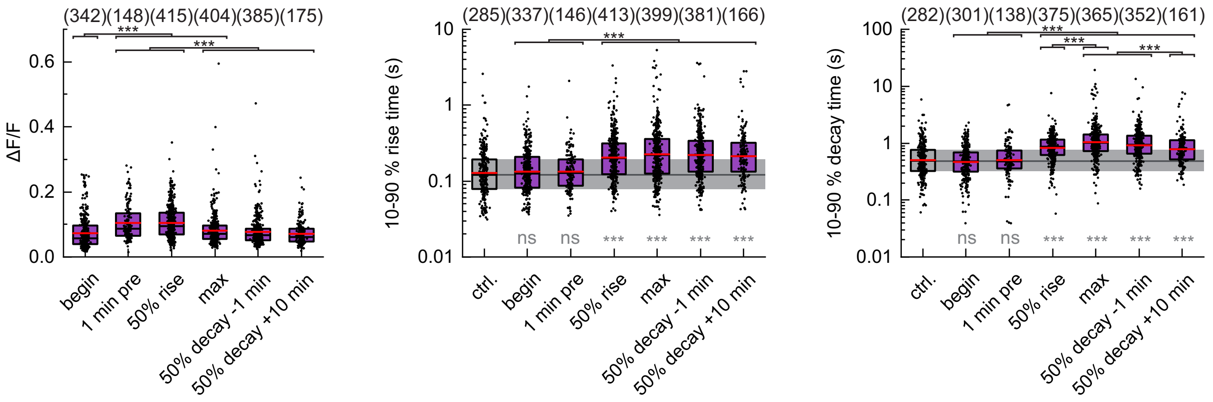


**Figure S20. Effects of chemical ischemia on plume parameters.** Plume properties were investigated at the indicated stages of the chemical ischemia from n = 11 slices. See **Fig. 6A** for an example experiment and **Fig. 6C** for additional parameters. ΔF/F and 10-90% rise and decay times of individual plumes are shown for the indicated time windows (5-50 s analysis windows). Kruskal-Wallis tests were used to test for differences between the different time windows, followed by Dunn's test (black symbols). Additionally, Mann-Whitney *U* tests were used for pair-wise comparisons to control (grey symbols). Numbers in parentheses give the number of analyzed plumes. Boxes show medians and 25-75% percentiles, red bars indicate the mean. ns not significant, **p* <0.05, ***p* <0.01, ****p* <0.001.

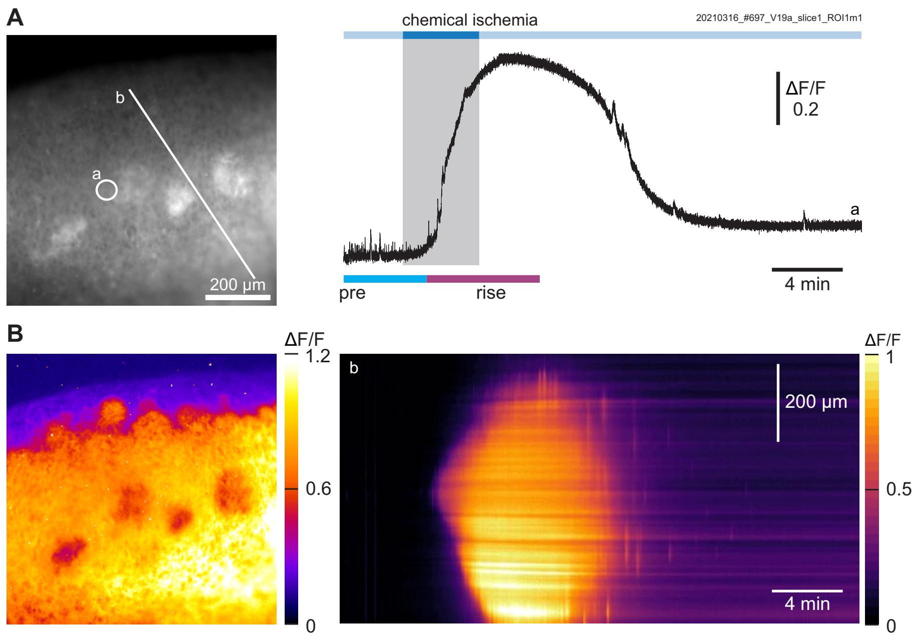


**Figure S21. Extracellular glutamate dynamics in an organotypic slice with culturing defects.** The slice was imaged at DIV18 and chemical ischemia was induced for 4 min. **(A)** SF-iGluSnFR(A184V) fluorescence before inducing ischemia (left) and ΔF/F trace (right) of region a. The pre-ischemic period and rise phase shown in **Fig. 6D** are indicated. Synchronous activity can be seen in the pre-ischemic phase but was not detected after ischemia. **(B)** Left: Glutamate accumulation due to chemical ischemia (ΔF/F at peak). The defect regions remain visible. Right: ΔF/F timecourse along the line region b.

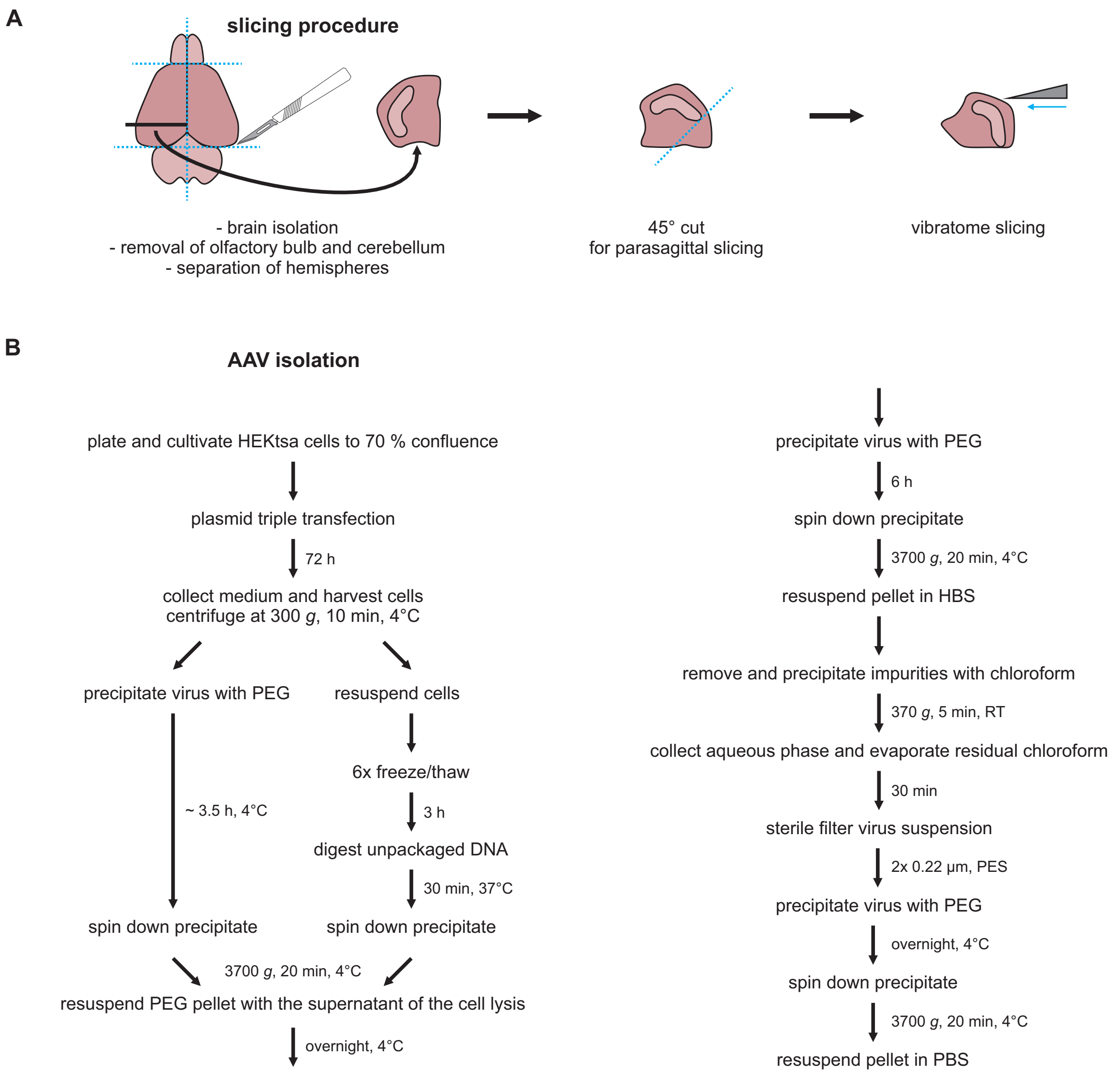


**Figure S22. Slicing procedure and isolation of rAAV particles. (A)** Slice preparation for organotypic slice cultures. The mouse brain is seen from top (left), before the indicated plane is shown from the caudal side. After slicing, the region containing the hippocampus and adjacent neocortex was cultured on membrane inserts. For details see **Methods**. **(B)** Scheme showing the steps used for the isolation of rAAV particles. For details see **Supplementary Methods**.

#### Supplementary Movie Information

**Movie S1. Extracellular glutamate accumulation upon inducing chemical ischemia in an organotypic brain slice.** The movie shows the SF-iGluSnFR(A184V) fluorescence intensity in grey (left) and the signal change ΔF/F normalized to the pre baseline in false colors (right). The movie corresponds to the slice shown in **Fig. 1A**. The movie speed is 25x real-time, the scalebar denotes 200 µm.

**Movie S2. Intracellular $Ca^{2+}$ loading upon inducing chemical ischemia in an organotypic brain slice.** The movie shows the GCaMP6f fluorescence intensity of the slice depicted in **Fig. 1E**. The movie speed is 25x real-time, the scalebar denotes 200 µm.

**Movie S3. Spontaneous plumes in a cortical brain region.** The movie shows the time after the last synchronous event depicted in **Fig. 2A** with the SF-iGluSnFR(A184V) fluorescence intensity in grey (left) and the signal change ΔF/F normalized with the baseline in **Fig. 2A** in false colors (right). The movie plays in real-time, the scalebar denotes 200 µm length.

**Movie S4. Example plume I.** The movie shows the plume depicted in **Fig. 3C** with the SF-iGluSnFR(A184V) fluorescence intensity in grey (left) and the signal change ΔF/F normalized with the baseline in **Fig. 2A** in false colors (right). The movie speed is 0.5x real-time, the scalebar denotes 25 µm.

**Movie S5. Example plume II.** The movie shows the slower plume, depicted in **Fig. 3E** with the SF-iGluSnFR(A184V) fluorescence intensity in grey (left) and the signal change ΔF/F normalized with the baseline in **Fig. 2A** in false colors (right). The movie speed is 0.5x real-time, the scalebar denotes 25 µm.

**Movie S6. SF-iGluSnFR(A184V) signals in response to TBOA application.** The movie shows the signal change ΔF/F normalized with the baseline at the beginning of the recording in false colors. The movie corresponds to the slice shown in **Fig. 5A** until 25 min. The movie speed is 25x real-time, the scalebar denotes 200 µm.

**Movie S7. Plumes during chemical ischemia in an organotypic brain slice.** The movie shows the SF-iGluSnFR(A184V) signal change ΔF/F normalized with the baseline at the beginning of the recording in false colors. The movie corresponds to the experiment shown in **Fig. 6A** until 16.7 min. The movie speed is 25x real-time, the scalebar denotes 200 µm.

**Movie S8. Chemical ischemia in a slice with defects.** The movie shows the slice from **Fig. 6D** with the SF-iGluSnFR(A184V) fluorescence intensity in grey (left) and the signal change ΔF/F normalized with the baseline at the beginning of the recording in false colors (right), see also **Fig. S21**. The movie speed is 25x real-time, the scalebar denotes 200 µm.

### Supplementary Methods

#### Preparation of rAAV particles

rAAVs were produced in-house (**Fig. S22B**) [Grieger et al., 2006]. For this HEK293Tsa cells with E1+ were grown in 151 cm$^2$ cell culture dishes with 20 ml DMEM (Sigma, D6429) with 7% fetal calf serum (Sigma, F9665) at 37°C and 5% $CO_2$. At 70% confluence plasmid triple transfections were performed. For this, serum-free DMEM (0.5 ml) was mixed with helper plasmid (12 µg), *trans*-plasmid (10 µg) and the respective *cis*-plasmid (6 µg) before adding polyethyleneimine (PEI) 25.000 (84 µg; Sigma, 408727; 1 mg/ml stock in water) per dish. AAV purification was performed similar to a published protocol [Eickelbeck *et al.*, 2019]. 72 h after transfection, cells were harvested with a cell scraper, separated from the medium by low-speed centrifugation (300 *g*, 4°C) and resuspended in 10 ml Tris-NaCl (150 mM NaCl, 50 mM Tris HCl; room temperature). Lysis was performed by six cycles of alternating freezing in dry ice/ethanol and thawing at 37°C, followed by 30 min incubation with DNase I (1.3 mg/ml; Roche, 11284932001) at 37°C. In parallel, virus particles were precipitated from the medium by adding 5 ml PEG-NaCl (Sigma 89510; 40% w/v in 2.5 M NaCl) and continuous shaking at 4°C (~3.5 h). Both, the lysed cells and the PEG-precipitated medium were then centrifuged for 20 min at 3700 *g* and 4°C. Subsequently, the supernatant of the lysed cells was used to resuspend the pellets obtained from the PEG precipitation. After resuspension and shaking at 4°C overnight, 2.5 ml PEG-NaCl was added for at least 6 hours to precipitate the AAV particles, the suspension was centrifuged for 20 min at 3700 *g* and 4°C, and the pellet was resuspended in 10 ml HBS (140 mM NaCl, 50 mM HEPES, pH 7.3). Chloroform (Fisher Scientific, 11408123) was added at 1:1 v/v ratio to remove impurities. After low-speed centrifugation at 370 *g* for 5 min at RT, the aqueous phase (without precipitate) was collected followed by 30 min of evaporation of residual chloroform. Then, the AAV suspension was sterile filtered twice (PES, 0.22 µm). Subsequently, 2.5 ml PEG-NaCl were added to precipitate the AAV particles overnight at 4°C. After 20 min centrifugation at 3700 *g* and 4°C, the pellet was resuspended in 80-100 µl PBS (Sigma, 806552) and stored at 4°C.

##### *Imaging of ATP levels (ATeam measurements)*

Experiments were carried out at the Heinrich Heine University Düsseldorf in strict accordance with the institutional guidelines as well as the European Community Council Directive (2010/63/EU). All experiments were communicated to and approved by the Animal Welfare Office at the Animal Care and Use Facility of the Heinrich Heine University Düsseldorf (Institutional act no. O50/05). In accordance with the recommendations of the European Commission [Close *et al.*, 1997], neonatal animals (7-9 days postnatal) were killed by rapid decapitation before preparation of brain slices.

Imaging of intracellular ATP changes in neurons was performed in layer II/III of the somatosensory cortex using the FRET-based nanosensor ATeam1.03$^{YEMK}$ ("ATeam") [Imamura *et al.*, 2009; Lerchundi *et al.*, 2020]. 0.5 µl of rAAV encoding for ATeam under the neuron-specific promoter human synapsin1 (hSyn1, AAV2/2) were applied on the cultured slices during the first 3 days *in vitro* [Lerchundi *et al.*, 2020]. The rAAV encoding for ATeam was generated at the Viral Vector Facility of the University/ETH Zürich (Switzerland). Slices were then kept in culture for at least 10 more days until experiments were performed. A confocal laser scanning microscope (Nikon Eclipse C1) equipped with a 40x Achroplan water immersion objective (Nikon, NA 0.8) was used to confirm neuron-specific expression of ATeam (**Fig. S1A**). ATeam was excited using a 488 nm argon ion laser and fluorescence emission was collected at 515 ± 15 nm. Maximum projections were constructed from *z*-stacks (step size 1.5 µm) using Fiji (ImageJ).

Prior to experiments, the glial scar on the transduced slices was removed as described [Lerchundi *et al.*, 2019]. Slices were placed in a bath chamber at room temperature and were superfused at 2-2.5 ml/min with Ringer's solution (130 mM NaCl, 2.5 mM KCl, 2 mM $CaCl_2$, 1 mM $MgCl_2$, 1.25 mM $NaH_2PO4$, 26 mM $NaHCO_3$ and 10 mM glucose, bubbled with 95% $O_2$ and 5% $CO_2$, resulting in a pH of 7.35). Fluorescence imaging was performed using an epifluorescence microscope (Nikon Eclipse FN-I) equipped with an 40x Achroplan water immersion objective (Nikon, NA 0.8). ATeam was excited at 434 nm using a Poly-V monochromator (Thermo Scientific/FEI) for 25-50 ms at a frequency of 0.5 Hz. Emitted light was split at 500 nm using a W-View GEMINI image splitter (Hamamatsu Photonics), band-pass filtered at 483/32 (eCFP, donor) and 542/27 (Venus, acceptor) and imaged with a sCMOS camera (Hamamatsu Photonics, Orca 4 LT Plus). Donor and acceptor fluorescence was collected from manually drawn regions of interest (ROIs) around neuronal cell bodies using the NIS-Elements software (Nikon). After background correction, the fluorescence ratio (Venus/eCFP, termed "ATeam ratio") was calculated for each ROI. Subsequent analysis was performed using Origin 2021 software (OriginLabs). Changes in the fluorescence ratio induced by chemical ischemia were normalized to the baseline and are expressed as % change in the ATeam fluorescence ratio (**Fig. S1B,C**).

#### SF-iGluSnFR(A148V) control experiments in HEK cells

The effect of azide and pH on SF-iGluSnFR(A148V) was tested 24-48 h after PEI-mediated transfection of HEK cells grown on plastic coverslips [see Pollok & Reiner, 2020]. Fluorescence imaging (**Fig. S2**) was performed on an inverted microscope (Leica DMi8). HEK cells were continuously perfused with extracellular solution (138 mM NaCl, 1.5 mM KCl, 1.2 mM $MgCl_2$, 2.5 mM $CaCl_2$ and 10 mM HEPES, pH 7.3 or pH 6.5) by a gravity-driven perfusion system. Epifluorescence excitation was provided by a collimator-coupled LED (Thorlabs, M470L3) with a 470/40 nm excitation filter (Chroma, ET 470/40x) and a 495 nm dichroic mirror (Chroma, T 495 LPXR). A 525/50 nm emission filter (Chroma, ET 525/50m) was used and continuous imaging with 2 fps was performed with an EMCCD camera (Photometrics, Evolve 512delta) at 16-bit 512 x 512 pixels or 256 x 256 pixels using MicroManager2.0 [Edelstein *et al.*, 2014]. The obtained images were processed using ImageJ 1.53t. For dose-response analysis of SF-iGluSnFR(A184V), four individual experiments at each, pH 7.3 or pH 6.5, were performed. From each recording, the fluorescence intensities from six different cells were exported and analyzed in Excel (Microsoft). $F_0$ was determined individually for each application to account for signal run up or bleaching, by averaging 10 frames (5 s) before glutamate application (0.2-1000 µM glutamate in extracellular solution at pH 7.3 or pH 6.5) and 10 frames after the signal had stabilized after washout. Relative fluorescence intensity changes ($\Delta F/F$) were calculated after averaging 10 frames at the plateau of each glutamate application. The obtained $\Delta F/F$ values were normalized to the signal change obtained with 1000 µM glutamate. $EC_{50}$ values were determined by fitting the dose-response curves with the Hill equation using ProFit (Quantumsoft) with the Levenberg-Marquardt algorithm and fixed Hill coefficient ($n = 1$) and $\Delta F/F_{min} = 0$. The obtained $EC_{50}$ values (**Fig. S2**) were close to the reported value for SF-iGluSnFR(A184V) (reported $EC_{50}$= 2.1 µM) [Marvin et al., 2018]. The relative signal changes upon reducing to pH 6.5 were calculated with $F_0$ being the signal at pH 7.3 in the absence and presence of saturating glutamate (100 µM). $NaN_3$ (5 mM) in extracellular solution was applied for 5 min.

##### *Details on data analysis, presentation, and statistics*

*Figure 1 and corresponding Supplementary Figures.* To obtain ΔF/F maps of glutamate accumulations (**Fig. 1A**, **Fig. 6A,** and **Fig. S21B**) and $Ca^{2+}$ accumulations during chemical ischemia, average intensity projections of 200 frames were generated at the peak of accumulation (F). The mean resulting from average intensity projections of 200 frames before and 200 frames after chemical ischemia was used for correction ($F_0$). For normalization of full-length traces from selected regions (**Fig. 1A,E**, **Fig. 6A**, **Fig. 7A**, **Fig. S3**, **Fig. S8A**, **Fig. S9A Fig. S21B**) the intensity $F_0$ from the beginning of the recordings was used (average intensity projections of 10-100 frames). To create pseudo-color time traces along line regions (**Fig. 1A** and **Fig. S21B**), intensity values of all pixels along a line region of interest were obtained from ΔF/F stacks, imported to ClampFit 11.1 (Molecular Devices) for manual baseline adjustments (linear segments), and visualized in Origin Pro 2021 (OriginLabs) using an inverted 'inferno' color scale.

Responding regions, i.e. those that showed signal increase, were manually defined (single polygon) and used to calculate peak ΔF/F, 50 % rise time (time from wash-in until 50 % max ΔF/F was reached), halfwidth duration and 50 % decay time (time from wash-out until 50 % decay) using ClampFit 11.1. The same regions responding to ischemia were also used to analyze synchronous activity in these slices (**Fig. 1D,H**, **Fig. S7** and **Fig. S9D,E**), however, using selected sub-stacks, which were normalized to $F_0$ images from these sub-stacks (see **Main Methods**). For frequency determinations, stack lengths were adapted to have at least 10 events or having a maximal length of 300 s. High $Ca^{2+}$ event frequencies (pre- and post-ischemic 'high', **Fig. 1H** and **Fig. S9D**) were analyzed in 30 s time windows around maximal activity.

To obtain ΔF/F maps of synchronous activity (**Fig. 2A, Fig. S3B** and **Fig. S10A**) maps, average intensity projections (AIPs) were obtained from peak ΔF/F frames of all synchronous events within 500 frames. Here, normalization had been performed with F average intensity projections from the beginning of the recording ($F_0$, 200 frames). For comparing ΔF/F maps of ischemia and glutamate application (**Fig. S3B**), 200 frames during the fluorescence peak were averaged and normalized with the same $F_0$ image. Ratio maps (**Fig. S3C**) were calculated after binning to 32 x 32 pixels by dividing the ΔF/F map of glutamate accumulation during ischemia with the ΔF/F map of synchronous activity or exogenous glutamate application, respectively. Then, all pixel values were measured and imported to Origin Pro 2021. The three parameters were plotted pairwise with red line indicating a correlation coefficient of 1 (**Fig. S3D**).

To quantify the signal loss of synchronous activity before the glutamate rise (**Fig. S7A**), all events in the 150 s preceding the loss of synchronous activity were analyzed. The ΔF/F values of these events were normalized using the average ΔF/F of synchronous activity that was determined in each slice at the beginning of the recording. The resulting normalized $\Delta F/F_{norm}$ values fluctuated but eventually declined. For better comparison, the events were binned relative to the first event after which events permanently showed a $\Delta F/F_{norm}$ <0.5. Two 50 s bins were created before this event and two 25 s bins after this event, before spontaneous activity was lost in all slices.

In control slices, in which synchronous glutamate and $Ca^{2+}$ activity and plumes were imaged over 50 min with no pharmacological applications (**Fig. S8**), all events were analyzed in 25-300 s time windows at the beginning of the recording and after 5, 10, 20, 30 and 40 min as described above (also shown in **Fig. 4**, **Fig. 5**, **Fig. 6C** and corresponding Supplementary Figures).

*Figures 2 & 3 corresponding Supplementary Figures.* Maximum intensity projections (MIP) showing plumes (**Fig. 2A** and **Fig. S10A**) were calculated from 500 frames as described in the **Main Methods**. Synchronous activity maps were generated as average intensity projections of peak ΔF/F frames of all synchronous events within 500 frames (see above). All plumes and synchronous activity occurring in these 500 frames were plotted as time traces (**Fig. 2A** and **Fig. S10A**). For plume detection see **Main Methods**. Frequencies of plumes and synchronous activity were determined for all slices subjected to chemical ischemia at the beginning of the recording as described above. We analyzed plume HW duration, plume intensity ΔF/F, and the ΔF/F ratio of plume intensity compared to synchronous activity in the same region of interest (**Fig. 2B**). Dashed lines reflect the plume detection limits based on intensity ($\Delta F/F \geq 2\%$,) and HW duration ($\geq 50$ ms). For comparing plume parameters, only plumes for which all parameters could be characterized were considered ($n = 296$ plumes, **Fig. 2B** and **Fig. 3B**). Pearson and Spearman correlations were analyzed and plotted with Origin Pro 2021 (**Fig. S12A**).

To show spatial and temporal characteristics of selected plumes, 8 single frames from the binned ΔF/F stacks were selected (**Fig. 3C-F)**. The corresponding time course and the spatial profile are shown for the depicted circled region (full-width plume) and box, respectively. The spatial profiles are depicted after another 2x2 binning. **Fig. S13B** shows unbinned ΔF/F data and binned raw intensity (F) data.

*Figures 4, Figure 5 and corresponding SI Figures.* For pharmacological characterizations, the analysis of synchronous glutamate activity, $Ca^{2+}$ activity, and identification and analysis of plumes was performed 1 min before applying blocker and at the indicated times after application by extracting the corresponding stacks. Frequencies were determined as described above. For low event frequencies the analyzed time windows were extended from 500 frames to 6000 frames. For high plume frequencies during TBOA application they were shortened to 100 frames.

*Figure 6 and corresponding SI Figures.* For plume analysis at the indicated time point of chemical ischemia experiments (beginning of the recording, 1 min before wash-in, at 50 % signal rise, maximum signal, 1 min before 50 % decay and 4 and 10 min after 50% decay) 500-2000 frames were extracted. The standard plume analysis (**Fig. 6C**) was based on 100-1000 frames.

To visualize glutamate accumulation without plumes and synchronous activity (**Fig. 6B** and **Fig. S19A,B**, Middle and Bottom rows), 100 frame ΔF/F minimum intensity projections (normalized with $F_0$ from the beginning of the recording) were calculated and displayed with the ImageJ LUTs "Fire" and "HiLo". To display glutamate accumulation and plumes, 500 frame ΔF/F maximum intensity projections were used (**Fig. S19A,B**, Top rows). For **Fig. 6D**, 200 frame ΔF/F maximum intensity projections (normalized with $F_0$ from the beginning of the recording) were generated to show plumes before inducing chemical ischemia (pre ischemia) and to display the spread of glutamate accumulation from defective regions during the rise phase.

*Figure 7*. Plume frequencies were determined as described in **Main Methods** at the beginning of the recording, 1 min before applying iGluR inhibitors, at the beginning of ischemia wash-in, 10 and 20 min after ischemia with inhibitors and 1 min before wash-in, at 50 % signal rise, max signal, 1 min before 50 % decay and 4 and 10 min after 50% decay of the inhibitor-free ischemia.

*Supplementary Movies*. To create **Supplementary Movies 1,2** and **6-8**, data were first reduced in ImageJ. In each stack (16-bit raw fluorescence intensity movies, 32-bit ΔF/F movies), 5 consecutive frames were averaged with the "Grouped Z Project" tool. For further reduction, the movies were converted to 8-bit with the lowest and highest values in the stacks as limits. For **Supplementary Movies 3-5** no averaging was performed. The speed was adapted as indicated in **Supplementary Movie Information** during export at 10-100 fps to AVI files with JPEG image compression. Compressed movies were imported into VideoStudio X10 (Corel) and scalebars and application markers were added. The final movies were exported in MPEG-4 format.

##### *Computational modeling of transient chemical ischemia*

The simulations are based on a computational model of a neuron and an astrocyte in a finite extracellular space that we had previously developed for a similar experimental setup [Kalia *et al.*, 2021; Engels *et al.*, 2021]. There we used it to explain changes in sodium, potassium and chloride concentrations as well as cell volume. The model also included $Ca^{2+}$ and glutamate, but those components were not the main focus. We adapted and used this computational model to further understand the observations reported here. The model with all compartments and active and passive transport processes is shown in **Fig. S4.**

In the following, we motivate the changes to our model: Previously, we used a finite, closed extracellular space, while in the experiments there is a continuous flow of solution, which refreshes the extracellular space surrounding the neuron and astrocyte (bath exchange). To account for this exchange, we included diffusion from the extracellular space to an external bath with constant concentrations. We connected the extracellular space to the external bath by diffusive coupling of all ions and glutamate. To model differences in exchange rate that might be seen in different regions in the slice (thicker or more dense regions *vs.* thinner or less dense regions) we choose various values for the strength of this diffusion. Further, the ATP measurements (**Fig. S1**) gave estimates of the drop in energy that was previously unknown. Motivated by the diffusion of the bath exchange, we adopted the modelling of available oxygen as in [Wei *et al.*, 2014], as a proxy for the available ATP as this is directly related. This model has been used in similar contexts. Specifically, the strength of the sodium potassium ATPase (NKA) pumps indirectly depends on the available extracellular oxygen, and the pumps indirectly consume oxygen, which is replenished through diffusion with the external bath. To simulate the response to chemical ischemia, we lowered the oxygen level from 100% to 10% of baseline in the external bath for four minutes, and then restored it back to 100%. The remaining 10% oxygen could account for reserve forms of energy such as lactate.

Next, there is considerable spontaneous network activity (see e.g. **Fig. 1**), and we thus added an input (‘applied current’, **Fig. S5**) to the neuron every two minutes during the entire simulation (before, during, and after ischemia). This input also gives information about neuronal excitability. Further, the GCaMP6 signals representing neuronal $Ca^{2+}$ (see e.g. **Fig. 1**) suggest more $Ca^{2+}$ inflow than our model showed previously. We thus increased the voltage-gated calcium permeability (‘gated $Ca^{2+}$’) 50-fold. Finally, for glutamate we now implemented the full nonlinear expression for the EAAT current from [Breslin *et al.*, 2018] (instead of its linearization), including two limiting terms as in [Flanagan *et al.*, 2018], which limit EAAT-mediated Glu uptake if there is too little glutamate in the extracellular space, or, if loading into vesicles is saturated.

For the simulations, we used the model by Kalia [Kalia *et al*., 2021] and all model components i.e. those for $Na^+$, $K^+$, and $Cl^-$, can be found there. Here we only describe the changes we made to the model, in particular for $Ca^{2+}$ and glutamate, and how we included diffusion.

For the neuronal or astrocytic compartment (x = n, a), we have the membrane potential $V_x$. The expression for the EAAT current follows the model from [Breslin *et al*., 2018]:

$$J_{EAAT} = P_{EAAT,x}\left(\exp\left(-\beta_{EAAT}\left(V_x - E_{EAAT_x}\right)\right) - 1\right),$$

where $\beta_{EAAT} = 0.0292\, mV^{-1}$ , $P_{EAAT,a} = 2 * 10^{-7} \mu m^3 (ms)^{-1}$ and $P_{EAAT,n} = P_{EAAT,a}/9$ to preserve a 1:9 ratio. The reversal potential is given by:

$$E_{EAAT,x} = \frac{RT}{2F} log\left(\frac{[Na^+]_e^3[K^+]_x[H^+]_e[Glu]_e}{[Na^+]_x^3[K^+]_e[H^+]_x[Glu]_x}\right).$$

Free intracellular glutamate $[Glu]_n$ is transported into a depot of vesicles with a time constant $\tau_{rec}$ until this process is saturated.

The neuronal and astrocytic NKA pumps consume energy modeled directly with oxygen as in [Wei *et al*., 2014]. The available extracellular oxygen $[O_2]$ changes as:

$$\frac{d[O_2]}{dt} = -\frac{\alpha_{O_2}}{F}\left(J_{NKA,n} + J_{NKA,a}\right) + \varepsilon_{O_2}\left([O_{2,bath}](t) - [O_2]\right)$$

where $\alpha_{O_2} = 0.1656\, g/mol$, $\varepsilon_{O_2} = 7.9688 * 10^{-6}\, ms^{-1}$. The baseline level we set as $[O_{2,bath}] = 58\, \mathrm{g/mol}$ and to model ischemia we set $[O_{2,bath}] = 5.8\, \mathrm{g/mol}$ from t=10 to t=14 minutes. The normal NKA pump current $I_{NKA,x}$ is modulated by oxygen as:

$$J_{NKA,x} = \frac{1}{1+exp\left((20-[O_2])/3\right)} I_{NKA,x}.$$

For completeness, we mention the active and passive calcium currents as they were not described in [Kalia *et al*., 2021]. The calcium leak currents are given by

$$J_{Ca,L,x} = \frac{4P_{Ca,L,x}F^2V}{RT}\left(\frac{[Ca]_x - [Ca]_e exp(-2FV_x/RT)}{1-exp(-2FV_x/RT)}\right)$$

and the gated neuronal calcium current is given by

$$J_{Ca,g,n} = \frac{4P_{Ca,g,n}F^2V_n}{RT}\left(\frac{[Ca]_n - [Ca]_e exp(-2FV_n/RT)}{1-exp(-2FV_n/RT)}\right).$$

For all extracellular ions and glutamate we add diffusion to the bath as:

$$\frac{dN_x}{dt} = \sum I_{ion,x} + B(N_B - N_x),$$

where we vary the diffusion strength as $B$ = 2*10$^{-4}$, 1*10$^{-3}$, 5*10$^{-3}$ ms$^{-1}$.

As in [Kalia *et al.*, 2021], we set the volume of the neuron and astrocyte to 2.0 and 1.7 mm$^3$, respectively, and then choose the volume of the extracellular space so it initially had 20% of the total volume. The synaptic cleft and neuron and astrocyte compartments where glutamate and calcium reside have a fixed volume of 1 µm$^3$, which is much smaller and thus negligible for the total volume.

The simulation code can be found at
https://gitlab.utwente.nl/m7686441/focalglutamate_modelsimulations

*General properties.* We investigated key components of the model by injecting currents every two minutes before we additionally induced chemical ischemia for 4 min (**Fig. S5**). The behavior was similar to the results obtained with the previous model [Kalia *et al.*, 2021]. The neuron was excitable under baseline conditions (see membrane potential) and maintained a normal resting potential and low $Na^+$, $Cl^-$ and $Ca^{2+}$ concentrations with significant NKA activity. Inducing chemical ischemia for 4 min resulted in deterioration of all neuronal and astrocytic ion gradients within a few minutes along with strong depolarization. Neuronal excitability was lost as well as NKA activity. After restoring oxygen supply, most components returned towards baseline levels and neuronal excitability was regained. The model also shows autonomous, high-frequency neuronal activity during the onset of ischemia, namely when neuronal depolarization has crossed the firing threshold but not reached the depolarization block, yet (see also [Kalia *et al.*, 2021]).

*Effect of bath coupling.* When we varied the diffusion strength to the bath from weak to strong ($B$ = 2*10$^{-4}$, 1*10$^{-3}$, 5*10$^{-3}$ ms$^{-1}$) we found that oxygen depletion in the extracellular space, i.e. the extent to which chemical ischemia was induced, did not vary (**Fig. S6A**). Also, strong neuronal $Ca^{2+}$ loading was seen in all conditions (**Fig. S6B**). However, the timecourse and extent of extracellular glutamate accumulation varied from ~75 µM (maximum at low coupling strength) to ~3 µM (maximum at high coupling strength) (**Fig. S6C**). The other model components showed overall similar behavior to the situation seen with weak bath coupling (**Fig. S5**), but partial recovery was somewhat faster with stronger bath coupling.

#### Supplementary References

Breslin K, Wade JJ, Wong-Lin K, Harkin J, Flanagan B, Van Zalinge H, Hall S, Walker M, Verkhratsky A, & McDaid L. (2018). Potassium and sodium microdomains in thin astroglial processes: A computational model study. *PLOS Computational Biology* **14**: e1006151.

Close B, Banister K, Baumans V, Bernoth E-M, Bromage N, Bunyan J, Erhardt W, Flecknell P, Gregory N, Hackbarth H, Morton D, & Warwick C. (1997). Recommendations for euthanasia of experimental animals: Part 2. *Laboratory Animals* **31**: 1–32.

Edelstein AD, Tsuchida MA, Amodaj N, Pinkard H, Vale RD, & Stuurman N. (2014). Advanced methods of microscope control using µManager software. *Journal of Biological Methods* **1**: e10.

Eickelbeck D, Karapinar R, Jack A, Suess ST, Barzan R, Azimi Z, Surdin T, Grömmke M, Mark MD, Gerwert K, Jancke D, Wahle P, Spoida K, & Herlitze S. (2019). CaMello-XR enables visualization and optogenetic control of Gq/11 signals and receptor trafficking in GPCR-specific domains. *Communications Biology* **2**: 60.

Engels M, Kalia M, Rahmati S, Petersilie L, Kovermann P, van Putten MJAM, Rose CR, Meijer HGE, Gensch T, & Fahlke C. (2021). Glial chloride homeostasis under transient ischemic stress. *Frontiers in Cellular Neuroscience* **15**: 735300.

Flanagan B, McDaid L, Wade J, Wong-Lin K, & Harkin J. (2018). A computational study of astrocytic glutamate influence on post-synaptic neuronal excitability. *PLOS Computational Biology* **14**: e1006040.

Grieger JC, Choi VW, & Samulski RJ. (2006). Production and characterization of adeno-associated viral vectors. *Nature Protocols* **1**: 1412–1428.

Imamura H, Huynh Nhat KP, Togawa H, Saito K, Iino R, Kato-Yamada Y, Nagai T, & Noji H. (2009). Visualization of ATP levels inside single living cells with fluorescence resonance energy transfer-based genetically encoded indicators. *Proceedings of the National Academy of Sciences USA* **106**: 15651–15656.

Kalia M, Meijer HGE, van Gils SA, van Putten MJAM, & Rose CR. (2021). Ion dynamics at the energy-deprived tripartite synapse. *PLOS Computational Biology* **17**: e1009019.

Lerchundi R, Huang N, & Rose CR. (2020). Quantitative imaging of changes in astrocytic and neuronal adenosine triphosphate using two different variants of ATeam. *Frontiers in Cellular Neuroscience* **14**: 80.

Lerchundi R, Kafitz KW, Färfers M, Beyer F, Huang N, & Rose CR. (2019). Imaging of intracellular ATP in organotypic tissue slices of the mouse brain using the FRET-based sensor ATeam1.03$^{YEMK}$. *Journal of Visualized Experiments* **154**: e60294.

Pollok S, & Reiner A. (2020). Subunit-selective iGluR antagonists can potentiate heteromeric receptor responses by blocking desensitization. *Proceedings of the National Academy of Sciences USA* **117**: 25851–25858.

Wei Y, Ullah G, Ingram J, & Schiff SJ. (2014). Oxygen and seizure dynamics: II. Computational modeling. *Journal of Neurophysiology* **112**: 213–223.
